# Sex-stratified adipose-liver circuits in human MASLD

**DOI:** 10.64898/2026.09.24.754015

**Authors:** Mijra Koning, Iman Man Hu, Artemiy Kovynev, Raymond Landgraaf, Amir H. Alizadeh Bahmani, Daniël Baars, Danijela Vojnović Milutinović, Nataša Veličković, Rutger Franken, Yair Acherman, Nordin M. Hanssen, Hilde Herrema, Aleksander Krag, Fredrik Bäckhed, Jacques J.G. Bergman, Max Nieuwdorp, Koen H.M. Prange, Abraham S. Meijnikman

## Abstract

Metabolic dysfunction-associated steatotic liver disease (MASLD) is a common, clinically heterogeneous condition associated with liver-related and cardiometabolic morbidity. Sex differences in MASLD prevalence and progression are well recognized, but how they extend across the liver and visceral adipose tissue (VAT), a major source of lipid and endocrine signals to the liver, remains unclear. Here, we profiled paired liver and VAT biopsies from women and men with and without MASLD, generating single-nucleus datasets of 34 liver samples and 40 VAT samples, including 32 patient-matched pairs. In liver, women with MASLD showed a higher Kupffer-cell triglyceride-catabolism score (1.30-SD female-male difference; FDR=0.0022), accompanied by directionally consistent hepatocyte and endothelial programs. In VAT, women with MASLD showed higher adipocyte lipid and oxidative metabolism (0.87-1.15 SD; FDR<0.05 across seven pathways) together with macrophage immune activation, whereas men with MASLD showed stronger stromal communication routes. Across matched patients, gene modules linked VAT adipocyte and macrophage states to liver hepatocyte stress and lipid-receipt states. Bulk RNA-seq module scoring and paired liver-VAT analyses in 293 patients supported hepatic stress in both sexes, lower adipocyte signaling in women with increasing steatosis, and cross-tissue adipocyte-hepatocyte coupling (r=0.30; FDR=2.1e-6). Together, these findings identify distinct sex-associated transcriptional states in liver and VAT in MASLD and show that these states are coordinated across both tissues within the same individuals.

---

Metabolic dysfunction-associated steatotic liver disease (MASLD) is defined by hepatic steatosis in the presence of cardiometabolic risk and without excessive alcohol consumption or another dominant cause of steatosis.^1^ MASLD can progress from simple steatosis to metabolic dysfunction-associated steatohepatitis (MASH), fibrosis, cirrhosis and hepatocellular carcinoma (HCC). As the most common liver disease worldwide, MASLD is now a major focus of therapeutic development.^2–5^ However, treatment development remains complicated by disease heterogeneity, including sex differences.

Sex differences in MASLD have been studied at clinical, metabolic and experimental levels. Men have a higher overall prevalence of MASLD and MASH and a higher reported risk of liver-related complications and HCC, whereas this pattern changes with age and reproductive status, with post-menopausal women showing increased risk of MASH and liver-related complications compared with men of similar age.^6,7^ Several mechanisms may contribute to this dimorphism. Men generally have more visceral adipose tissue (VAT), a depot linked to lipolysis, insulin resistance and ectopic lipid delivery to the liver.^8–10^ Sex hormone signaling, particularly estrogen receptor signaling, shapes hepatic lipid metabolism, inflammation and regeneration, while estrogen action in other metabolic tissues also contributes to systemic insulin sensitivity.^11–13^ In parallel, the hepatic immune compartment differs between women and men, and experimental models point to sex-dependent immune protection or susceptibility in steatohepatitis.^14–16^

Despite this literature, the cellular basis of sex differences in human MASLD remains incompletely resolved. Much of the mechanistic evidence comes from murine models, often with limited female representation or strain-specific context, and its relevance to human liver and adipose tissue biology is still being defined.^12,17^ Human bulk-transcriptomic work has shown that sex is a major determinant of hepatic molecular signatures in MASH,^18^ but bulk profiles cannot localize sex-associated programs to specific parenchymal, immune, stromal or vascular compartments. This is particularly important for MASLD, where liver disease is closely linked to VAT metabolism, adipose inflammation and inter-organ crosstalk.

Here, we analyzed single-nucleus RNA sequencing (snRNA-seq) data from liver and adipose tissue biopsies collected in the BARIA cohort^19^ from women and men with and without MASLD. We aimed to explore sex-specific differences in the cellular and molecular organization of human MASLD across liver and adipose tissue.

## Results

### Paired liver and visceral-fat single-nucleus datasets

To study sex-stratified MASLD biology across metabolic tissues, we generated snRNA-seq datasets from liver and visceral adipose tissue (VAT) biopsies (Fig. 1a). Clinical characteristics of all 42 patients contributing at least one tissue are summarized in Table 1. We included 21 women and 21 men with similar age, BMI, diabetes prevalence and MASLD-category distributions; HDL cholesterol was higher in women, whereas gamma-glutamyl transferase, aspartate aminotransferase and alanine aminotransferase were higher in men, and concomitant medication use is summarized in Supplementary Table 1. After pre- and post-sequencing quality control, 34 of 40 liver libraries and all 40 VAT libraries were retained, representing 146,534 liver nuclei and 374,312 VAT nuclei; 32 patients had both tissues available for paired cross-tissue analyses (Fig. 1b and Supplementary Table 2).

**Figure 1.**
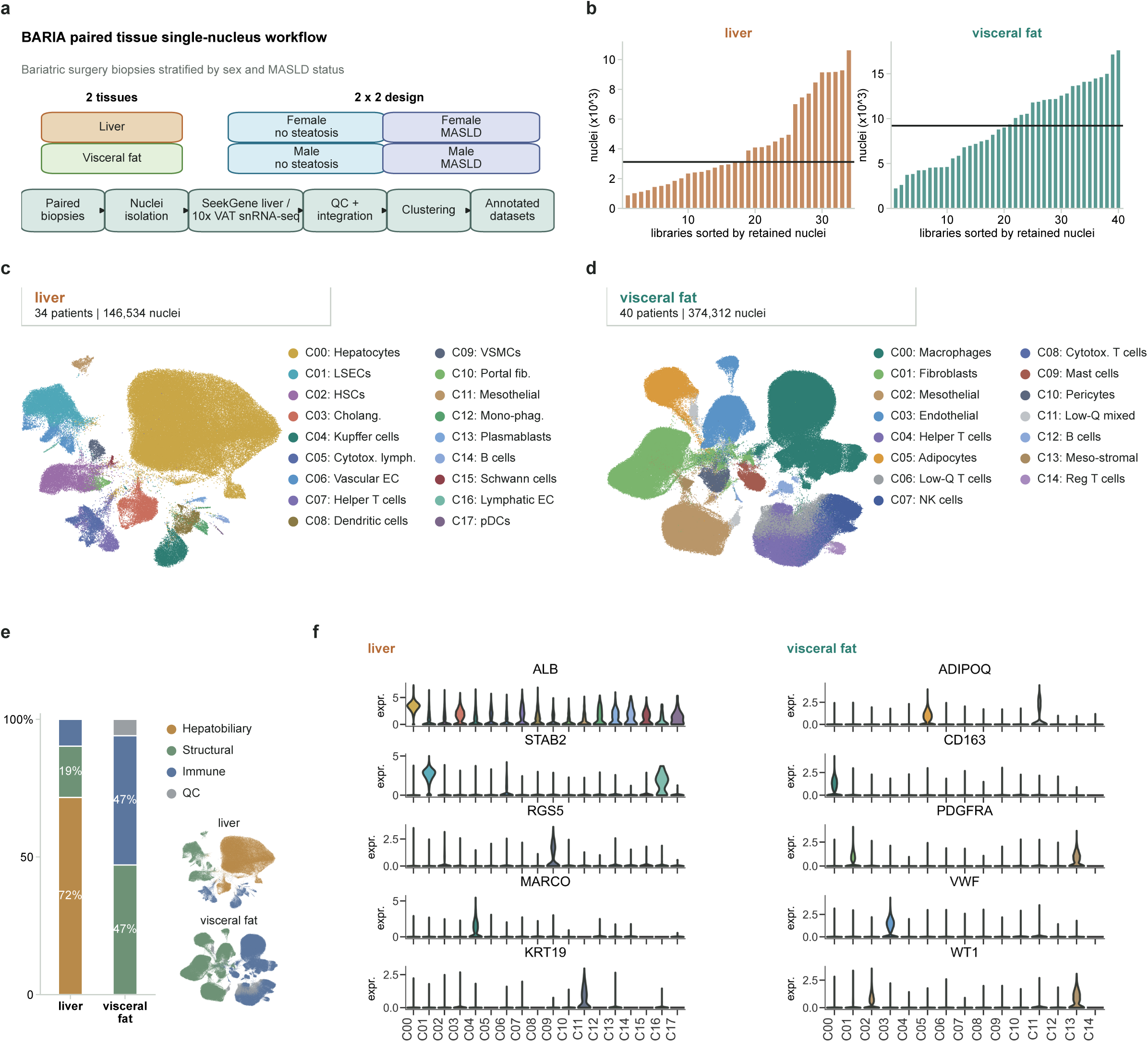
Paired liver and visceral-fat single-nucleus datasets. **a,** Study and single-nucleus workflow for paired liver and visceral-fat biopsies. **b,** Retained nuclei per patient library after QC, shown separately for liver and visceral fat; horizontal lines indicate tissue-specific medians. **c,d,** UMAPs of the final liver and visceral-fat datasets at the archetype annotation layer. Header boxes show the number of patient libraries and retained nuclei. **e,** Broad supercompartment composition of the liver and visceral-fat datasets, with inset UMAPs showing the location of the same supercompartment labels. **f,** Focused marker-gene violin plots validating major liver and visceral-fat cell populations. fib., fibroblasts; phag., phagocytes.

**Table 1.** Clinical characteristics of patients contributing to the single-nucleus study. The table describes all 42 patients who contributed liver, visceral adipose tissue, or both tissues to the final single-nucleus datasets.

| Characteristic | Men (n=21) | Women (n=21) | P value |
| --- | --- | --- | --- |
| **Demographics** |  |  |  |
| Age, years | 50 [43, 56] | 45 [40, 54] | 0.371 |
| BMI, kg/m<sup>2</sup> | 37.00 [35.00, 40.00] | 37.00 [34.00, 39.00] | 0.463 |
| Diabetes, n (%) | 4 (19.0%) | 5 (23.8%) | 1.000 |
| No hypertension, n (%) | 13 (61.9%) | 17 (81.0%) | 0.306 |
| GLP-1 receptor agonist use, n (%) | 1 (4.8%) | 0 (0.0%) | 1.000 |
| MASLD status, n (%) |  |  | 0.562 |
| No steatosis | 10 (47.6%) | 9 (42.9%) |  |
| MASLD | 6 (28.6%) | 9 (42.9%) |  |
| MASH | 5 (23.8%) | 3 (14.3%) |  |
| **Clinical laboratory values** |  |  |  |
| Alkaline phosphatase, U/L | 86 [75, 96] | 79 [58, 97] | 0.358 |
| Gamma-glutamyl transferase, U/L | 35 [28, 54] | 25 [17, 41] | 0.021 |
| Aspartate aminotransferase, U/L | 29 [22, 33] | 22 [18, 28] | 0.041 |
| Alanine aminotransferase, U/L | 40 [28, 54] | 25 [19, 42] | 0.041 |
| Total cholesterol, mmol/L | 4.84 (0.93) | 5.10 (0.96) | 0.373 |
| LDL cholesterol, mmol/L | 3.16 (0.83) | 3.35 (0.77) | 0.441 |
| HDL cholesterol, mmol/L | 1.00 [0.90, 1.10] | 1.20 [1.00, 1.30] | 0.045 |
| Triglycerides, mmol/L | 1.64 [1.19, 3.20] | 1.63 [1.18, 2.76] | 0.629 |
| Glucose, mmol/L | 5.70 [5.40, 6.20] | 6.00 [5.60, 6.60] | 0.332 |
| HbA1c, % | 5.6 [5.3, 6.1] | 5.5 [5.4, 6.2] | 0.900 |
| C-reactive protein, mg/L | 3.40 [2.08, 7.12] | 4.70 [2.20, 7.30] | 0.382 |
| **Pathology** |  |  |  |
| Steatosis grade, n (%) |  |  | 0.922 |
| <5% | 10 (47.6%) | 9 (42.9%) |  |
| 5-33% | 3 (14.3%) | 4 (19.0%) |  |
| 33-66% | 5 (23.8%) | 6 (28.6%) |  |
| >66% | 3 (14.3%) | 2 (9.5%) |  |
| Fibrosis stage, n (%) |  |  | 0.785 |
| 0 | 3 (15.0%) | 3 (14.3%) |  |
| 1 | 6 (30.0%) | 5 (23.8%) |  |
| 2 | 10 (50.0%) | 10 (47.6%) |  |
| 3 | 1 (5.0%) | 3 (14.3%) |  |
| Ballooning grade, n (%) |  |  | 0.717 |
| 0 | 15 (71.4%) | 17 (81.0%) |  |
| 1 | 6 (28.6%) | 4 (19.0%) |  |
| 2 | 0 (0.0%) | 0 (0.0%) |  |
| Lobular inflammation grade, n (%) |  |  | 0.418 |
| 0 | 4 (21.1%) | 1 (5.9%) |  |
| 1 | 6 (31.6%) | 6 (35.3%) |  |
| 2 | 9 (47.4%) | 10 (58.8%) |  |

Values are median [interquartile range], mean (standard deviation), or n (%). P values compare men and women using Wilcoxon rank-sum tests or unpaired t-tests for continuous variables and Fisher exact or chi-square tests for categorical variables, as appropriate. Missing values were omitted per variable. Histological categories follow the NASH Clinical Research Network classification.

Clinical characteristics of the 34 patients retained in the liver dataset are shown in Supplementary Table 3. Quality-control, library integration, clustering, sample-composition and cluster-level QC diagnostics supported the use of both datasets for downstream sex- and MASLD-stratified comparisons (Extended Data Fig. 1p1-p5).

We first examined whether the two datasets captured the expected tissue-specific cellular architecture. The liver dataset resolved 32 detailed populations spanning hepatocytes, liver sinusoidal endothelial cells, hepatic stellate and portal stromal populations, cholangiocytes, Kupffer and monocyte-derived phagocytes, lymphoid populations and rare vascular, mesothelial, neural and plasmacytoid dendritic populations (Fig. 1c). The visceral-fat dataset resolved 27 detailed populations spanning macrophage, fibroblast, mesothelial, endothelial, adipocyte, pericyte, lymphoid and mast-cell populations, together with a separate group of low-quality nuclei that was not assigned a biological identity (Fig. 1d). Major cell populations were represented across patient libraries rather than being driven by individual samples (Extended Data Fig. 1p4). Thus, the two datasets captured the distinct cellular organization of liver and visceral fat.

The broad cellular organization of the two tissues differed markedly. The liver dataset was dominated by hepatobiliary cells (104,964 nuclei; 72%), with smaller structural (stromal and vascular; 27,275 nuclei; 19%) and immune (14,295 nuclei; 10%) compartments. In contrast, VAT was divided almost equally between structural (stromal, vascular and adipocyte-associated; 176,237 nuclei; 47%) and immune cells (175,759 nuclei; 47%), with 22,316 low-quality nuclei (6%) retained as a separate labelled group (Fig. 1e). All major compartments occupied distinct regions of the liver and VAT UMAPs (Extended Data Fig. 1p6). Thus, liver was predominantly parenchymal, whereas VAT showed a more balanced structural and immune composition.

Canonical marker-gene expression confirmed the major cell populations. In liver, *ALB*, *STAB2*, *RGS5*, *MARCO* and *KRT19* marked hepatocytes, sinusoidal endothelium, stellate/stromal cells, Kupffer/myeloid cells and cholangiocytes, respectively. In visceral fat, *ADIPOQ*, *CD163*, *PDGFRA*, *VWF* and *WT1* marked adipocytes, macrophages, fibroblast/stromal cells, endothelial cells and mesothelial cells (Fig. 1f). Broader marker patterns across all annotated populations are shown in Extended Data Fig. 1p7,p8. Subset annotation further resolved finer populations within these compartments, including periportal, mid-lobular and pericentral hepatocytes (Extended Data Fig. 1p8). Together, these paired datasets establish the cellular reference frame for the tissue-specific and cross-tissue sex-by-MASLD analyses that follow.

### Liver MASLD differs by sex around hepatocyte and myeloid programs

We first examined where sex-associated transcriptional differences appeared within the liver dataset and whether they were already present without steatosis or were most apparent in MASLD. We compared women and men separately within the no-steatosis and MASLD groups. Differential-expression analysis showed limited and scattered sex differences in patients without steatosis (Extended Data Fig. 2a). In MASLD, a stronger signal localized to periportal and pericentral hepatocytes, with additional contributions from LSECs, Kupffer/myeloid cells and smaller populations (Fig. 2a and Extended Data Fig. 2b). Genes higher in women with MASLD were largely cell-type restricted rather than broadly shared across liver populations, supporting localized cell-state differences rather than a uniform tissue-wide sex effect (Extended Data Fig. 2c).

**Figure 2.**
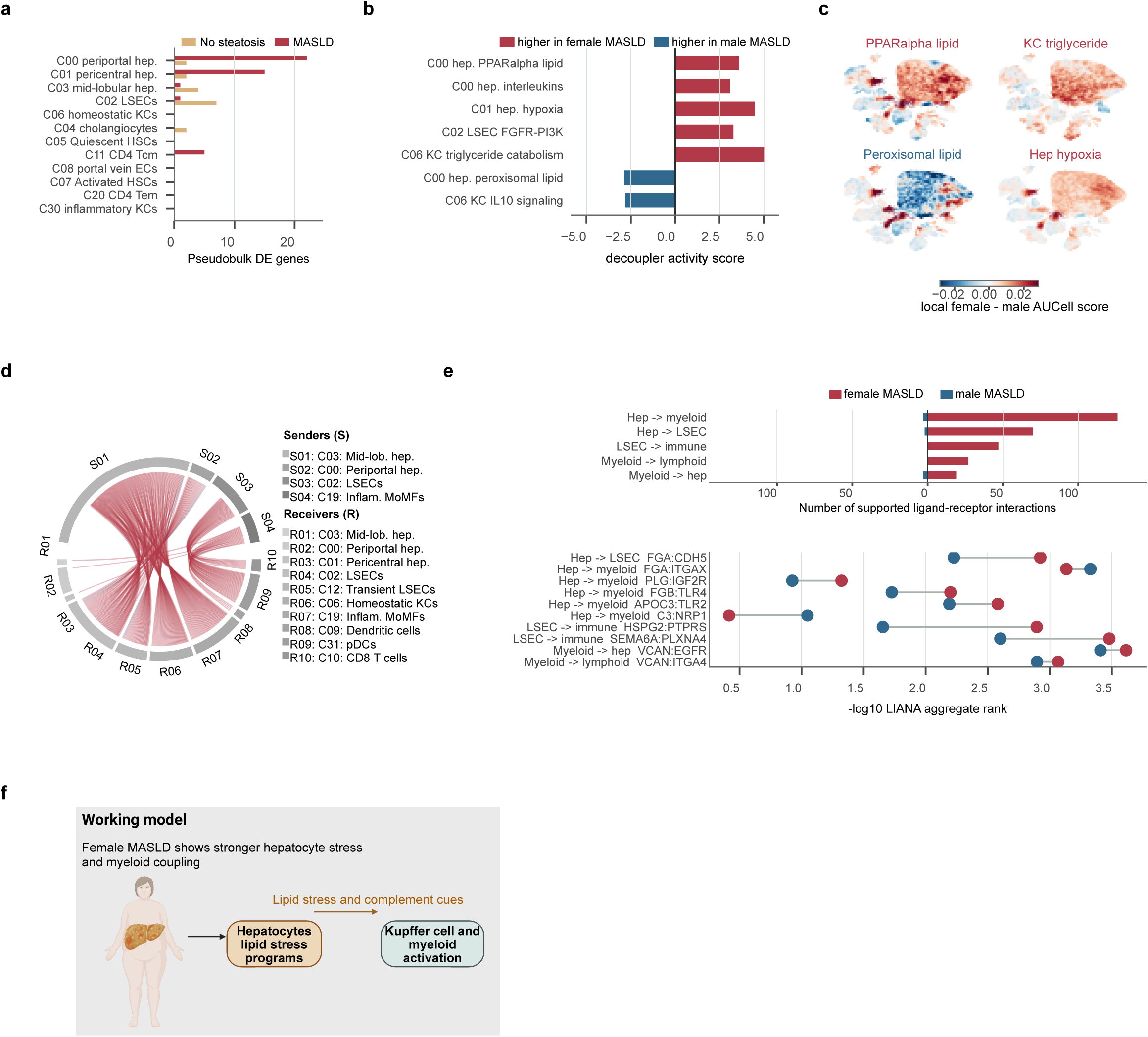
Sex-associated liver MASLD differences involve hepatocyte and myeloid programs. **a,** Female-versus-male liver differential-expression burden across liver r4 cell types in no steatosis and MASLD. Bars show pseudobulk-supported protein-coding DE gene counts on a linear scale after excluding X- and Y-chromosome genes. **b,** Selected MSigDB terms from the female.MASLD versus male.MASLD pseudobulk DE analysis, plotted as signed decoupler activity scores in hepatocytes, LSECs and Kupffer cells. Positive and negative values indicate higher activity scores in female and male MASLD, respectively. **c,** Local female-minus-male MASLD AUCell score differences for selected pathways from panel **b,** projected on the liver UMAP. **d,** LIANA-derived liver ligand-receptor communication pattern for MASLD across selected hepatocyte, LSEC, myeloid and lymphoid routes. S codes denote sender populations and R codes receiver populations. Edges were detected in both sex-specific MASLD tables and had an absolute female-minus-male lr_means difference greater than 0.25; color indicates direction and width indicates the absolute difference. **e,** Route-family summary and selected ligand-receptor examples from the panel-d edge set. **f,** Schematic summary of the sex-associated liver states in MASLD.

We then examined which biological programs distinguished women and men with MASLD in the liver. Pathway-activity scores showed higher PPARalpha-linked lipid regulation, interleukin signaling and hypoxia in hepatocytes from women with MASLD, together with higher FGFR-PI3K signaling in LSECs and triglyceride catabolism in Kupffer cells.

Hepatocytes and Kupffer cells from men with MASLD instead showed higher peroxisomal lipid metabolism and IL10-associated activity, respectively (Fig. 2b and Extended Data Fig. 2d-f). Patient-level AUCell scores agreed in direction for all seven displayed pathways.

Kupffer-cell triglyceride catabolism was higher in women (standardized difference, 1.30 SD; 95% CI, 0.59-2.01; FDR=0.0022), while hepatocyte interleukin signaling and hypoxia and LSEC FGFR-PI3K activity showed nominal support but did not pass FDR correction (Extended Data Fig. 2g). To determine which individual liver cells carried these pathway differences and how these cells were distributed, we projected female-minus-male MASLD score differences across the liver UMAP. The female-higher PPARalpha-linked lipid-regulation and hypoxia programs localized to hepatocyte regions of the UMAP, while triglyceride catabolism localized to the Kupffer-cell/myeloid region; peroxisomal lipid metabolism localized to hepatocytes (Fig. 2c and Extended Data Fig. 2h). Thus, the strongest patient-level pathway difference involved Kupffer-cell lipid processing, accompanied by directionally consistent hepatocyte and LSEC programs.

To determine whether the sex-associated hepatocyte, LSEC and immune states formed part of a coordinated multicellular response, we used LIANA^20^ to test whether ligands expressed in one population were matched by their cognate receptors in another. The liver interaction set, restricted to hepatocyte, LSEC, myeloid and lymphoid source-target routes, was predominantly higher in women with MASLD, with a smaller male-associated component (Fig. 2d,e). Leading female-higher candidates included PLG-*IGF2R*, *FGA*-*ITGAX*, *FGB*-*TLR4* and *APOC3*-*TLR2*, whereas *C3*-*NRP1* represented a recurrent male-higher interaction across hepatocyte zones (Extended Data Fig. 2i-k). Thus, women and men with MASLD differed not only in hepatocyte and myeloid pathway scores but also in the balance of inferred ligand-receptor routes linking these populations.

We also tested whether the sex-associated liver states were accompanied by detectable differences in cell-population abundance. No liver population was supported by both patient-level abundance methods in the female-versus-male MASLD comparison (Extended Data Fig. 2l). In this cohort, the clearest liver sex differences therefore involved transcriptional state and inferred communication rather than detectable population redistribution. Women with MASLD showed higher Kupffer-cell triglyceride-catabolism scores and a predominantly female-higher inferred interaction set, whereas male-associated hepatocyte peroxisomal and Kupffer-cell IL10 scores remained directional contrasts (Fig. 2f).

### Visceral-fat sex differences in MASLD organize around an adipocyte-macrophage axis

We compared women and men separately within the no-steatosis and MASLD groups to identify which VAT populations carried sex-associated transcriptional differences. In patients without steatosis, pseudobulk-supported differential expression was sparse and distributed among adipocyte, perivascular-fibroblast and lymphatic-endothelial populations, without a coherent adipocyte-macrophage program (Extended Data Fig. 3a). In MASLD, pseudobulk-supported differential expression was more extensive and localized primarily to mature adipocytes, homeostatic macrophages, mesothelial cells and monocyte-derived macrophages, with smaller endothelial and resident-macrophage contributions (Fig. 3a and Extended Data Fig. 3b). Genes higher in women with MASLD were entirely population specific rather than broadly shared across VAT populations, supporting distinct adipocyte and macrophage states rather than a uniform tissue-wide sex effect (Extended Data Fig. 3c).

**Figure 3.**
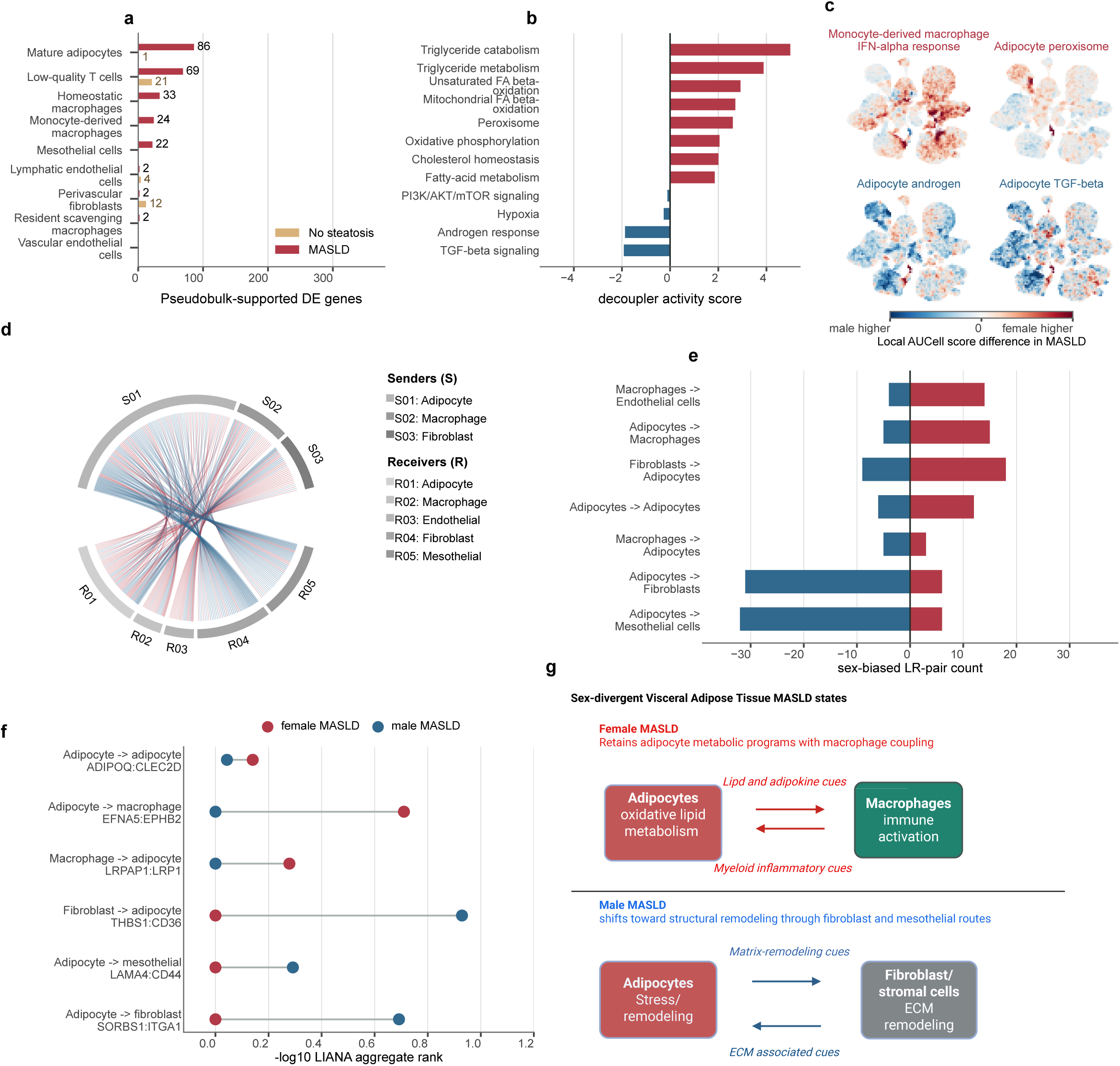
An adipocyte-macrophage axis characterizes sex differences in visceral adipose tissue in MASLD. **a,** Female-versus-male VAT differential-expression burden across r4 cell types in no steatosis and MASLD. Bars show pseudobulk-supported protein-coding DE gene counts after excluding X- and Y-chromosome genes. **b,** Selected MSigDB terms from the female.MASLD versus male.MASLD pseudobulk DE contrast in mature adipocytes, plotted as signed decoupler activity scores. Positive and negative values indicate higher activity scores in female and male MASLD, respectively. **c,** Local female-minus-male MASLD AUCell score differences for selected pathways from panel **b,** projected on the VAT UMAP. **d,** LIANA-derived VAT ligand-receptor communication pattern for MASLD across 166 selected adipocyte, macrophage, endothelial, fibroblast and mesothelial edges. S codes denote sender populations and R codes receiver populations. Color indicates direction and width indicates the absolute female-minus-male lr_means difference. **e,** Route-level counts of the panel-d edges higher in female and male MASLD. **f,** Six selected LIANA ligand-receptor examples from the panel-d edge set, displayed as aggregate-rank support in female and male MASLD. **g,** Schematic summary of the sex-associated VAT states in MASLD.

We then examined whether VAT from women and men with MASLD was characterized by distinct adipocyte and macrophage programs. Pathway-activity scores showed higher triglyceride metabolism, mitochondrial fatty-acid beta-oxidation, cholesterol homeostasis, peroxisome and oxidative phosphorylation in adipocytes from women with MASLD. Adipocytes from men with MASLD instead had higher decoupler activity scores for androgen response, TGF-beta, hypoxia and PI3K-AKT-mTOR-related pathways (Fig. 3b and Extended Data Fig. 3d). In women with MASLD, monocyte-derived macrophages showed higher interferon-alpha, interferon-gamma, IL6-JAK-STAT3 and complement activity, while homeostatic macrophages showed higher antigen-presentation, IL6 and IL12 signaling (Extended Data Fig. 3e,f). Patient-level AUCell analysis supported seven of the eight displayed female-higher adipocyte lipid and oxidative programs, with standardized differences of 0.87-1.15 SD and FDR<0.05; oxidative phosphorylation remained directionally higher but was not significant. The male-higher decoupler terms were less consistent at patient level: androgen response remained male-directed but did not pass FDR correction, TGF-beta was null, and hypoxia and PI3K-AKT-mTOR activity changed direction (Extended Data Fig. 3g). Per-cell maps localized interferon-alpha and peroxisome scores to macrophage-rich and adipocyte regions of the VAT UMAP, respectively, and showed the distribution of androgen-response and TGF-beta scores in adipocytes (Fig. 3c and Extended Data Fig. 3h). Thus, women with MASLD showed a robust adipocyte lipid-metabolic program together with inflammatory macrophage activation, whereas the male-associated adipocyte pathway pattern was less consistent across scoring approaches.

To determine whether these sex-associated adipocyte and macrophage states were accompanied by different local communication patterns, we examined ligand-receptor relationships within VAT. Inferred route scores were higher in women with MASLD for adipocyte self-routes, adipocyte-macrophage and macrophage-adipocyte routes, and macrophage-endothelial routes. Men with MASLD instead showed higher inferred adipocyte-fibroblast, adipocyte-mesothelial and fibroblast-adipocyte routes (Fig. 3d,e). Female-higher candidates comprised adipocyte self-signaling through *ADIPOQ*-*CLEC2D*, an adipocyte-to-macrophage *EFNA5*-*EPHB2* route and macrophage-to-adipocyte *LRPAP1*-*LRP1* feedback. Male-higher candidates instead included adipocyte-to-fibroblast *SORBS1*-*ITGA1* and other adipocyte-stromal routes involving *LAMA4*-*CD44* and *THBS1*-*CD36* (Fig. 3f and Extended Data Fig. 3i-k). Together, these inferred routes emphasized adipocyte-macrophage exchange in women and adipocyte-stromal exchange in men, as relative shifts within overlapping VAT communication patterns.

We also tested whether the contrasting VAT states were accompanied by detectable cell-population abundance differences. No VAT population was supported by both patient-level abundance methods in the female-versus-male MASLD comparison (Extended Data Fig. 3l). The clearest MASLD sex contrast in the single-nucleus dataset therefore involved cellular state and inferred communication rather than detectable population redistribution. Compared with men with MASLD, women with MASLD showed greater adipocyte lipid-metabolic activity, inflammatory macrophage programs and candidate adipocyte-macrophage communication, whereas the male-associated communication pattern favored fibroblast- and mesothelial-associated matrix remodeling (Fig. 3g).

### Paired liver and visceral adipose tissue analyses identify sex-associated interorgan coordination in MASLD

Because VAT and liver are metabolically connected, we next examined whether the sex-associated cellular programs identified in the two tissues covaried within the same patients. We defined focused gene modules using evidence including, for example, sex-associated differential-expression genes, pathway leading-edge genes and candidate ligand-receptor genes. The modules represented adipocyte signalling, adipocyte lipid metabolism and lipid release, macrophage activation, hepatocyte stress, hepatocyte lipid receipt, LSEC signaling and myeloid communication; full gene membership and provenance are provided in Supplementary Data 1. The labels summarize the dominant cell-state axis represented by each gene set; signalling and communication modules combine regulatory and ligand-sender or receiver genes rather than representing a single canonical pathway. UMAP projections confirmed that the modules localized to the expected VAT and liver populations (Extended Data Fig. 4a,b). These modules provided tissue-state measures for paired cross-tissue comparison.

We then tested whether VAT and liver states covaried within the same individuals. Across 32 matched VAT-liver pairs, VAT adipocyte signalling correlated positively with the liver hepatocyte stress-loop module (r=0.67, P=2.35e-5), and this association remained after accounting for differences among the sex and MASLD groups (partial r=0.61, P=4.28e-4).

Related positive correlations linked VAT adipocyte signalling to hepatocyte lipid receipt and VAT macrophage activation to hepatocyte stress (Fig. 4a and Extended Data Fig. 4c). By contrast, the male-associated VAT adipocyte-stress module correlated inversely with the liver hepatocyte stress-loop module. Thus, both the female-associated adipocyte-signalling and macrophage states and the male-associated adipocyte-stress state tracked with liver programs, supporting coordinated patient-level variation between adipose and liver transcriptional states.

**Figure 4.**
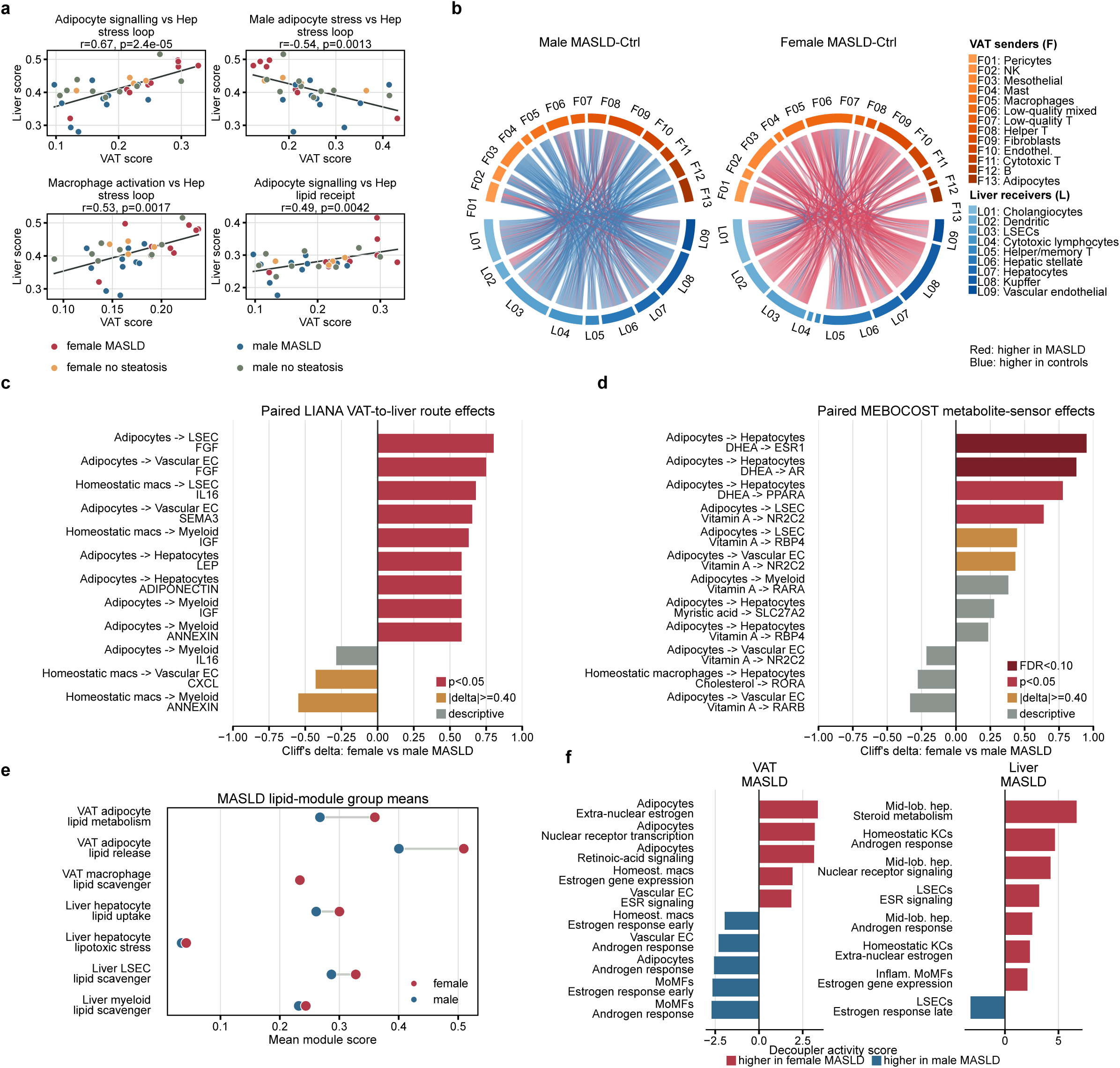
Paired liver and visceral adipose tissue analyses identify sex-associated interorgan coordination in MASLD. **a,** Patient-level scatterplots of the top cell-type-scoped module couplings across 32 matched VAT-liver pairs. Each point represents one patient; colors indicate sex and MASLD group, and trend lines show linear fits with 95% confidence intervals. **b,** Broad patient-level VAT-to-liver ligand-receptor chord summary across all annotated populations included in the screen. F codes denote VAT sender populations and L codes liver receiver populations. Chords show within-sex MASLD-versus-no-steatosis comparisons with nominal P<0.05 and an absolute patient-level mean rank-score difference of at least 0.10; red and blue indicate higher scores in MASLD and no steatosis, respectively. **c,** Focused patient-level ligand-receptor route effects across selected VAT and liver populations. Bars show Cliff’s delta for female versus male MASLD; positive values indicate higher patient-level scores in females. **d,** Focused metabolite-sensor event effects across the same VAT-to-liver comparison, summarized as Cliff’s delta and including candidate steroid, retinoid, fatty-acid and cholesterol-sensor events. **e,** Group-mean module activity for VAT adipocyte lipid-metabolism/lipid-release and liver lipid-receipt/stress programs in MASLD groups. **f,** Signed decoupler activity scores for steroid-metabolism, nuclear-receptor, ESR-related, androgen-response and retinoid programs in VAT and liver populations; positive values indicate higher activity in female MASLD.

To determine whether the coordinated VAT and liver states were accompanied by candidate cross-tissue communication routes, VAT ligands were matched to cognate receptors in liver populations at patient level using a consensus framework.^20^ Across all annotated populations, the strongest routes in the female comparison connected VAT fibroblast, immune and adipocyte populations with liver immune, endothelial and hepatocyte populations and were generally higher in MASLD, whereas the strongest routes in the male comparison were generally higher without steatosis (Fig. 4b). Focusing on metabolically relevant VAT and liver cell populations, we found higher adiponectin and SEMA3 route families in women than men with MASLD, whereas GAS, TGF-beta and annexin families were higher in men (Fig. 4c and Extended Data Fig. 4d,e). We subsequently tested which candidate VAT ligands had target networks matching the female-versus-male MASLD transcriptional programs within each liver population using ligand-target analysis.^21^ This supported *ADIPOQ*, *SEMA3A*, *SEMA3C* and *GAS6* as candidate links between VAT and liver transcriptional states (Extended Data Fig. 4f). Together, the female-associated communication pattern emphasized candidate adiponectin-mediated endocrine signaling together with semaphorin-associated vascular and immune communication, whereas the male-associated pattern emphasized GAS, TGF-beta and annexin routes associated with growth-factor signaling, tissue remodeling and stress responses. The contrast between adiponectin/semaphorin and GAS/TGF-beta/annexin route families prompted us to ask whether inferred metabolite-sensor communication showed a corresponding sex difference.

We therefore examined whether the sex difference extended to inferred metabolic cues from VAT to liver. Metabolite-sensor analysis matches metabolite-producing enzymes in VAT populations with metabolite-responsive sensors in liver populations.^22^ The strongest female-higher events linked VAT adipocytes with hepatocytes through DHEA-*ESR1* and DHEA-*AR*, with DHEA-*PPARA* and vitamin A-*NR2C2*/*RBP4* providing additional support. Myristic acid- *SLC27A2* represented a weaker adipocyte-to-hepatocyte candidate. Male-higher events were fewer and included a homeostatic-macrophage-to-hepatocyte cholesterol-*RORA* route and selected adipocyte-to-vascular-endothelial vitamin-A routes (Fig. 4d). Because several prioritized metabolites were lipid-derived or acted through lipid-regulatory nuclear receptors, we also examined whether these candidate routes occurred within broader sex differences in VAT and liver lipid handling. VAT adipocyte lipid-metabolism and lipid-release programs and liver lipid-receipt and stress programs were highest in women with MASLD (Fig. 4e). Women with MASLD therefore combined adipocyte lipid-metabolism and lipid-release transcriptional programs with candidate steroid and retinoid signaling toward hepatocytes and LSECs, alongside hepatic lipid receipt and stress. Men with MASLD instead showed a narrower metabolic pattern centered on macrophage cholesterol and vascular retinoid routes.

Finally, we assessed whether the candidate steroid and retinoid routes were accompanied by broader hormone-responsive programs in the corresponding VAT and liver populations. We used focused pathway-activity scores to examine these programs.^23^ Adipocytes from women with MASLD showed higher extra-nuclear estrogen, nuclear-receptor and retinoic-acid activity, whereas adipocyte androgen-response activity was higher in men. In liver, women with MASLD showed higher steroid-metabolism and nuclear-receptor activity in hepatocytes and higher ESR- and androgen-related activity across LSECs and homeostatic Kupffer cells, whereas late estrogen-response activity in LSECs was higher in men (Fig. 4f). Patient-level gene expression supported this cellular localization, including higher *ESR1*, *NR3C1*, *AKR1C1* and *STS* in female VAT adipocytes, steroid-processing genes in female hepatocytes, and *ESR1*, *PPARA* and *SORT1* in female liver myeloid cells (Extended Data Fig. 4g-i). Thus, women with MASLD showed coordinated steroid, retinoid and nuclear-receptor activity across VAT and liver, whereas male-higher hormone-response activity was concentrated in adipocyte, macrophage and LSEC programs.

Together, the coupled transcriptional states and candidate communication routes showed partly different patterns of VAT-liver coordination in women and men with MASLD. These sex-associated patterns extended to steroid, retinoid and lipid-related programs and involved different combinations of metabolic, endocrine, immune and structural activity.

### Bulk RNA-seq in a larger cohort supports the sex-stratified cell-state model

To determine whether the sex-associated cell states and VAT-liver relationships were also detectable at tissue level in a larger series, we analysed bulk RNA-seq data from a non-overlapping set of participants from the same study cohort. Liver and VAT profiles were available for approximately 300 participants, including 293 with paired tissues (225 women and 68 men; Fig. 5a; Table 2).

**Figure 5.**
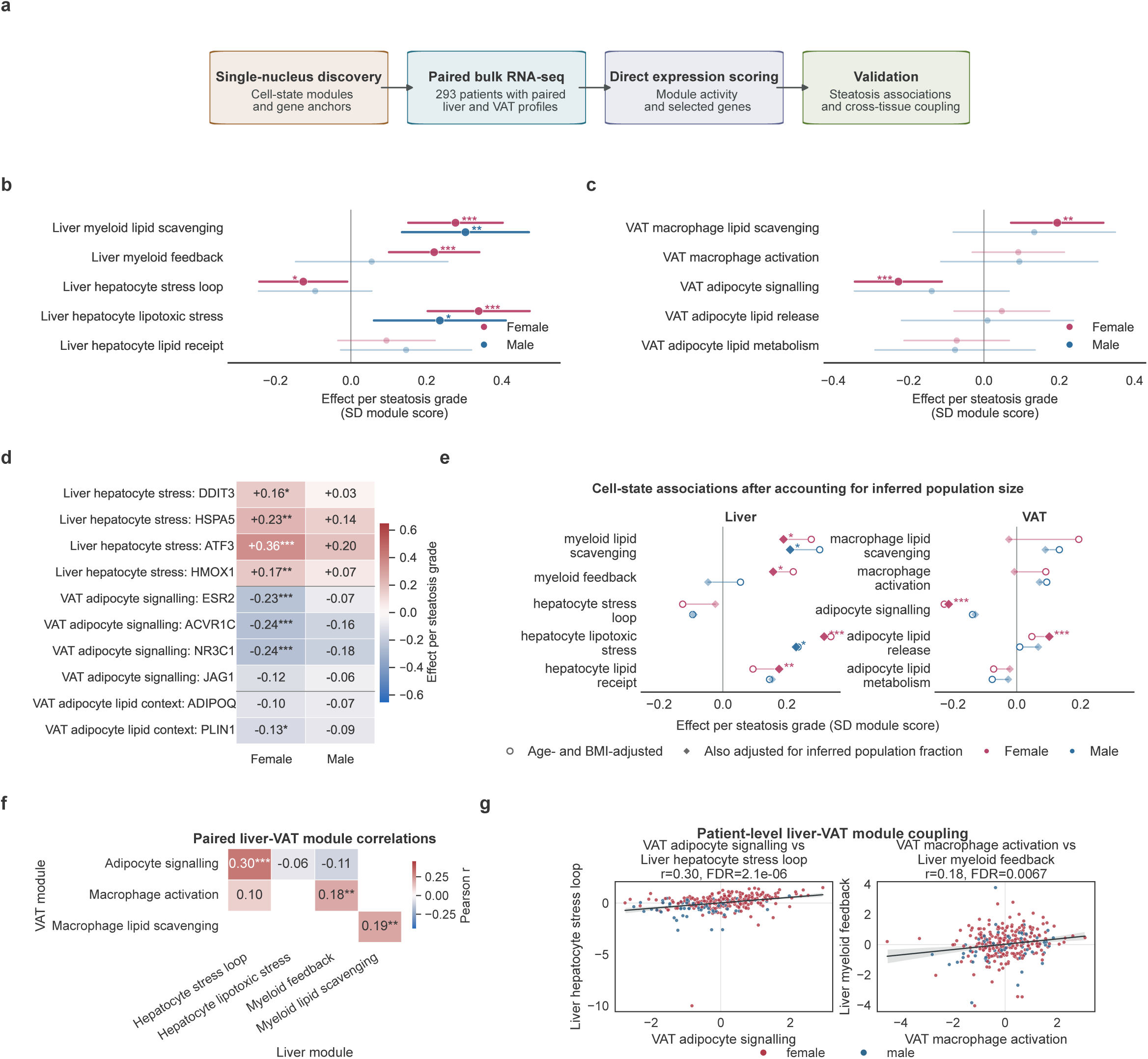
Bulk RNA-seq supports sex-stratified liver-VAT cell-state and coupling signals. **a,** Validation workflow. Cell-state modules and gene anchors derived from the single-nucleus analyses were scored directly in paired liver and VAT bulk RNA-seq, followed by sex-stratified steatosis association and patient-level cross-tissue coupling analyses. **b,** Liver module associations with steatosis grade, modelled separately by sex and adjusted for age and BMI. Effects are shown per one-grade increase in steatosis as SD module-score changes with 95% confidence intervals. **c,** VAT module associations with steatosis grade using the same model structure. **d,** Representative gene anchors from the corresponding single-nucleus-derived modules tested in the same sex-stratified steatosis models. Cell color and text show the effect per one-grade increase in steatosis as SD gene-expression changes. **e,** Sensitivity analysis comparing the principal module-steatosis estimates before and after adjustment for the corresponding MuSiC-inferred population fraction in the same deconvolution-complete patients. Open circles show age- and BMI-adjusted estimates; filled diamonds show estimates additionally adjusted for inferred population fraction. Lines connect estimates from the same module and sex. **f,** Focused Pearson correlation map for predefined liver-VAT module pairs across 293 patients with both tissues available. **g,** Patient-level examples for VAT adipocyte signalling versus the liver hepatocyte stress loop and VAT macrophage activation versus liver myeloid feedback. Each point is a patient with paired liver and VAT profiles; lines and shaded regions show the pooled linear fit and 95% confidence interval. Faded forest-plot estimates indicate non-FDR-significant associations; stars denote FDR levels. Sex-by-steatosis interaction tests for module slopes and module-coupling-by-sex interaction tests are reported in the source tables.

**Table 2.** Clinical characteristics of the paired bulk RNA-seq cohort. The table describes the 293 patients with paired liver and visceral adipose tissue bulk RNA-seq profiles used for cross-tissue analyses.

| Characteristic | Men (n=68) | Women (n=225) | P value |
| --- | --- | --- | --- |
| **Demographics** |  |  |  |
| Age, years | 52.00 [45.00, 56.25] | 47.00 [38.00, 53.00] | <0.001 |
| BMI, kg/m<sup>2</sup> | 39.28 [36.73, 42.68] | 38.97 [36.49, 41.24] | 0.672 |
| Diabetes, n (%) | 14 (20.6%) | 27 (12.0%) | 0.112 |
| No hypertension, n (%) | 31 (45.6%) | 154 (68.4%) | 0.001 |
| MASLD status, n (%) |  |  | 0.011 |
| No steatosis | 23 (33.8%) | 121 (53.8%) |  |
| MASLD | 31 (45.6%) | 78 (34.7%) |  |
| MASH | 14 (20.6%) | 26 (11.6%) |  |
| **Clinical laboratory values** |  |  |  |
| Alkaline phosphatase, U/L | 76.00 [65.25, 93.50] | 82.00 [67.00, 98.00] | 0.076 |
| Gamma-glutamyl transferase, U/L | 43.00 [28.50, 62.00] | 26.00 [20.00, 38.50] | <0.001 |
| Aspartate aminotransferase, U/L | 29.00 [24.00, 35.00] | 23.00 [20.00, 28.00] | <0.001 |
| Alanine aminotransferase, U/L | 42.00 [30.00, 52.00] | 29.00 [22.00, 39.00] | <0.001 |
| Total cholesterol, mmol/L | 4.56 (1.18) | 4.90 (0.99) | 0.021 |
| LDL cholesterol, mmol/L | 2.92 (0.98) | 3.23 (0.93) | 0.024 |
| HDL cholesterol, mmol/L | 0.94 [0.82, 1.10] | 1.20 [1.03, 1.40] | <0.001 |
| Triglycerides, mmol/L | 2.42 (1.63) | 1.66 (1.21) | <0.001 |
| Glucose, mmol/L | 6.00 [5.50, 7.20] | 5.60 [5.10, 6.20] | <0.001 |
| HbA1c, % | 5.90 [5.50, 7.00] | 5.60 [5.30, 6.30] | 0.007 |
| C-reactive protein, mg/L | 3.30 [2.00, 6.85] | 6.15 [3.10, 9.77] | <0.001 |
| **Pathology** |  |  |  |
| Steatosis grade, n (%) |  |  | 0.009 |
| <5% | 22 (32.8%) | 120 (53.6%) |  |
| 5-33% | 12 (17.9%) | 38 (17.0%) |  |
| 33-66% | 23 (34.3%) | 40 (17.9%) |  |
| >66% | 10 (14.9%) | 26 (11.6%) |  |
| Fibrosis stage, n (%) |  |  | 0.121 |
| 0 | 9 (13.4%) | 52 (23.2%) |  |
| 1 | 16 (23.9%) | 60 (26.8%) |  |
| 2 | 26 (38.8%) | 76 (33.9%) |  |
| 3 | 13 (19.4%) | 34 (15.2%) |  |
| 4 | 3 (4.5%) | 2 (0.9%) |  |
| Ballooning grade, n (%) |  |  | 0.309 |
| 0 | 51 (76.1%) | 186 (83.0%) |  |
| 1 | 13 (19.4%) | 27 (12.1%) |  |
| 2 | 3 (4.5%) | 11 (4.9%) |  |
| Lobular inflammation grade, n (%) |  |  | 0.580 |
| 0 | 0 (0.0%) | 5 (2.2%) |  |
| 1 | 8 (11.9%) | 31 (13.8%) |  |
| 2 | 43 (64.2%) | 131 (58.5%) |  |
| 3 | 16 (23.9%) | 57 (25.4%) |  |

Values are median [interquartile range], mean (standard deviation), or n (%). P values compare men and women using Wilcoxon rank-sum tests or unpaired t-tests for continuous variables and Fisher exact or chi-square tests for categorical variables, as appropriate. Missing values were omitted per variable. Histological categories follow the NASH Clinical Research Network classification.

We first examined whether steatosis grade in the larger cohort was associated with broad differences in tissue composition. We selected MuSiC as the best-fitting deconvolution approach (Extended Data Fig. 5a) and used the single-nucleus datasets as references to estimate broad liver and VAT population fractions in the bulk profiles (Extended Data Fig. 5b,c). Across both tissues, steatosis grade showed little evidence of broad population redistribution. A higher inferred VAT macrophage fraction was associated with steatosis in women, but this association did not differ significantly between sexes (Extended Data Fig. 5d-g). Thus, the larger cohort again pointed primarily to cellular-state differences rather than widespread shifts in population abundance.

We then tested whether the cell-state programs identified by single-nucleus analysis were reflected in bulk tissue as steatosis increased. The corresponding gene modules were scored directly from bulk expression and associated with steatosis grade separately by tissue and sex (Fig. 5b,c). Liver hepatocyte lipotoxic stress increased with steatosis grade in women (beta per grade, 0.34 SD; 95% CI, 0.20-0.47; FDR=4.0e-6) and men (beta, 0.24 SD; 95% CI, 0.06-0.41; FDR=0.041). Liver myeloid lipid scavenging also increased in women (beta, 0.28 SD; 95% CI, 0.15-0.40; FDR=5.6e-5) and men (beta, 0.30 SD; 95% CI, 0.14-

0.47; FDR=0.0048), whereas liver myeloid feedback was associated with steatosis in women (beta, 0.22 SD; 95% CI, 0.10-0.34; FDR=7.5e-4) but not men (Fig. 5b). Among women, increasing steatosis was associated with lower VAT adipocyte-signalling activity (beta, -0.23 SD; 95% CI, -0.34 to -0.11; FDR=5.6e-4) and higher VAT macrophage lipid scavenging (beta, 0.19 SD; 95% CI, 0.07-0.32; FDR=0.0066), whereas the broader VAT adipocyte lipid-metabolism module was not associated with steatosis in either sex (Fig. 5c).

To identify individual genes anchoring the stress, lipid-handling and adipocyte-signalling programs detected in bulk tissue, we tested selected gene-level associations with steatosis. In liver samples from women, increasing steatosis was associated with higher expression of the hepatocyte stress genes *HSPA5* (beta, 0.23 SD; 95% CI, 0.10-0.36; FDR=0.0011), *ATF3* (beta, 0.36 SD; 95% CI, 0.20-0.52; FDR=5.7e-5), *HMOX1* (beta, 0.17 SD; 95% CI, 0.06-0.27; FDR=0.0024) and *DDIT3* (beta, 0.16 SD; 95% CI, 0.04-0.28; FDR=0.013) (Fig. 5d).

Together, *HSPA5*, *ATF3* and *DDIT3* marked chaperone, integrated-stress and unresolved-stress components of the endoplasmic-reticulum stress response, while *HMOX1* marked oxidative stress. *TREM2* and *LIPA* marked lipid-associated myeloid remodeling and lysosomal lipid processing; liver *TREM2* increased with steatosis in both sexes, while a broader lipid-associated myeloid marker panel showed more extensive associations in women across liver and VAT (Extended Data Fig. 5h). In VAT samples from women, increasing steatosis was accompanied by lower expression of the adipocyte-signalling genes *ESR2*, *ACVR1C* and *NR3C1*. Among metabolite-related genes, VAT *STS* increased and *ADH1B* decreased with steatosis in women, whereas liver sensor-gene associations were weaker and did not pass FDR correction (Extended Data Fig. 5i). Thus, although the single-nucleus comparison identified higher hormone-responsive adipocyte features in women than men with MASLD, the larger cohort showed that, within women, *ESR2*, *ACVR1C*, *NR3C1* and the broader adipocyte-signalling program decreased as steatosis grade increased. This decline accompanied hepatocyte endoplasmic-reticulum stress and myeloid lipid processing, while part of the myeloid response was shared with men.

Because bulk module scores can reflect both expression within a population and the abundance of that population, we repeated the central models after accounting for the corresponding inferred population fraction. This preserved hepatocyte lipotoxic stress and myeloid lipid scavenging in both sexes, liver myeloid feedback in women, and the inverse association between steatosis and VAT adipocyte signalling in women (Fig. 5e). The VAT macrophage lipid-scavenging association in women was attenuated, whereas the adipocyte lipid-release estimate strengthened after adipocyte-fraction adjustment. Several module scores also correlated with the inferred abundance of their matching population (Extended Data Fig. 5j,k). These analyses supported the principal liver stress and VAT adipocyte-signalling findings as cell-state associations, while indicating a greater contribution of tissue composition to parts of the VAT macrophage and adipocyte lipid signal.

Finally, we examined whether paired liver and VAT bulk profiles preserved the cross-tissue relationships observed in the single-nucleus cohort. The focused correlation map showed covariation between adipocyte-hepatocyte and macrophage-myeloid programs across the paired cohort (Fig. 5f). VAT adipocyte signalling correlated with the liver hepatocyte stress-loop program (Pearson r=0.30; FDR=2.1e-6), and this association remained after adjustment for age, BMI and steatosis grade (adjusted r=0.25; FDR=2.0e-4) (Fig. 5g and Extended Data Fig. 5l). VAT macrophage activation correlated with liver myeloid feedback (r=0.18; FDR=0.0067), and VAT macrophage lipid scavenging correlated with liver myeloid lipid scavenging (r=0.19; FDR=0.0067). At the individual-gene level, correlations between candidate metabolite-producing genes in VAT and sensor genes in liver were modest and did not pass FDR correction, whereas matched lipid-handling and scavenger genes showed broader cross-tissue associations in women, including *TREM2*, *LIPA* and *GPNMB* (Extended Data Fig. 5m,n). Thus, adipocyte and macrophage programs in VAT covaried with liver stress and myeloid activity across patients, with adipose-liver coordination more evident at the level of transcriptional programs than individual metabolite-sensor genes.

Together, bulk RNA-seq data from the larger cohort supported hepatic stress and myeloid lipid handling as features of steatosis in both sexes. Among women, increasing steatosis was additionally associated with lower VAT adipocyte-signalling activity and greater liver myeloid feedback, while the broader VAT adipocyte lipid-metabolism module remained unchanged. Together, the population-adjusted associations of these single-nucleus-derived programs and their coupling across paired VAT and liver profiles supported coordinated cell-state organization across the two tissues in sex-stratified analyses.

## Discussion

MASLD arises within systemic metabolic dysfunction involving both liver and adipose tissue. In this cohort, the strongest transcriptional separation emerged when women and men with MASLD were compared directly across matched liver and visceral adipose tissue (VAT).

Compared with men with MASLD, women with MASLD showed coordinated VAT adipocyte and macrophage programs together with hepatocyte stress, lipid-receipt and liver myeloid programs. These findings support partly different tissue-state and candidate inter-organ communication patterns in women and men with MASLD.

This finding fits an emerging literature in which sex modifies MASLD risk, progression and molecular state. Clinical and experimental studies have emphasized the role of estrogen, hepatic estrogen-receptor signaling and broader sex-associated lipid and immune biology in shaping liver disease susceptibility.^7,24^ Recent human liver atlases show that healthy sex- and age-associated variation and MASLD progression are organized through cell-type-specific states and, in disease, spatial niches.^25,26^ Our data extend this framework by resolving sex-stratified MASLD programs simultaneously in liver and VAT from the same individuals and by identifying coordinated, tissue-specific cellular programs across the two organs.

This cross-tissue perspective was enabled by profiling liver and VAT at cellular resolution in the same individuals. We analysed the tissues as separate datasets to preserve their organ-specific cellular architecture, then linked them through patient-matched transcriptional programs and communication analyses.

The liver findings position sex as a source of biological heterogeneity within MASLD. Spatial and single-cell studies have described lipid-associated macrophages, hepatocyte metabolic and stress programs, endothelial-stellate communication and zonated metabolic remodeling across MASLD stages.^25^ Our data extend this framework by showing that comparable MASLD disease strata can be organized through different hepatocyte, myeloid and intercellular signalling states in women and men. In women, continued hepatocyte lipid handling occurred alongside stress-associated activity and fibrinogen-linked immune signalling, whereas the male-associated pattern emphasized peroxisomal metabolism, Kupffer-cell IL10 activity and complement-linked communication. These patterns are better understood as distinct transcriptional and intercellular wiring within MASLD rather than evidence that women had simply progressed further.

The VAT findings extend this principle beyond the liver. In women with MASLD, greater oxidative and lipid-metabolic adipocyte activity co-occurred with inflammatory macrophage programs, whereas men showed stronger adipocyte stress and stromal-remodeling programs. The limited evidence for broad abundance differences again points primarily to altered states within existing cellular compartments.

The macrophage component of this VAT state should be interpreted carefully. Human and mouse adipose studies have made lipid-associated macrophage states central to obesity biology, and published human adipose atlases support the idea that adipose immune and stromal niches are remodeled across obesity, depot-specific and weight-loss contexts.^27–30^ However, our VAT dataset does not show a clean standalone *TREM2*-high LAM or inflammatory LAM branch. The absence of a separate branch may reflect depot- and disease-context differences in macrophage organization, while single-nucleus sampling and discrete clustering may also distribute a relatively sparse or continuous LAM-like state across neighboring macrophage populations. Lipid-associated features, including genes such as *GPNMB*, *LPL*, *CD9*, *SPP1*, *FABP5* and *LGALS3*, are distributed mainly across monocyte-derived macrophages and, to a lesser extent, resident scavenging macrophages. The female-associated VAT signal in MASLD is therefore better described as a mixed resident and monocyte-derived macrophage program than as canonical LAM expansion.

The paired analysis connected these tissue-specific states at patient level. Module and communication analyses showed associations consistent with endocrine, growth-factor and lipid-related coordination between adipose tissue and liver, in line with broader adipose-liver crosstalk literature and experimental adipocyte-hepatocyte systems.^31–33^ The patient-matched design resolves these associations across defined cell populations in both tissues rather than only in bulk tissue, circulating markers or single-organ profiles.

The larger bulk RNA-seq cohort provided orthogonal support for the central liver-stress, adipocyte-signalling and macrophage programs, with little evidence for broad population redistribution as steatosis increased. The female VAT associations were program specific: adipocyte signalling declined with increasing steatosis, whereas the broader adipocyte lipid-metabolism module remained unchanged. This agreement with the single-nucleus analyses was strongest at the level of coordinated cell-state modules rather than individual metabolite-sensor transcript pairs.

The translational relevance of these findings lies primarily in patient stratification and study design. The data do not support treating sex only as a covariate to adjust for after disease groups have been defined. Instead, women and men can occupy overlapping clinical MASLD categories while differing in the distribution of tissue programs. Future interventional studies could therefore benefit from prespecified sex-stratified tissue, biomarker and histology readouts, particularly for therapies aimed at adipose function, macrophage inflammation, lipid handling, nuclear-receptor signaling or adipose-liver endocrine crosstalk. The present data identify cellular programs that could be followed in trials or mechanistic studies to ask whether the same intervention normalizes the same tissue axis in women and men.

Because of some limitations of the current study, several open questions remain. The tissue data are cross-sectional and therefore do not capture longitudinal remodeling. Because all participants had severe obesity, the no-steatosis group represents a metabolically affected comparison group without histological steatosis rather than a healthy control group. Although the single-nucleus groups were balanced by sex and MASLD status, their patient-level size limited sensitivity to smaller transcriptional and abundance differences. The paired design and balanced single-nucleus groups strengthen the sex-stratified comparison, but the inferred endocrine and lipid axes will require direct validation. Histology, spatial profiling and targeted hormone, metabolite or lipid measurements will therefore be important to test how these transcriptional programs relate to hepatocyte zonation, physical adipose and liver niches, fibrosis patterns, portal inflammatory burden and measurable metabolite flux.

Together, these data support a model in which MASLD is organized by sex, tissue context and inter-organ coordination. The paired tissue programs position MASLD as a sex-stratified manifestation of systemic metabolic dysfunction rather than a uniform liver phenotype.

## Methods

### Study cohort and tissue collection

This study used samples from 42 participants with severe obesity who met the BARIA eligibility criteria for primary bariatric surgery: BMI ≥40 kg m^−2^ or BMI ≥35 kg m^−2^ with an obesity-related comorbidity. The study was approved by the Amsterdam UMC Medical Ethics Committee (BARIA study, trial registration NL8983), and all participants provided written informed consent. Visceral adipose tissue (VAT) was sampled from the appendices epiploicae, and liver biopsies were collected during the same surgical procedure. Complete cohort inclusion and exclusion criteria are described in the cohort design paper.^19^

Liver histology was classified according to the NASH Clinical Research Network (NASH CRN) criteria.^34^ HistoIndex was used for digital pathology scoring.^35^ Histological labels distinguished no steatosis, MASLD and metabolic dysfunction-associated steatohepatitis (MASH). For the primary single-nucleus contrasts, MASLD and MASH labels were collapsed into a steatotic MASLD category and compared with no steatosis unless stated otherwise.

### Single-nucleus RNA-sequencing preprocessing

Frozen liver and VAT biopsies were processed as separate single-nucleus RNA-sequencing datasets. Liver libraries were generated with the SeekGene platform and FASTQ files were processed with the SeekGene SeekSoul Tools software stack.^36^ VAT libraries were generated with the 10x Genomics platform and processed per library with Cell Ranger v10.0.0.^37^ Intronic reads were included, consistent with single-nucleus RNA-sequencing, and reads were aligned to a GRCh38 reference transcriptome. Platform-specific output matrices were then processed with CellBender v0.3.0 using remove-background --cuda with default parameters.^38^ The CellBender-filtered matrices were used as the count source for downstream dataset construction.

We selected 40 liver and 40 VAT samples for single-nucleus RNA sequencing, comprising 20 women and 20 men per tissue. The tissue-specific selections were not completely overlapping and together represented 42 unique patients. One liver library failed pre-sequencing quality control. Of the remaining 39 sequenced liver libraries, five were excluded during post-sequencing quality control, leaving 34 liver libraries from 14 women and 20 men. All 40 VAT libraries were retained, and 32 patients had usable data from both tissues.

Downstream analyses used the binary contrast of no steatosis versus MASLD, with MASLD and MASH combined in the MASLD group.

### Quality control and dataset construction

CellBender-filtered matrices were loaded with scomnom load-and-filter using sample_id as the batch key. scOmnom is a modular scVerse-based single-cell analysis workflow; documentation for the public workflow and command structure is available at https://prangelab.org/scomnom.^39^ The workflow was run largely with default parameters. The explicit upstream filtering settings were --min-genes 200 --max-pct-mt 30. Nuclei with fewer than 200 detected genes or more than 30% mitochondrial counts were removed. Default upper-tail filters were applied for detected genes and total counts, and genes detected in fewer than three nuclei were removed. Raw and CellBender-filtered count layers were retained where available for QC comparison and CellBender effect diagnostics.

Doublets were scored with SOLO after scVI model fitting.^40–42^ The expected doublet rate was set to 0.1, and sample-specific score thresholds were inferred from the doublet-score distributions. After QC and doublet filtering, the final liver dataset contained 146,534 nuclei from 34 libraries and 31,023 genes. The final VAT dataset contained 374,312 nuclei from 40 libraries and 30,968 genes. Median retained nuclei per library were 3,126 for liver and 9,218.5 for VAT.

### Tissue-specific integration and annotation

Liver and VAT were integrated independently rather than combined into a single cross-tissue embedding. For each tissue, scomnom evaluated Harmony, Scanorama, scVI and scANVI integration outputs and used scIB benchmark metrics to select the best-performing embedding for downstream analysis.^40,42–45^ Harmony was selected for the liver dataset and Scanorama was selected for the VAT dataset.

Clustering was performed after integration on the selected tissue-specific embedding. The scomnom Biology Informed Structural Clustering (BISC) step was used as a benchmark-informed Leiden resolution-selection procedure, after which closely related clusters were compacted into a broad annotation backbone for manual review and subset refinement. In liver, BISC-guided resolution selection identified Leiden resolution 0.5 for the initial clustering. In VAT, the active clustering report recorded 11 compacted broad clusters before subset refinement. For both tissues, cluster-size distributions, sample and batch composition, silhouette diagnostics, stability plots and UMAP overlays were used to assess integration and clustering behavior.

CellTypist was used as an orientation layer rather than as the final annotation source.^46^ The liver integration used Healthy_Human_Liver.pkl as the CellTypist model, whereas VAT used the default Immune_All_High.pkl model. Non-immune labels, especially in the immune-centered VAT model, were treated as provisional and refined using marker evidence. After initial compacted annotation, a secondary annotated scANVI run was trained from the compacted label structure in each tissue to generate a cleaner display projection while preserving the tissue-specific annotation framework.^42^

Initial tissue-level clusters were assigned broad identities, after which major compartments were subsetted, reclustered and reannotated. Annotation decisions used marker gene expression, pathway and regulator summaries, QC diagnostics and manual review of expected tissue architecture. Subset-derived labels were merged back into the main tissue objects. The final detailed annotation layer was r4_subset_annotation in both tissues, with higher-level archetype, compartment and supercompartment layers used for presentation and integration.

Final scOmnom AnnData archives were stored as compressed Zarr archives for downstream analysis and figure generation.

### Marker discovery and annotation support

Marker discovery was run with the scomnom markers-and-de workflow on the tissue-specific annotated datasets. Cell-level marker testing used unpaired Wilcoxon rank-sum tests comparing each candidate cluster with all other nuclei within the same tissue. Patient identity was not modelled in this supportive cell-level test; the accompanying pseudobulk marker analysis accounted for target and rest aggregates from the same patient by including sample in the design. Pseudobulk marker testing aggregated the CellBender count layer by sample and candidate cluster, then tested the binary cluster membership contrast with a sample-adjusted design. The marker design was sample + binary_cluster for pseudobulk cluster markers. Within each cluster-versus-rest marker test, cell-level P values were adjusted across genes using the Benjamini-Hochberg false-discovery-rate procedure. Pseudobulk adjusted P values were obtained from PyDESeq2 using its default independent filtering and Benjamini-Hochberg correction. Cell-level and pseudobulk marker evidence were inspected together during annotation, with pseudobulk summaries used to reduce sensitivity to unequal cell numbers across patients. Focused marker violins were generated with om.plotting.plot_de_violin_grid_genes using the final archetype annotation layer. The selected marker genes were scored directly on the final objects and recorded in the Figure 1 asset tables.

### Differential expression and abundance

Differential-expression analyses were run after final annotation merging. Female-versus-male DE summaries for liver and VAT were taken from the corresponding round-3 r4 differential-expression outputs. Cell-level DE used Wilcoxon and logistic-regression tests within the relevant annotation stratum. Pseudobulk DE aggregated CellBender counts by patient within each annotated population and tested the specified biological contrast using the corresponding sample-level design formula. The main sex, MASLD, sex-within-MASLD and interaction designs were sample + sex, sample + MASLD, sample + MASLD.sex and sample + sex + MASLD + sex:MASLD, respectively. For sex-stratified DE calls used in the DE-burden and DE-overlap support panels, genes were filtered before testing with --gene-filter “gene_chrom not in [’X’,’Y’]” and --gene-filter “gene_type == ’protein-coding’”. Cell-level DE was retained as supportive context, whereas pseudobulk DE was treated as the primary inferential layer when available.

DE-burden panels used pseudobulk-supported DE counts on a linear axis. DE-overlap support panels used cluster-level pseudobulk DE tables for the focal sex contrasts. In VAT, the overlap diagnostic was restricted to female-biased pseudobulk DE genes with adjusted P value below 0.05 and summarized whether the same genes recurred across VAT cell-type groups. These overlap panels were used as diagnostics for recurrent signal structure and broad shared technical artifacts; they were not used to define pathway or communication claims.

Differential-abundance analyses were run after the final marker and DE workflows and interpreted in the context of the finalized tissue-specific annotation layers. DA context was taken from existing liver and VAT r4 composition outputs for pooled sex and sex within MASLD. Patient-level abundance differences were assessed using centered-log-ratio (CLR) testing and scCODA.^47^ Supplemental summaries used the exact female-versus-male coefficient for each comparison and reported method-specific candidates and their overlap. CLR candidates required FDR<0.05 and scCODA candidates required inclusion probability at least 0.95. No population in either tissue was supported by both methods in the female-versus-male MASLD comparison.

### Pathway and regulator analysis

Pathway and activity summaries were generated with decoupler resources where available, including MSigDB Hallmark and Reactome gene sets, DoRothEA and PROGENy.^23,48–50^ Pathway evidence for the liver and VAT sex-stratified panels was taken from pseudobulk decoupler MSigDB activity tables for the focal female.MASLD_vs_male.MASLD contrast.

Selected terms were displayed as signed decoupler activity scores, with positive values following the female MASLD direction and negative values following the male MASLD direction. Separate supplemental panels showed unbiased top positive and negative MSigDB terms for selected cell types.

In the liver, selected pathway terms summarized hepatocyte, LSEC and Kupffer-cell biology while retaining male-enriched contrasts. In VAT, the main pathway panel was restricted to mature adipocytes and macrophage populations to summarize adipocyte metabolic-retention, macrophage immune-activation and male adipocyte remodeling contrasts.

For broader cross-tissue pathway interpretation, gene-level and pathway-level evidence was taken from the tissue-specific scomnom differential-expression workflows. Sex-stratified MASLD contrasts were evaluated within the relevant annotated cell populations. Preranked GSEA was run through the scomnom MSigDB GSEA layer using gseapy v1.1.12.^51,52^ Decoupler results were treated as directional activity scores, whereas the GSEA layer supplied enrichment significance and leading-edge genes. A joint evidence layer was used to identify pathway terms with concordant decoupler direction and significant preranked GSEA support. Pathway claims were therefore graded by concordance between gene-level DE, decoupler activity, GSEA significance and leading-edge support.

### Per-cell pathway and module scoring

Selected MSigDB gene sets from the liver and VAT pathway panels were extracted from MSigDB. Per-cell pathway activity was scored with scomnom.markers_and_de.module_score(…, module_score_method=“aucell”), which uses the rank-based decoupler AUCell implementation.^23,53^ Local female-minus-male MASLD maps were generated by computing local mean pathway scores in UMAP neighborhoods separately among female MASLD and male MASLD cells, then plotting the local score difference. The VAT maps used up to 80 nearest neighbors per sex group.

Additional gene modules were curated from project evidence rather than from unconstrained literature gene lists. Candidate genes were annotated against sex-stratified MASLD DE tables, decoupler pathway activity, significant GSEA leading-edge membership, LIANA ligand-receptor evidence and MEBOCOST metabolite-sensor evidence. Genes were retained when they had direct support from one or more project-data layers. For lipid-handling and cellular-stress modules, canonical genes were retained only when no stronger project-specific alternative was available and were marked as literature-supported fallback genes. The final module definitions and gene-level evidence and provenance tables are provided in Supplementary Data 1.

Module activity was scored at single-cell level using the rank-based decoupler.mt.aucell implementation. Genes absent from the active dataset were omitted and module coverage was recorded. AUCell scoring was run with a minimum target size of three detected genes. Resulting module scores were stored as per-cell score tables and mirrored into the AnnData observation layer for visualization. UMAP overlays were generated with the scomnom plotting API. Compact localization heatmaps were generated by calculating mean module scores within selected clusters and row-scaling each module across the displayed clusters.

### Patient-level module summaries and cross-tissue coupling

For each module, patient-level scores were calculated only within the cell type or cell-type group for which that module was defined. VAT adipocyte modules were summarized within VAT adipocytes, VAT macrophage modules within VAT macrophage populations, hepatocyte modules within liver hepatocyte populations, LSEC modules within liver LSEC populations and myeloid-feedback modules within liver myeloid populations. Patient-module summaries with fewer than 10 cells in the relevant scope were set to missing. Matched VAT and liver summaries were joined by patient and tested across paired samples using Pearson correlation. Spearman correlation was retained as a rank-based sensitivity measure in the output tables. For the primary adipocyte-signalling and hepatocyte-stress association, a partial Pearson correlation was also calculated after residualizing both module scores against the four-level sex-by-MASLD group factor. The partial-correlation P value used n minus the design-matrix rank minus one residual degrees of freedom.

### Cell-cell communication analysis

Within-tissue cell-cell communication analyses used tissue-specific LIANA per-group tables for female and male no-steatosis and MASLD groups.^20,54^

For the liver, route-family panels were restricted to prespecified hepatocyte, LSEC, myeloid and lymphoid source-target routes representing the principal parenchymal, endothelial and immune interactions. The chord panel used edges observed in both female and male MASLD tables with absolute female-minus-male MASLD lr_means difference greater than 0.25.

For VAT, main MASLD route panels merged the female and male MASLD tables by source, target, ligand and receptor, then calculated female-minus-male lr_means differences. Interactions not returned in one group-specific table were assigned an lr_means value of zero for this descriptive comparison. This denotes group-specific detection under the LIANA expression and filtering criteria, rather than evidence that the interaction was biologically absent, and the resulting differences were not treated as formal tests of a sex effect. The chord panel and route overview used selected adipocyte, macrophage, endothelial, fibroblast and mesothelial route families with absolute female-minus-male MASLD lr_means difference of at least 0.25, giving 166 ligand-receptor edges. Supplemental LIANA panels used the same per-group LIANA tables to show edge-level support for adipocyte-macrophage, endothelial-feedback and structural-remodeling routes across no-steatosis and MASLD groups. For each selected edge, group-specific aggregate-rank support and lr_means values were retained in derived tables.

Across liver and VAT within-tissue communication panels, edge color encoded the female-minus-male direction and edge width encoded the absolute lr_means difference. Focused route examples used -log10(aggregate_rank) support for matched ligand-receptor edges.

### Paired liver and VAT framework

The liver and VAT data were analysed as separate annotated single-nucleus RNA-sequencing datasets. A single global liver-VAT embedding was not used for the primary analyses, because organ identity, library chemistry and sequencing depth differed between datasets. Instead, tissue-specific annotations were retained and downstream analyses were performed on patient-level summaries or on focused merged objects that preserved the original tissue and cell-type labels. Absolute nuclei yields were not compared between tissues because library generation used different platforms and nuclei recovery was tissue-specific. Cross-tissue analyses used matched patient-level summaries, so patients rather than individual nuclei were the unit of comparison. Paired analyses used the shared patient identifier present in both tissue metadata tables. Sex and MASLD status were retained as the main stratification variables, with MASLD encoded as no steatosis versus MASLD.

For communication-focused analyses, a derived tissue key was added after merging selected VAT and liver populations. This key was used to restrict communication direction to VAT-to-liver or liver-to-VAT routes where appropriate, while preserving the original source cluster labels.

### Broad patient-level cross-organ ligand-receptor screening

An initial broad cross-organ ligand-receptor analysis was performed at patient level. The broad screen started from a metadata-merged liver-VAT object generated with scomnom adata-ops merge. The reproducible workflow loaded the resulting final scOmnom dataset archive through native scomnom.load_dataset and derived harmonized patient, tissue and organ-cell-type labels from the observation metadata. For each paired patient, liver and VAT cells were subsetted from the merged object, cell types with fewer than 30 cells were removed, counts were normalized to 10,000 counts per cell and log1p transformed, and LIANA rank_aggregate was run with the combined organ-cell-type label as the grouping variable. Only cross-organ ligand-receptor rows were retained. Communication direction was assigned as liver-to-VAT or VAT-to-liver and the two directions were analysed separately.

Secreted and endocrine annotations were derived from OmniPath intercellular annotations, with optional manual curation.^55^ Ligand-receptor pairs were classified as secreted ligand only, secreted ligand and receptor, or membrane/ECM-associated.

Patient-level LIANA results were concatenated across patients. Missing patient-by-axis combinations were assigned a score of zero and flagged as imputed. LIANA scores were re-ranked within each patient’s secreted subset and within direction to reduce score compression after filtering. Axes detected in fewer than 50% of paired patients were excluded from statistical summaries. For each direction, patient-level LIANA scores were evaluated across male no steatosis, male MASLD, female no steatosis and female MASLD groups using prespecified pairwise Welch tests. Benjamini-Hochberg correction was applied within each test family. Chord diagrams display within-sex MASLD-versus-no-steatosis axes with nominal P<0.05 and an absolute patient-level mean rank-score difference of at least 0.10.

### Focused paired ligand-receptor rescoring

Focused paired LIANA rescoring was performed with the scomnom cell-cell communication workflow on a merged object containing biologically plausible VAT sender populations and liver receiver populations. Candidate ligand-receptor pairs for this focused analysis were first enumerated from a pooled cross-tissue LIANA run on the same focused VAT-to-liver object, using rank_aggregate and input_mode = lognorm with VAT as sender tissue and liver as receiver tissue. The pooled screen was used only to define the candidate ligand-receptor universe for rescoring; patient-level effects were then calculated by the paired LIANA backend. Included VAT populations comprised adipocytes, homeostatic macrophages, monocyte-derived macrophages, resident scavenging macrophages and a vascular endothelial population. Included liver populations comprised hepatocyte zones, LSEC populations, liver myeloid populations and vascular endothelial populations. The focused communication analyses used the expression genes retained in the merged object after standard scOmnom filtering; ligand-receptor testing was then limited by database-supported LIANA ligand-receptor pairs and by the specified VAT-to-liver tissue direction.

Because the liver and VAT libraries were generated using different technical platforms, the focused scomnom rescoring analyses used input_mode = lognorm. In this mode, the counts_cb layer was normalized to 10,000 counts per cell and log1p transformed before communication scoring. Paired rescoring used the patient identifier as pairing key, merge_source_cluster_composite as the grouping variable, and sex within MASLD as the primary comparison. The paired backend required at least five sender cells, five receiver cells and three scored patients per group. For each ligand-receptor edge, patient-level communication-potential scores were calculated for each sender-receiver branch and compared between female and male MASLD patients. Route-family summaries were generated by grouping related edges into families such as adiponectin, IGF, HGF, semaphorin, ANGPT, VEGF, GAS and galectin signaling. Edge-level missingness and route-level missingness tables were retained for quality control.

### Metabolite-mediated communication

Metabolite-mediated communication was analysed with MEBOCOST v1.2.2 through the scomnom communication workflow using the same focused VAT-to-liver object and input_mode = lognorm.^22^ Patient-level paired rescoring compared female and male MASLD patients for metabolite-sensor events in the selected sender and receiver populations. The paired backend used the same minimum sender-cell, receiver-cell and scored-patient thresholds as the LIANA paired analysis. Candidate events were summarized by sender branch, receiver branch, metabolite, sensor gene and sensor class. Completeness tables were retained to distinguish unsupported events from events with sufficient patient-level scoring.

### Ligand-target prioritization

Candidate ligands from the focused cross-tissue communication analyses were integrated with NicheNet ligand-target predictions using the scomnom ccc nichenet workflow.^21^ Receiver programs were defined from sex-stratified MASLD liver receiver states. Ligand activity was evaluated using the NicheNet Pearson activity score, and candidate ligands were prioritized when they were both present in paired LIANA ligand-receptor edges toward the corresponding receiver branch and ranked among the higher-scoring NicheNet ligands for that receiver program. This analysis was used as a ligand-prioritization layer linking upstream ligand availability to receiver-state transcriptional programs.

### Hormone-axis, lipid-axis and receiver-gene follow-up

Hormone-axis pathway support was evaluated in the original tissue-specific DE enrichment outputs. Terms related to androgen signaling, estrogen signaling, ESR activity, steroid metabolism, retinoid signaling, vitamin-D signaling, glucocorticoid signaling and nuclear-receptor activity were extracted from Hallmark and Reactome decoupler outputs for the relevant VAT and liver populations. Gene-level follow-up panels summarized log-normalized expression at patient level within the relevant cell population. Dotplot summaries displayed mean expression and the fraction of expressing cells; paired z-score dotplots standardized mean expression within each gene to emphasize sex-associated differences. Heatmaps of gene effects used patient-level standardized effect sizes.

Lipid modules were analysed using the same patient-level module-score framework. VAT adipocyte lipid-metabolism, adipocyte lipid-release and macrophage lipid-scavenging modules were compared with liver hepatocyte lipid-receipt, hepatocyte lipotoxic-stress, LSEC lipid-scavenger and liver myeloid lipid-scavenger modules. Group means were calculated within sex-by-MASLD strata.

### Bulk RNA-sequencing deconvolution and module scoring

Bulk RNA-seq deconvolution was performed separately for liver and VAT using MuSiC as the primary estimator.^56^ The liver deconvolution workflow compared MuSiC, MuSiC2, BayesPrism, DWLS, Bisque and Scaden in a five-fold patient hold-out pseudobulk benchmark with known cell-type proportions.^56–61^ MuSiC had the lowest mean RMSE in the selected liver benchmark input and was therefore used for the main liver and VAT deconvolution analyses. The hold-out pseudobulk benchmark assessed estimator performance where cell proportions were known, but did not reproduce all differences between nuclear and whole-tissue RNA profiles. We therefore interpreted the MuSiC estimates as relative, RNA-informed composition estimates rather than direct cell counts and restricted biological interpretation to broad compartments. Liver and VAT MuSiC proportions were linked to clinical metadata using the curated sample-to-patient identifiers rather than numeric tokens in the library filenames. The metadata audit identified 296 liver and 306 VAT bulk profiles with available MuSiC estimates. Digital HistoIndex steatosis grades were used where available; original NASH CRN grades were retained for samples without a digital score. One VAT-only patient lacked an available grade and was excluded from steatosis analyses. Histological steatosis was encoded as the ordered categories <5%, 5%-33%, 33%-66% and >66%. The complete-case ordinal analyses included 228 females and 68 males in liver and 232 females and 73 males in VAT.

### Sex-stratified trait associations

Sex-stratified ordinal logistic models related ordered steatosis grade to inferred cell fraction while adjusting for age and BMI. Fractions were standardized within tissue, population and sex, and effects are reported as odds ratios per 1-SD higher inferred fraction with 95% confidence intervals. Complete liver and VAT ordinal screens are shown in Extended Data. Sex differences were tested in combined ordinal models containing standardized inferred fraction, sex, their interaction, age and BMI. Fractions were standardized within tissue and population before fitting these combined-sex models. Interaction P values were adjusted across tested compartments within each tissue using the Benjamini-Hochberg method. Sex-stratified Spearman correlations between inferred fractions and steatosis grade were used as rank-based sensitivity analyses.

### Bulk module and gene scoring

Figure 4 module definitions were reused for the bulk RNA-seq module analyses. The main bulk module panels used the liver hepatocyte lipid-receipt, hepatocyte lipotoxic-stress, myeloid-feedback and myeloid lipid-scavenger modules, and the VAT adipocyte signalling, adipocyte lipid-metabolism, adipocyte lipid-release, macrophage activation and macrophage lipid-scavenger modules. Candidate-evidence sender/receiver gene bundles were retained in source tables but omitted from the main panels. Bulk count matrices were restricted to liver and VAT samples with available MuSiC estimates.

Counts were converted to log2(CPM + 1) separately within each tissue. Each gene was centered and scaled across samples within tissue. Module scores were computed as the mean z-scored expression of module genes, followed by within-tissue standardization of each module score. These direct bulk-expression scores were computed independently of the MuSiC estimates. Module-trait models were fit separately by tissue, sex and trait using heteroskedasticity-robust linear models with standardized_module_score ∼ trait + age + BMI. The main module panels display steatosis grade effects. For modules that mapped to a broad MuSiC population, sensitivity models additionally included the corresponding inferred fraction. The unadjusted and population-fraction-adjusted estimates were fitted in the same patients with complete deconvolution data. Matching populations were hepatocytes, Kupffer/myeloid cells, LSECs, adipocytes or macrophages as applicable. Spearman correlations between each core module score and its matching inferred population fraction were calculated across all patients and within sex strata, with Benjamini-Hochberg correction across the tested modules within each stratum. Sex differences in module-trait slopes were tested using the interaction term from standardized_module_score ∼ trait * sex + age + BMI.

A focused gene-anchor panel was selected from the principal single-nucleus-derived modules tested in the bulk analysis. Selection prioritized genes with direct support from the project data and clear biological interpretability; canonical stress or lipid-handling genes were used where needed to provide recognizable anchors for the corresponding module. The panel represented liver hepatocyte stress (*DDIT3*, *HSPA5*, *ATF3*, *HMOX1*), liver myeloid lipid handling (*TREM2*, *LIPA*, *LGALS3*), VAT adipocyte signalling (*ESR2*, *ACVR1C*, *NR3C1*, *JAG1*), VAT macrophage lipid handling (*TREM2*, *LIPA*), and VAT adipocyte lipid context (*ADIPOQ*, *PLIN1*). Gene-level associations used the same standardized logCPM expression scale and the same age/BMI-adjusted sex-stratified steatosis model as the module analyses. A supplemental selected MEBOCOST gene-anchor panel used the same workflow for transcript anchors from the paired metabolite-sensor analysis.

Paired liver-VAT bulk module coupling was tested in 293 patients with both tissues available, including 225 females and 68 males. Pearson and Spearman correlations were calculated across all paired patients and within sex strata for predefined module pairs derived from the single-nucleus cross-tissue analysis. As a sensitivity analysis, both module scores were residualized against age, BMI and ordinal steatosis grade before calculating adjusted Pearson correlations; the adjusted all-patient analysis included 292 complete cases.

Differences in coupling by sex were tested with heteroskedasticity-robust linear models containing the VAT module score, sex, their interaction, age, BMI and ordinal steatosis grade. Interaction P values were corrected across the predefined module pairs. Single-nucleus correlation coefficients were retained as contextual evidence in the aggregate output tables.

Targeted paired gene coupling was tested for MEBOCOST-prioritized VAT sender and liver sensor genes and for matched liver-VAT lipid/scavenger genes. Bulk counts were transformed to log2(CPM + 1), and each gene was centered and scaled within tissue before pairing by patient. Pearson and Spearman correlations were calculated separately in females and males. Complete-case sensitivity analyses residualized both genes against age, BMI and ordinal steatosis grade within sex before calculating adjusted Pearson correlations. Benjamini-Hochberg correction was applied within each gene-pair family and sex.

### Statistical summaries

Patient was used as the statistical unit for paired module, gene-expression and communication analyses. Pearson correlation was used for displayed cross-tissue module-coupling heatmaps and scatterplots, with Spearman correlation retained as a non-parametric sensitivity measure. Patient-level expression and module-score group differences were reported as Hedges’ g, with positive values indicating higher values in female MASLD patients and negative values indicating higher values in male MASLD patients. Focused paired LIANA and MEBOCOST communication-potential comparisons were summarized with Cliff’s delta because the rescored communication values were sparse, bounded and rank-like. Cliff’s delta ranges from -1 to 1, with positive values indicating higher stochastic dominance in female MASLD patients and negative values indicating higher stochastic dominance in male MASLD patients. Mann-Whitney U tests were used for non-parametric group comparisons. Benjamini-Hochberg false-discovery-rate correction was applied within each tested feature family where applicable.^62^

### Software environment

All analyses were performed using custom Python and R scripts in a dedicated scOmnom_env conda environment. Python analyses used Python v3.11.14, scomnom v0.8.1, scanpy v1.11.5, anndata v0.12.7, numpy v2.3.5, pandas v2.3.3, scipy v1.15.3, scikit-learn v1.8.0, statsmodels v0.14.6, numba v0.64.0, networkx v3.6.1, decoupler v2.1.4, gseapy v1.1.12, LIANA v1.7.1, MEBOCOST v1.2.2, matplotlib v3.10.8 and seaborn v0.13.2.^63–65^ R-based plotting and supporting analyses were run with R v4.5.2 and included ggplot2 v4.0.3, circlize v0.4.18 and ComplexHeatmap v2.26.1.^66–68^

### LLM assistance

OpenAI Codex was used to assist with generating portions of the scOmnom codebase and panel-specific analysis scripts. All code, analyses and outputs were curated by the authors.

## Additional information

### Ethics approval and consent

This study was approved by the Amsterdam UMC Medical Ethics Committee (BARIA study, trial registration NL8983). All participants provided written informed consent.

### Data availability

Human single-nucleus data and associated clinical metadata generated and analysed in this study are available from the corresponding author upon reasonable request, subject to institutional approval, applicable data-use agreements and the General Data Protection Regulation (GDPR) as implemented in the Netherlands.

### Code availability

Custom code, workflows, analysis notebooks and figure-generation scripts used for this study are available from the corresponding authors upon reasonable request.

### Author contributions

Mijra Koning collected samples, performed experiments, interpreted the findings, and drafted and revised the manuscript. Iman Man Hu analyzed the data, interpreted the findings, and drafted and revised the manuscript. Artemiy Kovynev analyzed the data, interpreted the findings, and drafted and revised the manuscript. Raymond Landgraaf curated the clinical data, contributed to clinical interpretation, and reviewed and edited the manuscript. Amir H. Alizadeh Bahmani curated the clinical data, contributed to clinical interpretation, and reviewed and edited the manuscript. Daniël Baars curated the clinical data, contributed to clinical interpretation, and reviewed and edited the manuscript. Danijela Vojnović Milutinović provided resources, contributed to interpretation of the findings, and reviewed and edited the manuscript. Nataša Veličković provided resources, contributed to interpretation of the findings, and reviewed and edited the manuscript. Rutger Franken provided resources, contributed to interpretation of the findings, and reviewed and edited the manuscript. Yair Acherman provided resources, contributed to interpretation of the findings, and reviewed and edited the manuscript. Nordin M. Hanssen contributed to study conceptualization and reviewed and edited the manuscript. Hilde Herrema contributed to study conceptualization and reviewed and edited the manuscript. Aleksander Krag provided resources, contributed to interpretation of the findings, and reviewed and edited the manuscript. Fredrik Bäckhed contributed to study conceptualization and reviewed and edited the manuscript. Jacques J.G. Bergman contributed to study conceptualization and reviewed and edited the manuscript. Max Nieuwdorp conceived and designed the study, provided cohort resources, acquired funding, interpreted the findings, and drafted and revised the manuscript. Koen H.M. Prange conceived and designed the study, developed the analytical strategy, analyzed the data, interpreted the findings, and drafted and revised the manuscript. Abraham S. Meijnikman conceived and designed the study, supervised the clinical cohort and sample collection, interpreted the findings, and drafted and revised the manuscript. All authors reviewed and approved the final manuscript and accept responsibility for their contributions.

## Extended Data figure legends

**Extended Data Figure 1.** Quality control, dataset integration and annotation support for Figure 1. Panels are referenced by page and panel label; p1a denotes page 1, panel a. p1a-**f,** Library filtering, retained nuclei and doublet diagnostics for liver and VAT datasets. p2a-**h,** Integration benchmarking and pre/post batch-mixing checks. p3a-**f,** BISC-guided clustering-resolution diagnostics for the root liver and VAT datasets. p4a-**f,** Final sample overlays, batch composition and cluster size distributions. p5a-**d,** Final cluster-level QC and silhouette summaries. p6a-**d,** Liver and VAT supercompartment and compartment UMAPs. p7a,**b,** Marker heatmaps for liver and VAT annotation layers. p8a-**d,** Detailed population UMAPs and marker dotplots.

**Extended Data Figure 2.** Supporting analyses for liver sex differences in MASLD. **a,** No-steatosis pseudobulk sex-DE burden across liver clusters, split by direction. **b,** MASLD-only normalized pseudobulk and cell-level DE support across r4 liver clusters. **c,** Overlap among pseudobulk DE genes higher in female MASLD across liver cell types. **d,** Uncurated top-five female- and male-higher MSigDB decoupler activity terms for the hepatocyte clusters shown in Fig. 2b. **e,** Selected and uncurated top LSEC activity terms. **f,** Selected and uncurated top Kupffer-cell activity terms. **g,** Patient-level female-versus-male MASLD AUCell effects for the seven displayed pathways. Scores were averaged within the indicated population for each patient. Points show standardized effects with 95% confidence intervals; filled points indicate within-tissue FDR<0.05. **h,** Per-cell AUCell pathway-score UMAPs, each shown on its own score scale. **i,** Edge-level LIANA support for LSEC-associated population pairs in MASLD. **j,** Hepatocyte-to-myeloid ligand-receptor pairs supported in both sexes and ranked by the absolute female-minus-male MASLD lr_means difference after deduplication across hepatocyte zones. **k,** Edge-level LIANA support for myeloid-hepatocyte and myeloid-lymphoid population pairs in MASLD. **l,** Patient-level differential-abundance results for pooled-sex and female-versus-male MASLD contrasts, showing CLR- and scCODA-supported populations and their overlap.

**Extended Data Figure 3.** Supporting analyses for visceral-fat sex differences in MASLD. **a,** No-steatosis pseudobulk sex-DE burden across VAT clusters, split by direction. **b,** MASLD-only normalized pseudobulk and cell-level DE support across VAT clusters. **c,** Overlap among pseudobulk DE genes higher in female MASLD in mature adipocytes, homeostatic macrophages and monocyte-derived macrophages. **d,** Uncurated top Hallmark and Reactome decoupler activity terms for mature adipocytes. **e,** Selected and uncurated top MSigDB activity terms in monocyte-derived macrophages. **f,** Selected and uncurated top MSigDB activity terms in homeostatic macrophages. **g,** Patient-level female-versus-male MASLD AUCell effects for the twelve displayed adipocyte pathways. Scores were averaged within mature adipocytes for each patient. Points show standardized effects with 95% confidence intervals; filled points indicate within-tissue FDR<0.05. **h,** Per-cell AUCell pathway-score UMAPs, each shown on its own 5th-99th percentile score scale. **i,** Edge-level LIANA support for adipocyte-macrophage routes in MASLD. **j,** Edge-level LIANA support for endothelial-adipocyte, endothelial-macrophage and macrophage-endothelial routes in MASLD. **k,** Edge-level LIANA support for adipocyte-stromal, adipocyte-mesothelial, stromal-mesothelial and mesothelial-stromal routes in MASLD. **l,** Patient-level differential-abundance results for pooled-sex and female-versus-male MASLD contrasts, showing CLR- and scCODA-supported populations and their overlap.

**Extended Data Figure 4.** Supporting evidence for Figure 4 module localization, focused communication and gene-level effects. **a,** VAT UMAP overlays showing per-cell enrichment of adipocyte-signalling, lipid-metabolism, lipid-release, adipocyte-stress, macrophage-activation and evidence-derived sender-candidate modules. **b,** Liver UMAP overlays showing per-cell enrichment of hepatocyte-stress, lipid-receipt, lipotoxic-stress, myeloid-feedback and evidence-derived receiver-candidate modules. **c,** Full cell-type-scoped module-correlation heatmap across matched VAT-liver pairs; pooled and sex-by-MASLD-group-adjusted statistics are provided in the corresponding source-data table. **d,** UMAP of the selected VAT and liver populations included in the focused communication analysis, with tissue and cell-type labels. **e,** Focused LIANA ligand-receptor dotplot showing exact candidate interactions underlying the route-level effects in Fig. 4c. **f,** LIANA-supported NicheNet ligand-prioritization panel for liver receiver programs. **g-i,** Patient-level gene-effect heatmaps for hormone-axis genes, VAT sender genes and liver receiver genes.

**Extended Data Figure 5.** Supporting bulk RNA-seq validation checks. **a,** Liver pseudobulk benchmark used to select MuSiC as the primary deconvolution method. **b,** Liver broad population composition. The left panel shows mean MuSiC-inferred proportions in bulk RNA-seq; the right panel shows mean observed nuclei proportions across the single-nucleus dataset after aggregation and renormalization to the compartments represented in the MuSiC reference. Error bars indicate 95% confidence intervals across patients. **c,** VAT broad population composition shown using the same inferred-versus-observed layout. **d,** Liver sex-stratified ordinal associations between inferred population fractions and steatosis grade, adjusted for age and BMI. **e,** VAT sex-stratified ordinal associations using the same model. Effects in d and e are odds ratios per 1-SD higher inferred fraction with 95% confidence intervals. **f,** Liver sex-stratified Spearman correlations between inferred population fractions and steatosis grade. **g,** VAT sex-stratified Spearman correlations using the same analysis. **h,** Additional iLAM-like markers tested in sex-stratified steatosis models. **i,** Selected MEBOCOST sender and sensor gene anchors from the paired metabolite-sensor analysis tested in bulk steatosis models. **j,** Complete comparison of unadjusted and matching-population-adjusted story-module steatosis effects. Filled markers show population-fraction-adjusted estimates; open markers show the corresponding unadjusted estimates. **k,** Spearman correlations between each core bulk module score and the MuSiC-inferred fraction of the population in which the module was defined. Dark grey, red and blue points show all patients, females and males, respectively; faded points did not pass FDR correction. **l,** Full paired liver-VAT bulk module correlation heatmap for predefined cross-tissue axes. Cell labels show bulk Pearson r and, where available, the corresponding single-nucleus r. **m,** Paired VAT sender and liver sensor gene coupling for selected MEBOCOST axes in patients with both tissues available. **n,** Paired liver-VAT coupling of matched lipid/scavenger marker genes. Cell labels in panels m and n show bulk Pearson r with FDR stars.

## Supplementary information

**Supplementary Table 1.** Concomitant medication use in the single-nucleus cohort. Medication use is reported for all 42 patients contributing liver, visceral adipose tissue, or both tissues to the final single-nucleus datasets.

**Supplementary Table 2.** Sample availability across the single-nucleus datasets. Summary of tissue-specific dataset and paired cross-tissue sample availability. Across the final post-QC datasets, 42 unique patients contributed at least one tissue. The datasets contained 40 VAT libraries and 34 liver libraries, and 32 patients had matched VAT-liver data.

**Supplementary Table 3.** Clinical characteristics of patients retained in the liver dataset. The table describes the 34 patients whose liver libraries passed quality control and were retained in the final liver single-nucleus dataset. Of these, 32 also had visceral adipose tissue data and entered paired cross-tissue analyses.

**Supplementary Data 1.** Evidence-derived module definitions and gene-level provenance. Workbook containing the VAT and liver module gene memberships used for the paired analyses, with evidence classes and inclusion provenance for differential-expression, pathway, GSEA leading-edge and cell-cell-communication support.

