## Extended Data Figure 1 for "Sex-stratified adipose-liver circuits in human MASLD"

### Extended Data Figure 1 Page 1 (Fig. 1b): Library and filtering QC

Filtering, retained nuclei and doublet diagnostics for the two tissue atlases.

**p1a** Liver filter effects

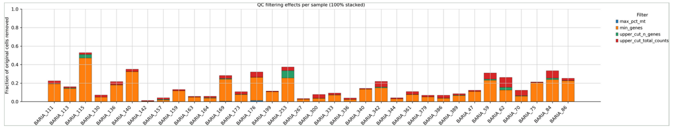

**p1b** VAT filter effects

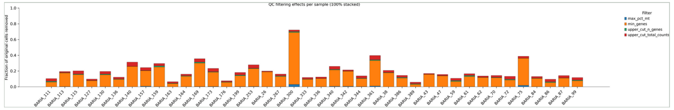

**p1c** Liver retained nuclei

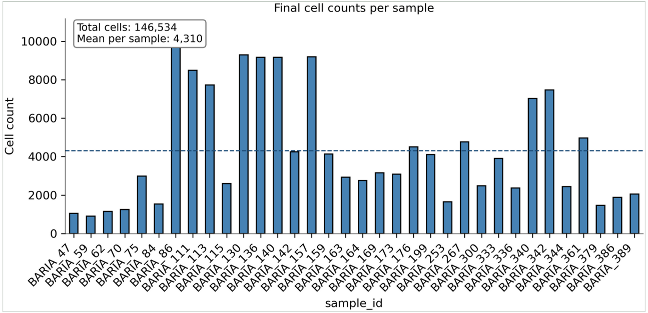


**p1d** VAT retained nuclei

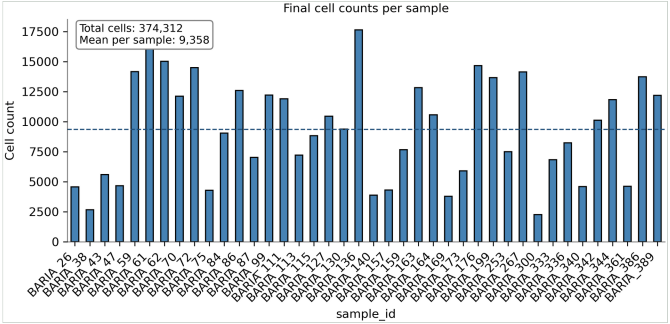


**p1e** Liver doublet score

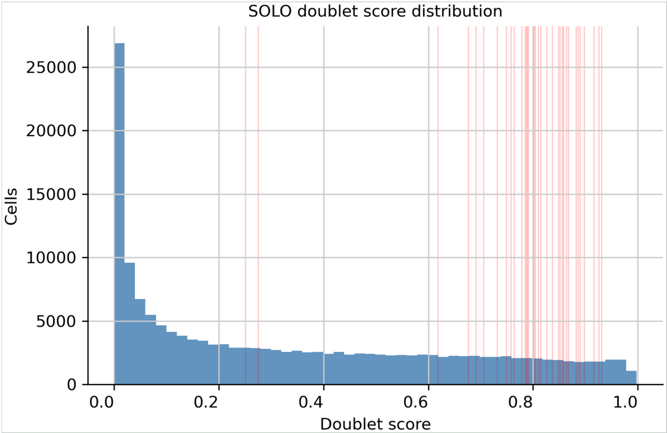


**p1f** VAT doublet fraction

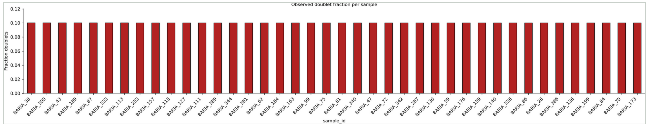

### Extended Data Figure 1 Page 2 (Fig. 1a-c): Integration benchmarking

Candidate integrations and selected pre/post batch-mixing checks.

**p2a** Liver unintegrated

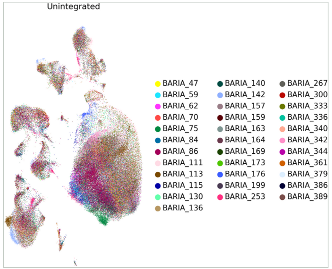


**p2b** Liver Harmony

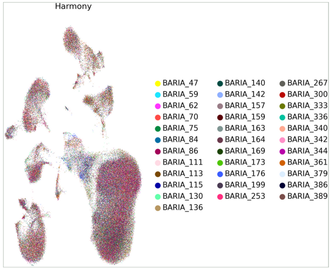


**p2c** Liver scANVI

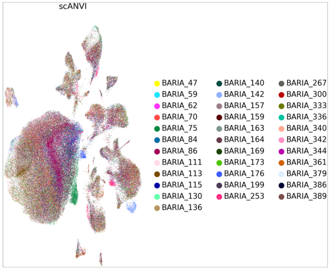


**p2d** Liver pre/post batch

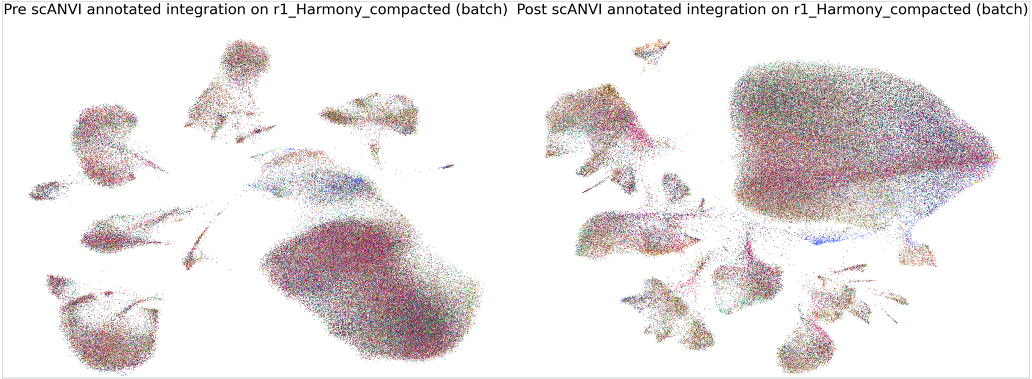


**p2e** VAT unintegrated

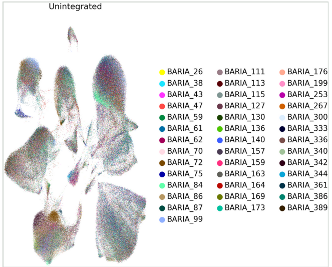


**p2f** VAT Harmony

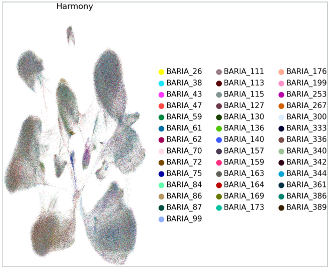


**p2g** VAT scANVI

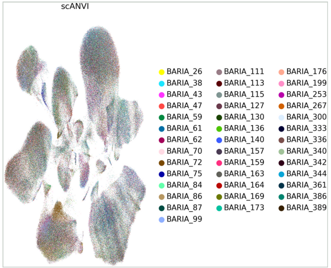


**p2h** VAT pre/post batch

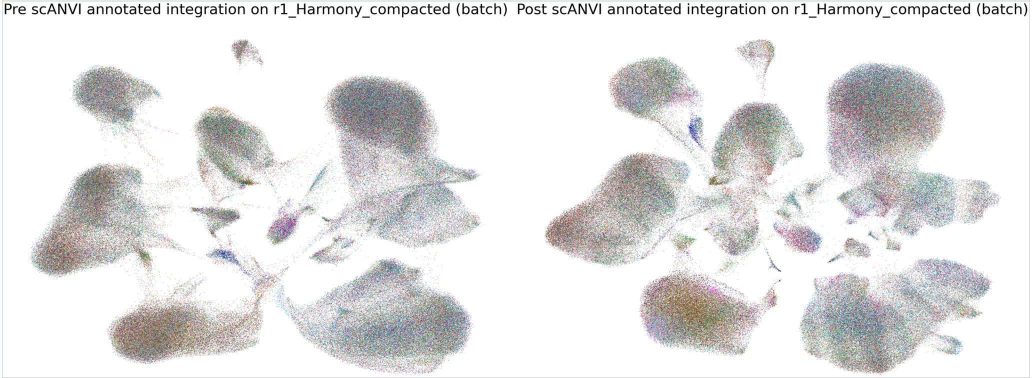

### Extended Data Figure 1 Page 3 (Fig. 1a): BISC-guided clustering resolution

Resolution sweeps and stability diagnostics used to avoid subjective cluster-resolution selection.

**p3a** Liver BISC resolution sweep

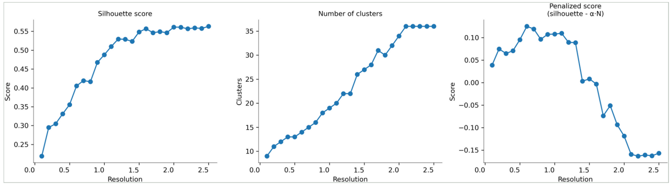


**p3b** Liver selection stability

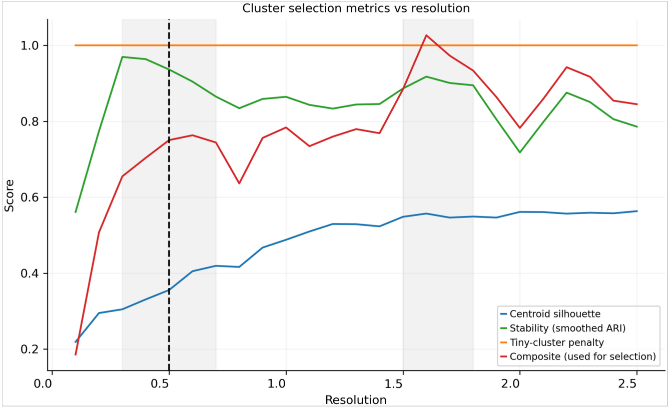


**p3c** Liver ARI stability

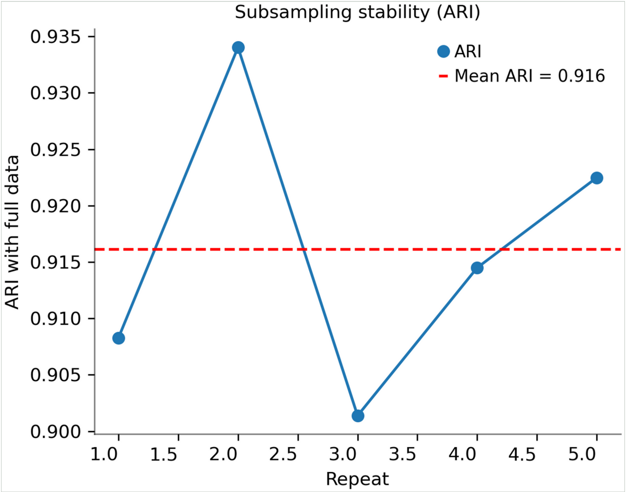


**p3d** VAT BISC resolution sweep

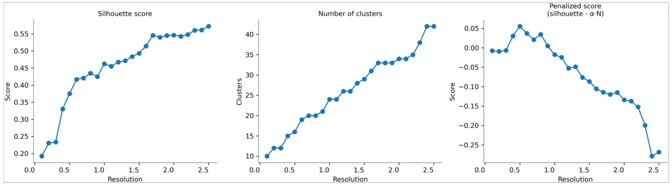


**p3e** VAT selection stability

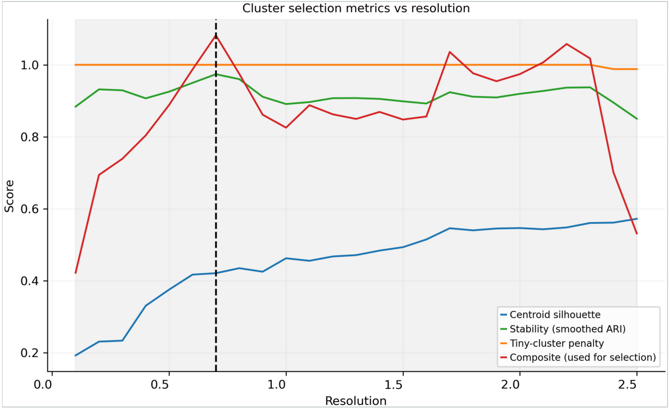


**p3f** VAT ARI stability

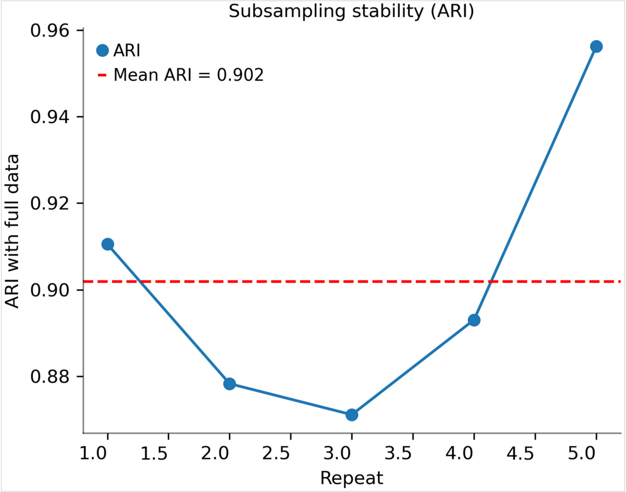

### Extended Data Figure 1 Page 4 (Fig. 1b-e): Final sample and batch composition

Final dataset sample overlays, batch composition and population size checks.

**p4a** Liver sample overlay

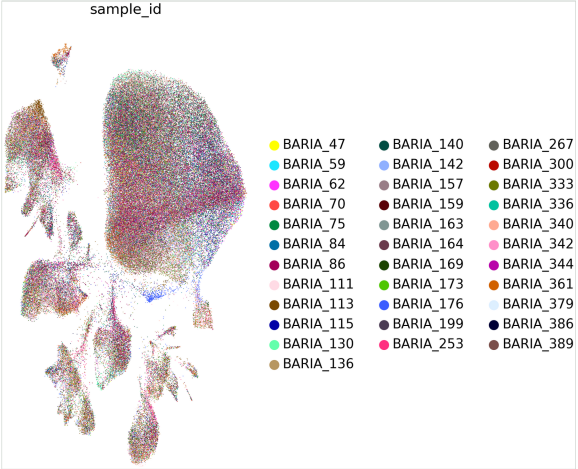


**p4b** Liver batch composition

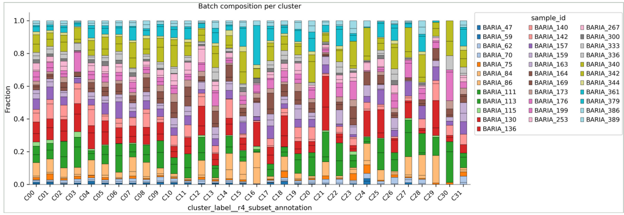


**p4c** Liver cluster sizes

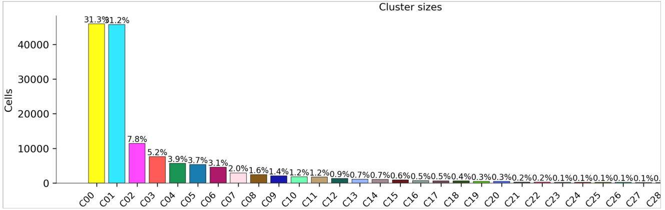


**p4d** VAT sample overlay

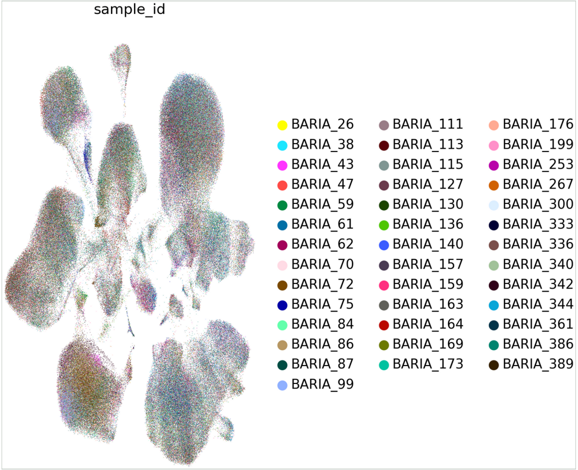


**p4e** VAT batch composition

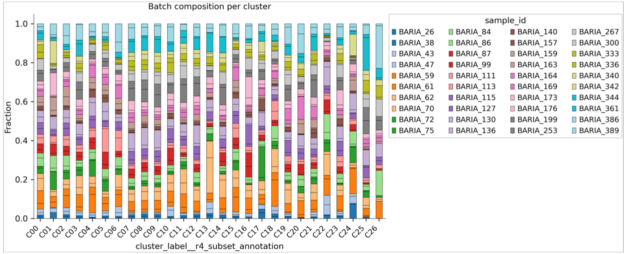


**p4f** VAT cluster sizes

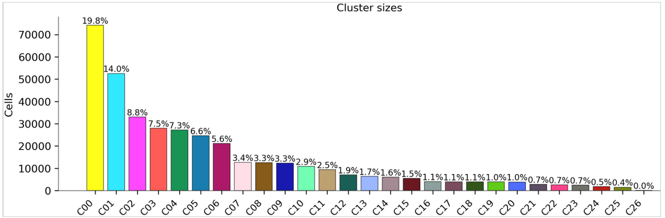

### Extended Data Figure 1 Page 5 (Fig. 1c-d): Final cluster QC

Final cluster-level QC summaries and silhouette support for both atlases.

**p5a** Liver cluster QC

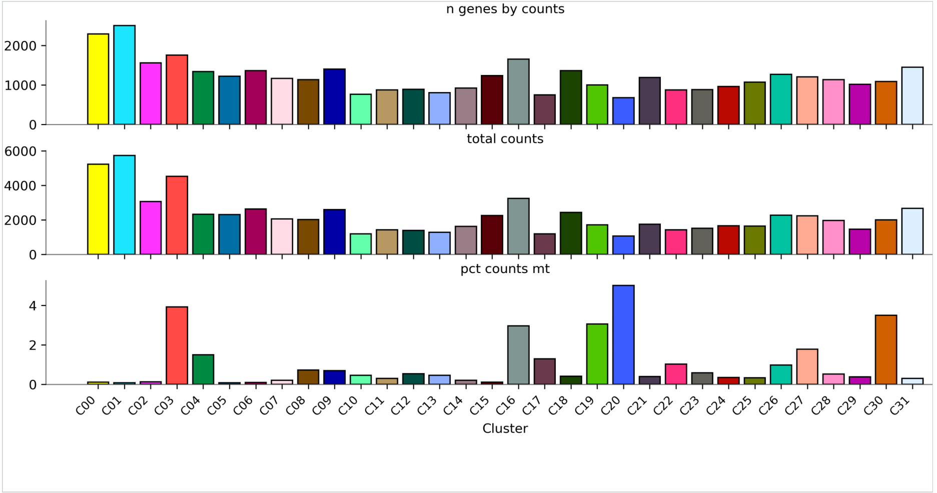


**p5b** Liver silhouette

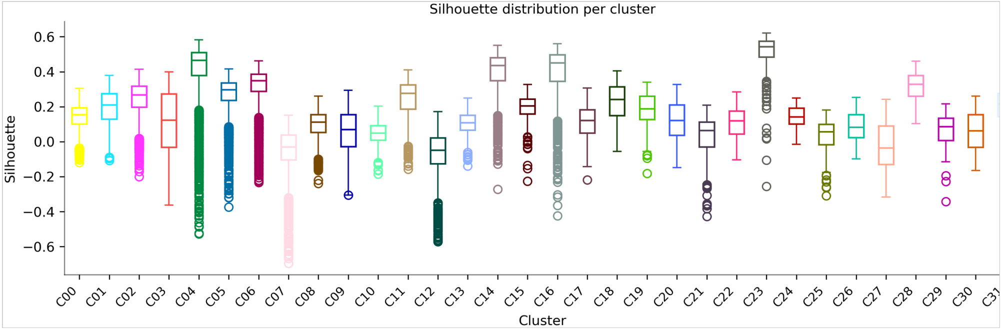


**p5c** VAT cluster QC

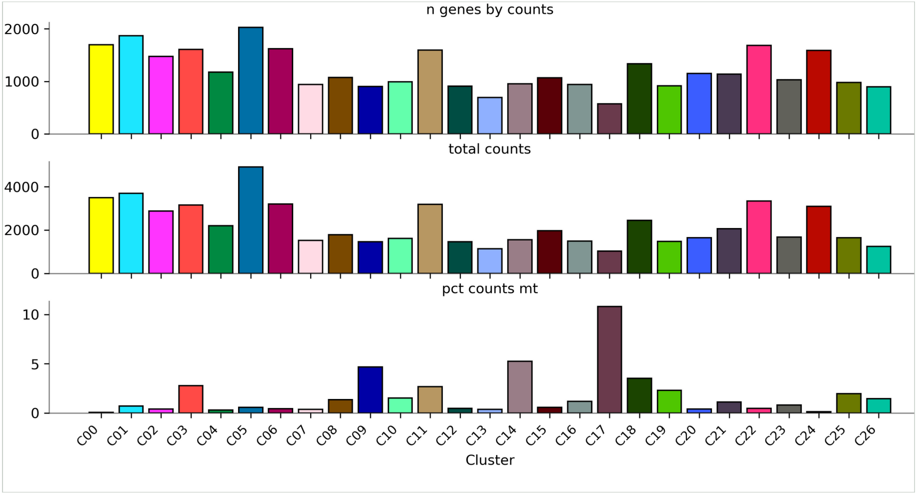


**p5d** VAT silhouette

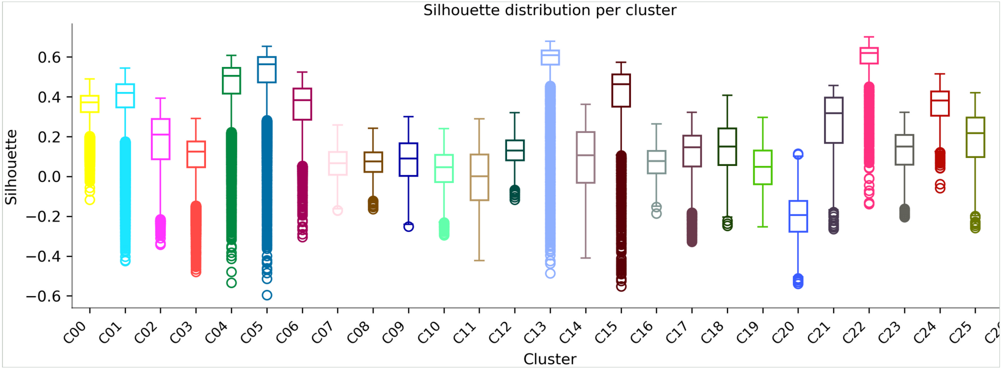

#### Extended Data Figure 1 Page 6 (Fig. 1e): Compartment and supercompartment maps

UMAP locations for the broad composition layers summarized in the main figure.

**p6a** Liver supercompartments

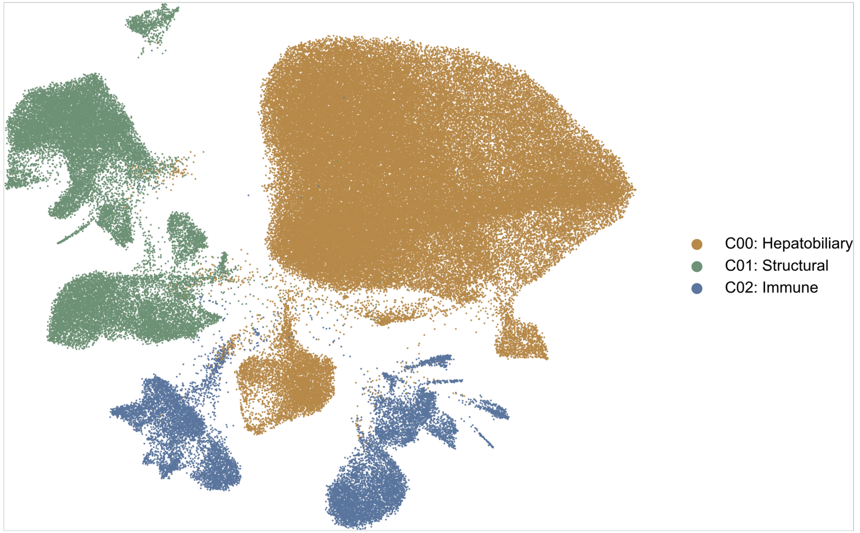


**p6b** Liver compartments

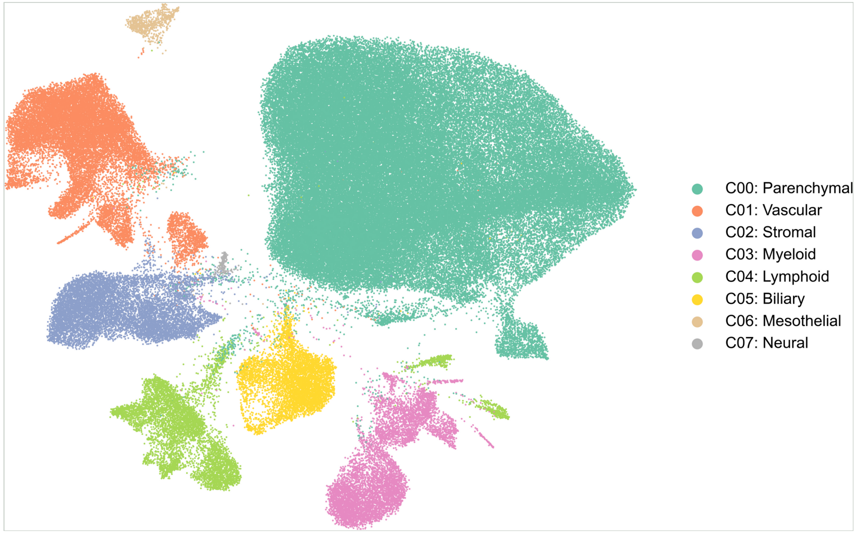


**p6c** VAT supercompartments

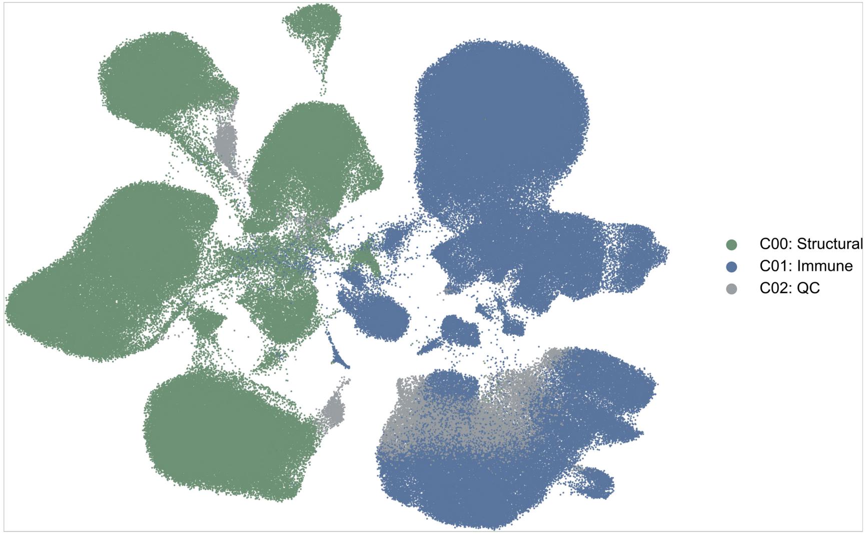


**p6d** VAT compartments

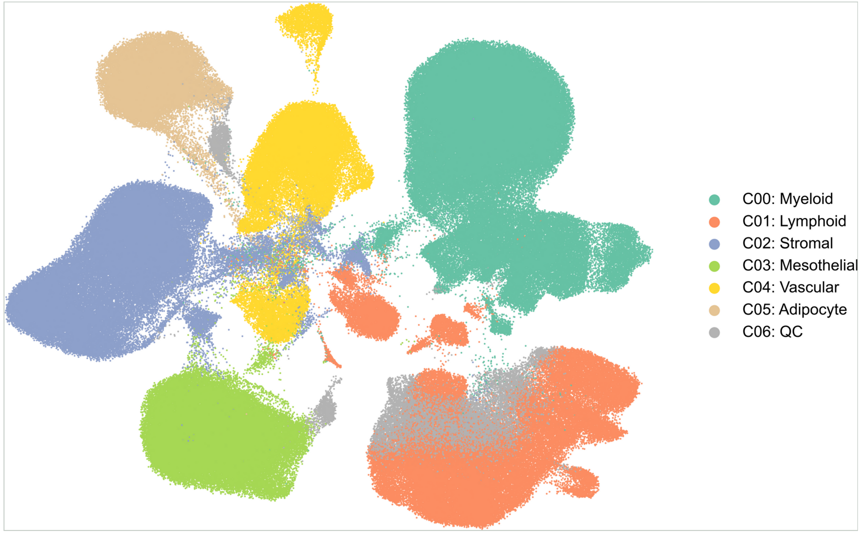

### Extended Data Figure 1 Page 7 (Fig. 1f): Pseudobulk marker heatmaps

Pseudobulk top marker-gene heatmaps from the liver and visceral-fat marker output folders.

**p7a** Liver pseudobulk marker heatmap

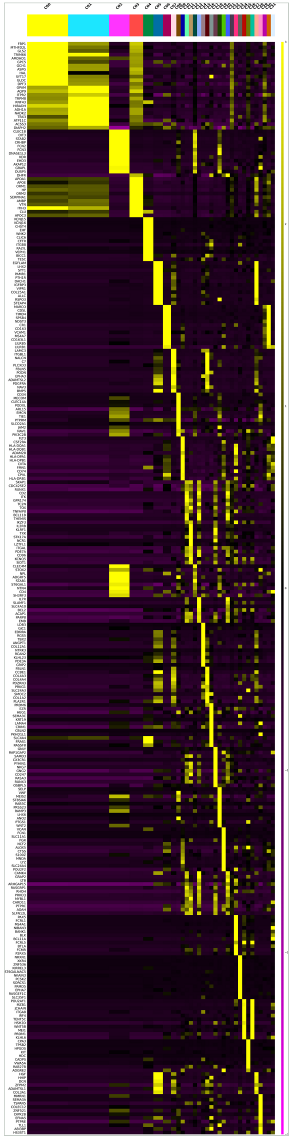

**p7b** VAT pseudobulk marker heatmap

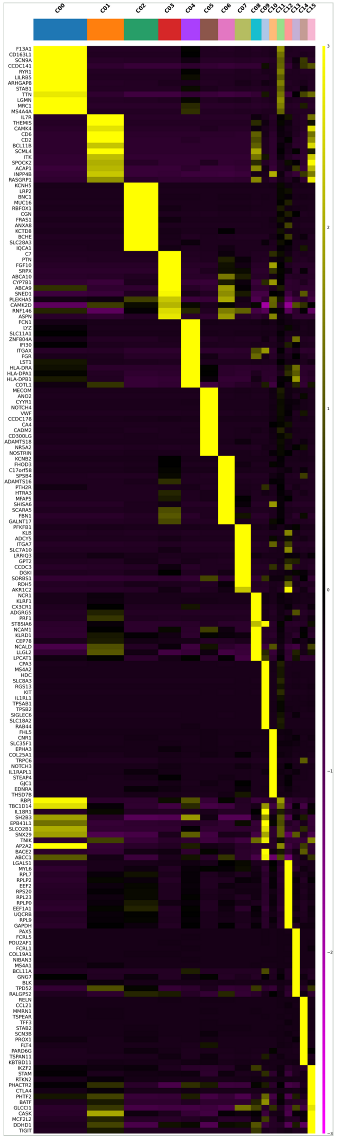

### Extended Data Figure 1 Page 8 (Fig. 1c-d, f): Detailed dataset annotation and marker dotplots

Dense marker checks are kept supplemental so the main figure stays readable.

## p8a

Liver detailed UMAP

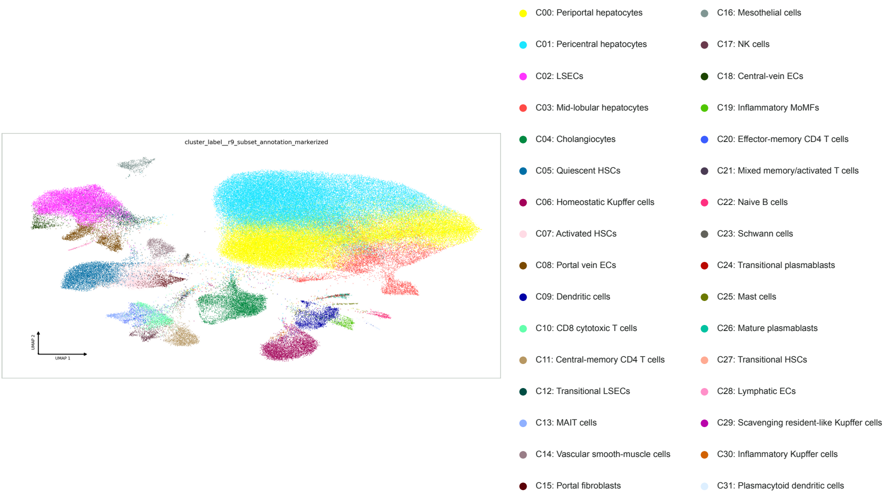


## p8b

Liver detailed marker dotplot

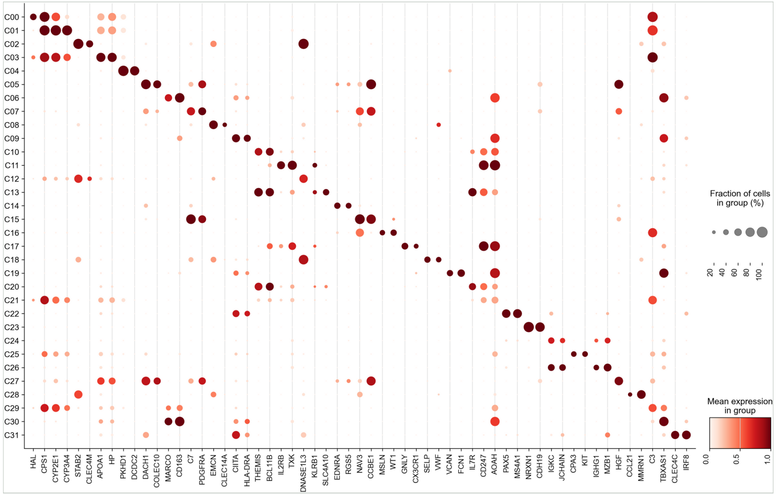


## p8c

VAT detailed UMAP

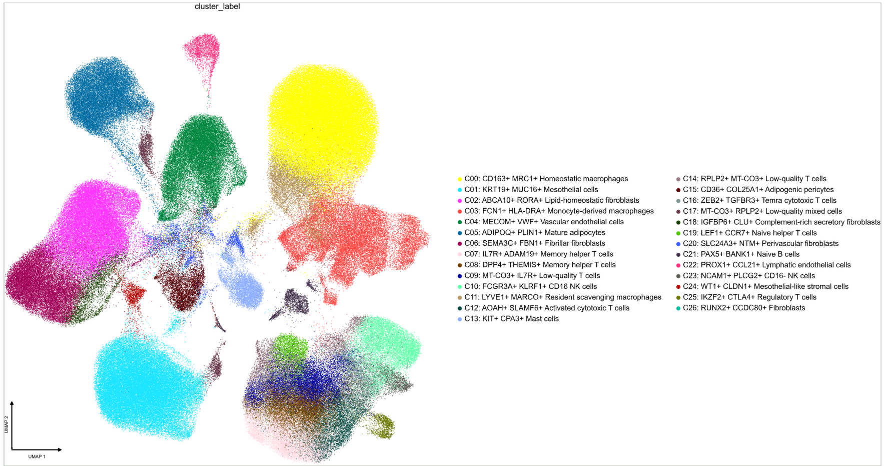


## p8d

VAT detailed marker dotplot

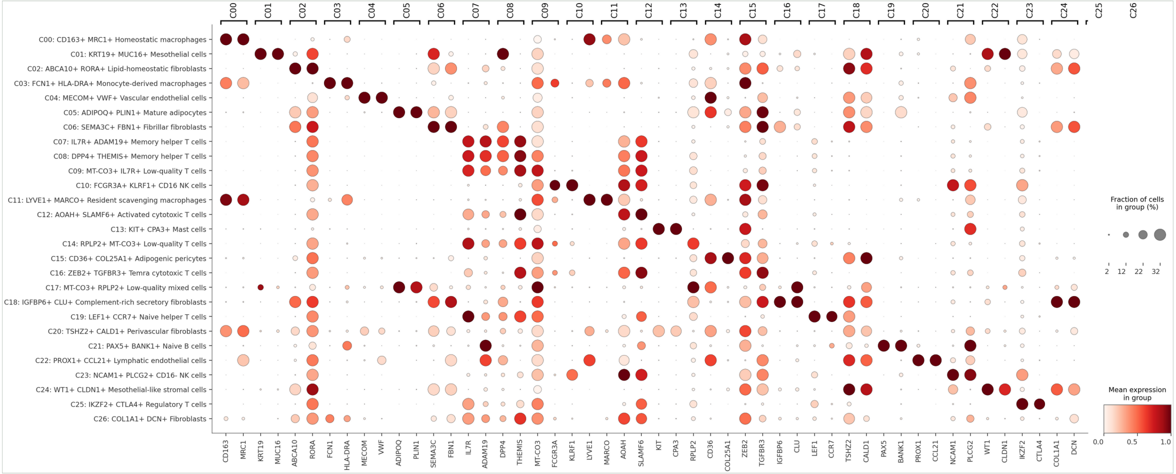
