## Extended Data Figure 2 for "Sex-stratified adipose-liver circuits in human MASLD"

Extended Data Figure 2, page 1

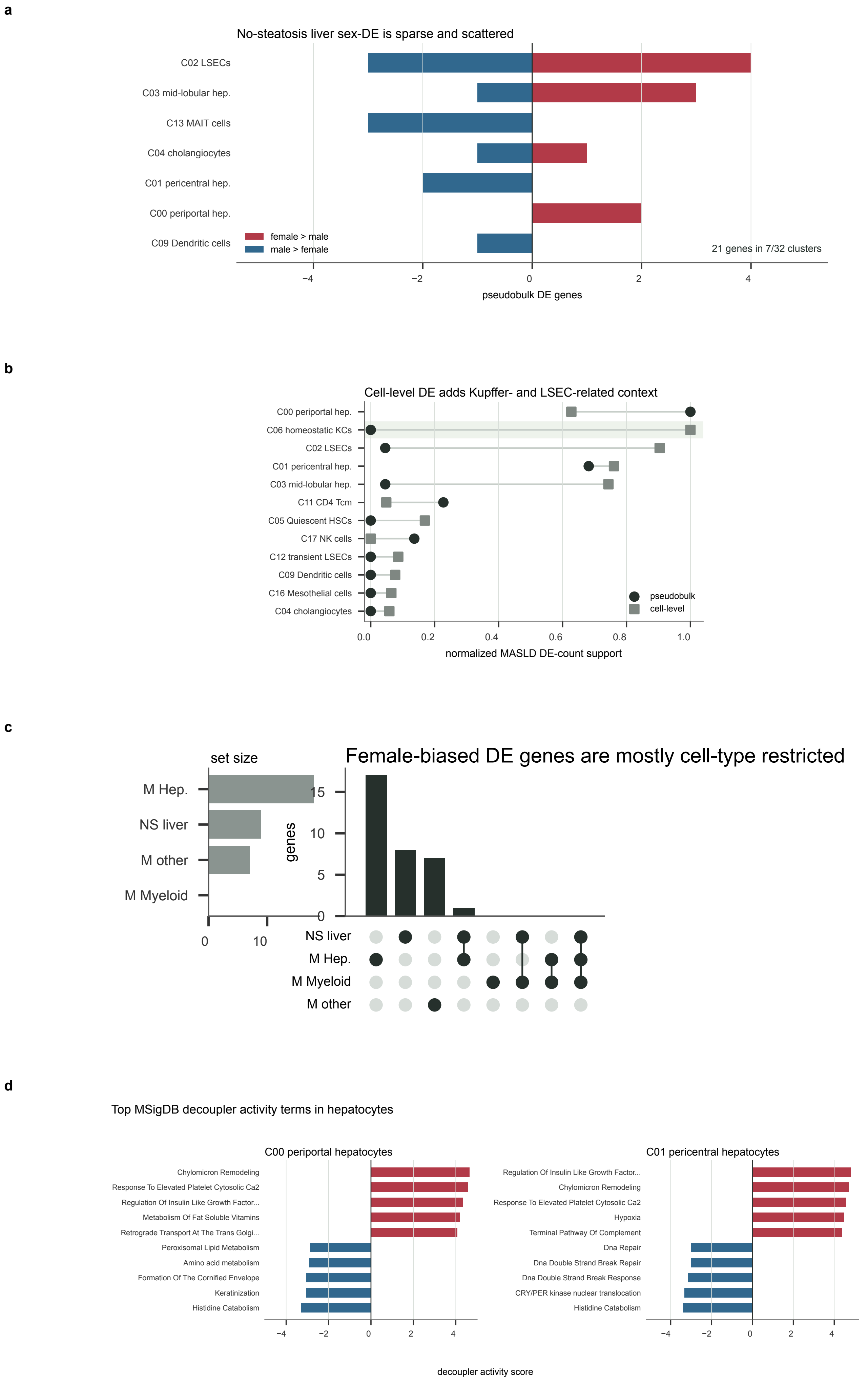

Extended Data Figure 2, page 2

**e**

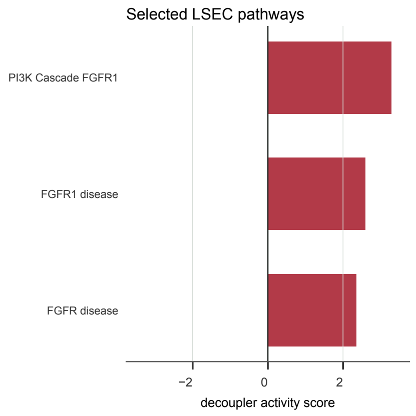


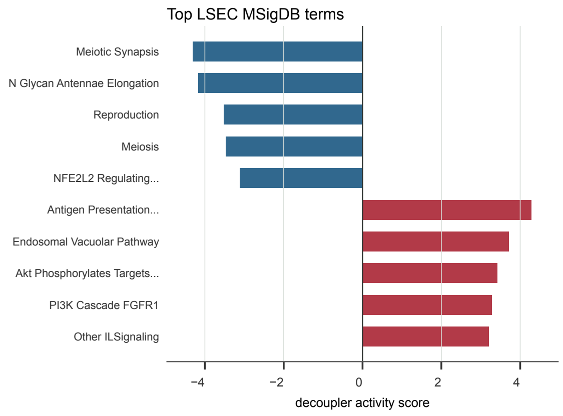


**f**

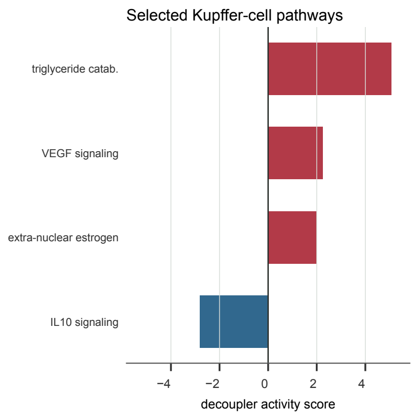


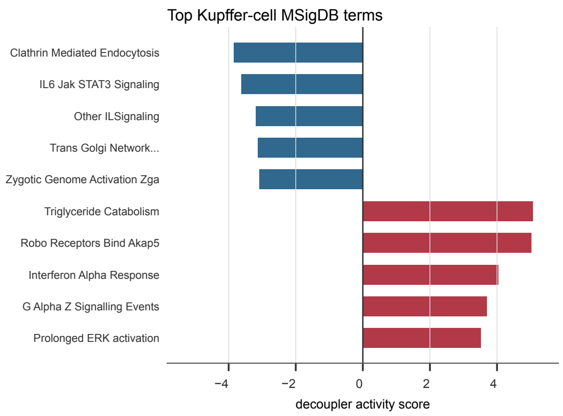


**g**

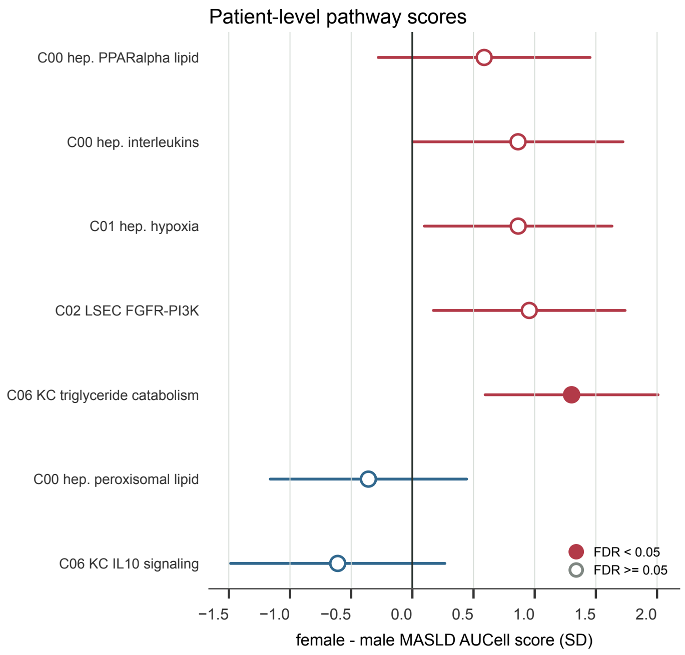


**h**

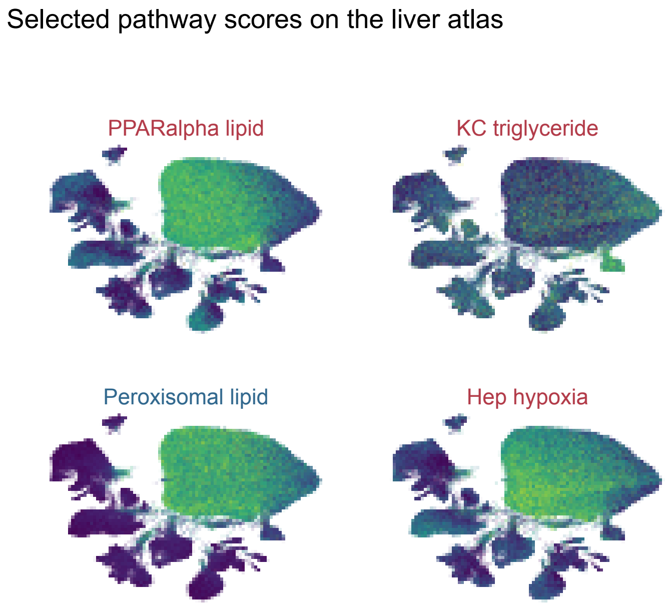

i

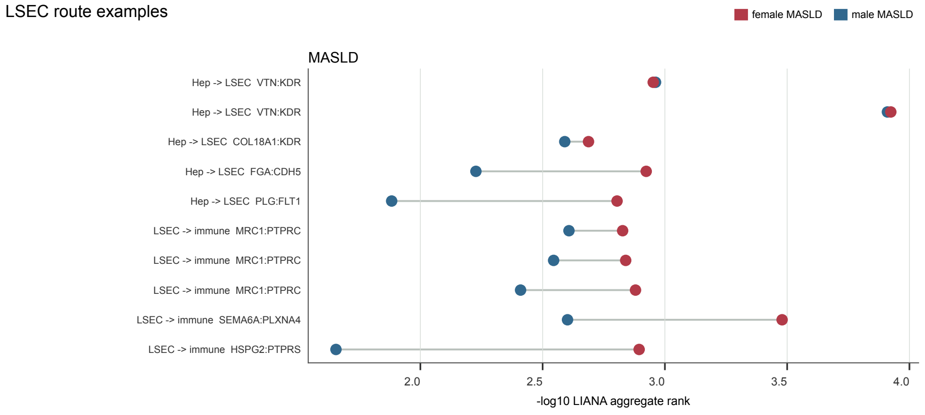


j

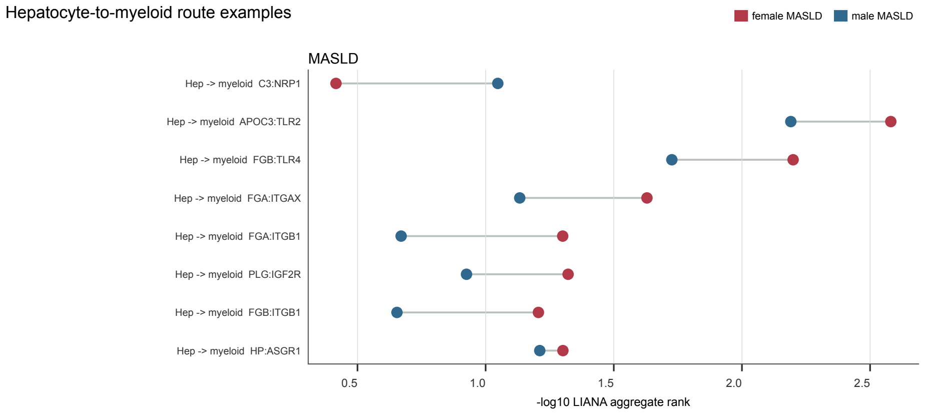


k

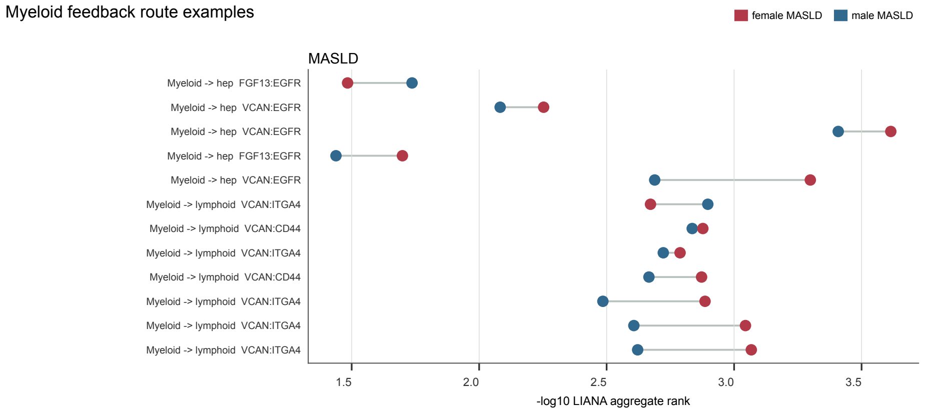


l

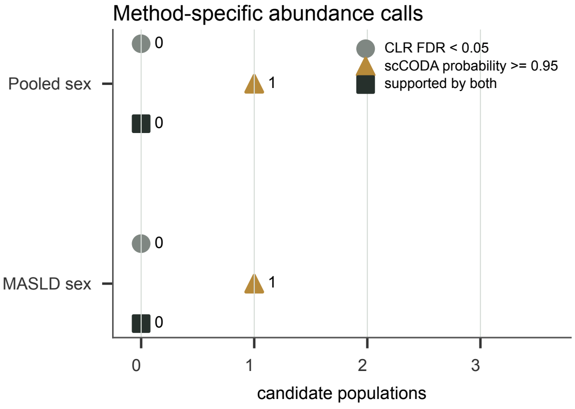


No population was supported by both methods

Pooled sex
CLR: none
scCODA: Mid-lobular hepatocytes (Zone 2) (female higher)

MASLD sex
CLR: none
scCODA: Periportal hepatocytes (Zone 1) (male higher)
