## Extended Data Figure 3 for "Sex-stratified adipose-liver circuits in human MASLD"

**a**

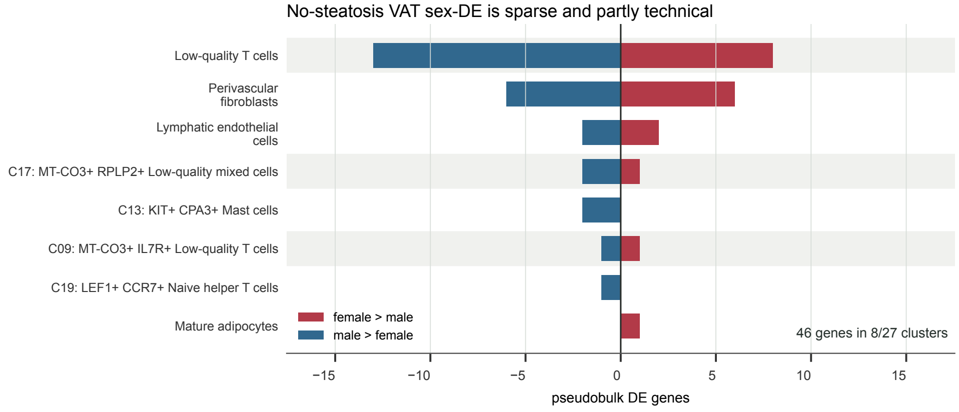
No-steatosis VAT sex-DE is sparse and partly technical
Low-quality T cells
Perivascular fibroblasts
Lymphatic endothelial cells
C17: MT-CO3+ RPLP2+ Low-quality mixed cells
C13: KIT+ CPA3+ Mast cells
C09: MT-CO3+ IL7R+ Low-quality T cells
C19: LEF1+ CCR7+ Naive helper T cells
Mature adipocytes
female > male
male > female
46 genes in 8/27 clusters
−15
−10
−5
0
5
10
15
pseudobulk DE genes

**b**

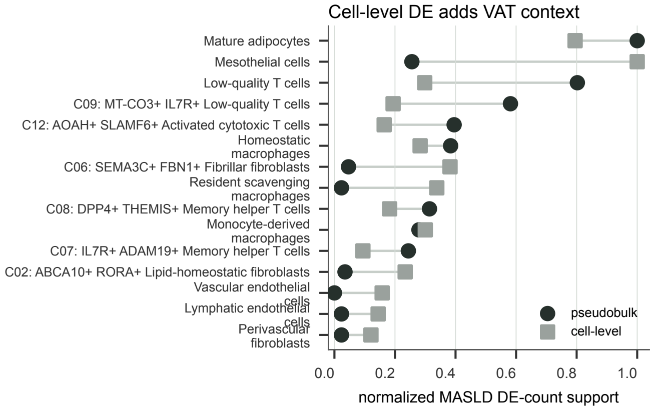
Cell-level DE adds VAT context
Mature adipocytes
Mesothelial cells
Low-quality T cells
C09: MT-CO3+ IL7R+ Low-quality T cells
C12: AOAH+ SLAMF6+ Activated cytotoxic T cells
Homeostatic macrophages
C06: SEMA3C+ FBN1+ Fibrillar fibroblasts
Resident scavenging macrophages
C08: DPP4+ THEMIS+ Memory helper T cells
Monocyte-derived macrophages
C07: IL7R+ ADAM19+ Memory helper T cells
C02: ABCA10+ RORA+ Lipid-homeostatic fibroblasts
Vascular endothelial cells
Lymphatic endothelial cells
Perivascular fibroblasts
pseudobulk
cell-level
0.0
0.2
0.4
0.6
0.8
1.0
normalized MASLD DE-count support

**c**

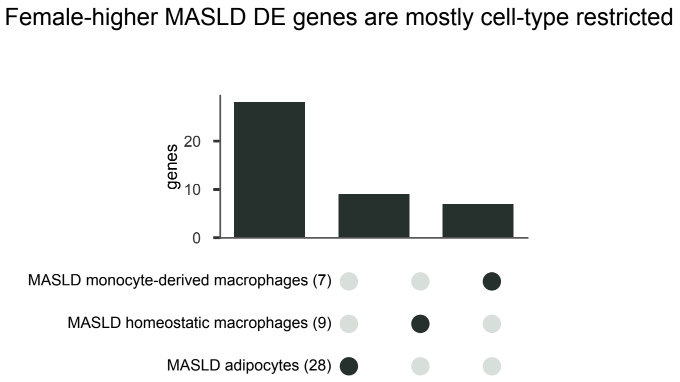
Female-higher MASLD DE genes are mostly cell-type restricted
genes
0
10
20
MASLD monocyte-derived macrophages (7)
MASLD homeostatic macrophages (9)
MASLD adipocytes (28)

**d**

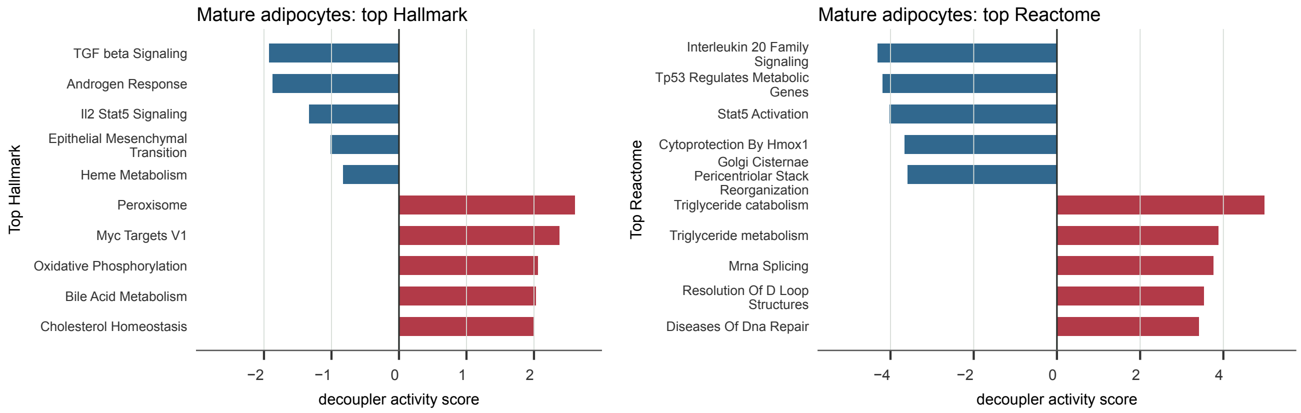
Mature adipocytes: top Hallmark
Top Hallmark
TGF beta Signaling
Androgen Response
Il2 Stat5 Signaling
Epithelial Mesenchymal Transition
Heme Metabolism
Peroxisome
Myc Targets V1
Oxidative Phosphorylation
Bile Acid Metabolism
Cholesterol Homeostasis
−2
−1
0
1
2
decoupler activity score
Mature adipocytes: top Reactome
Top Reactome
Interleukin 20 Family Signaling
Tp53 Regulates Metabolic Genes
Stat5 Activation
Cytoprotection By Hmox1
Golgi Cisternae Pericentriolar Stack Reorganization
Triglyceride catabolism
Triglyceride metabolism
Mrna Splicing
Resolution Of D Loop Structures
Diseases Of Dna Repair
−4
−2
0
2
4
decoupler activity score

Extended Data Figure 3, page 2

**e**

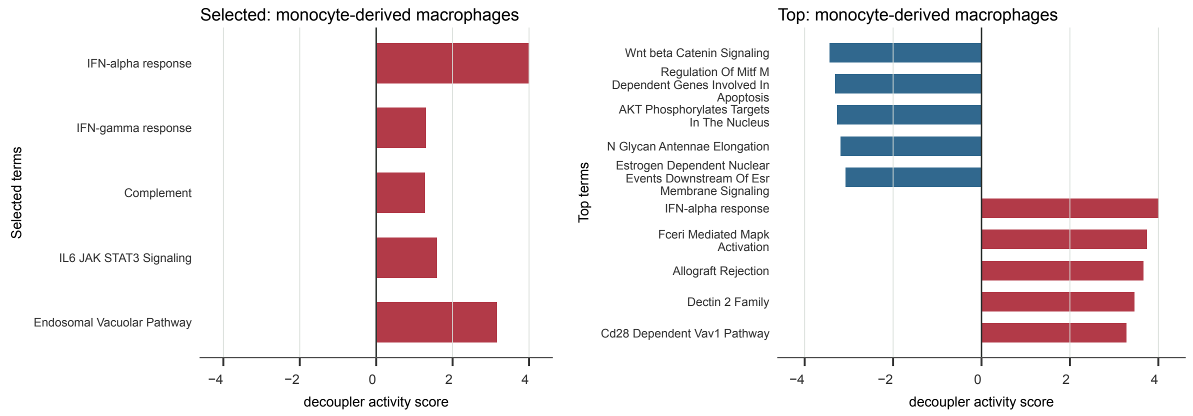


**f**

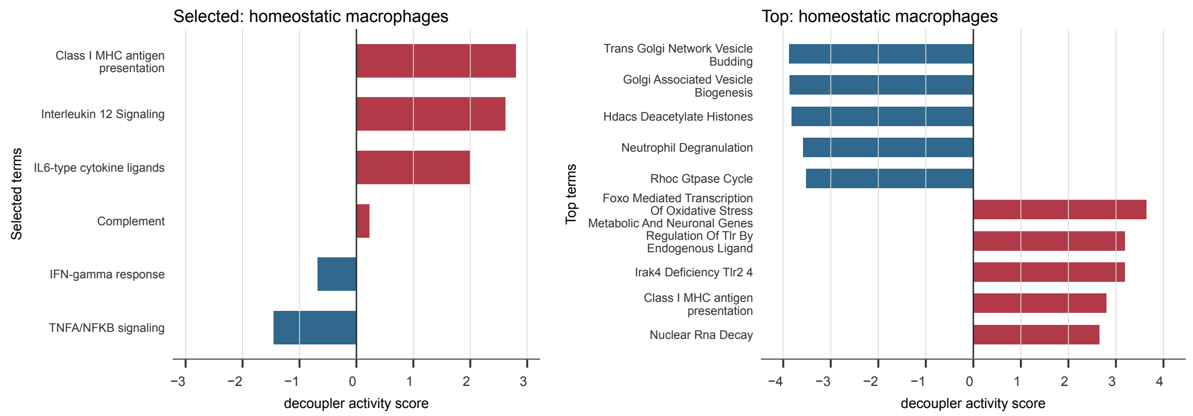


**g**

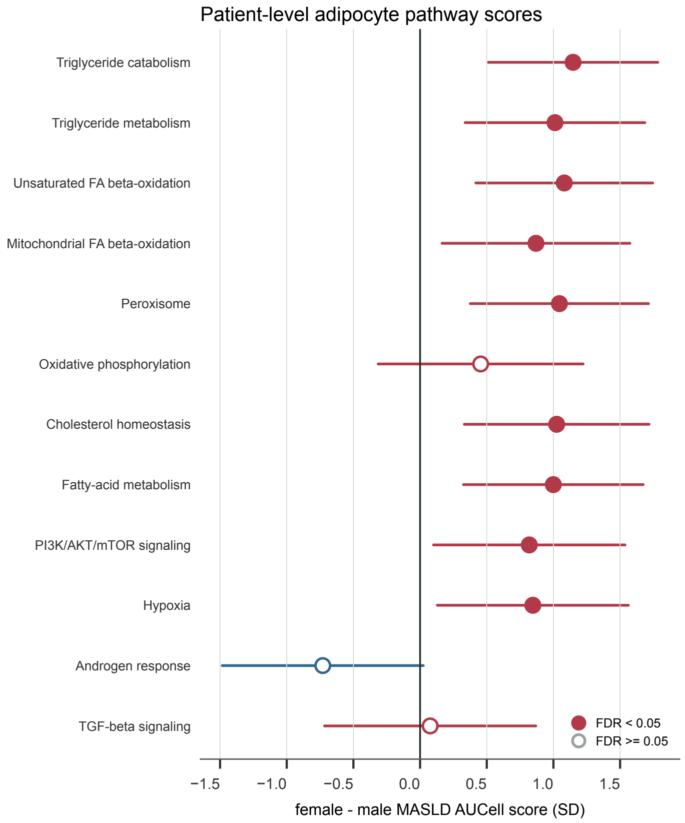


**h**

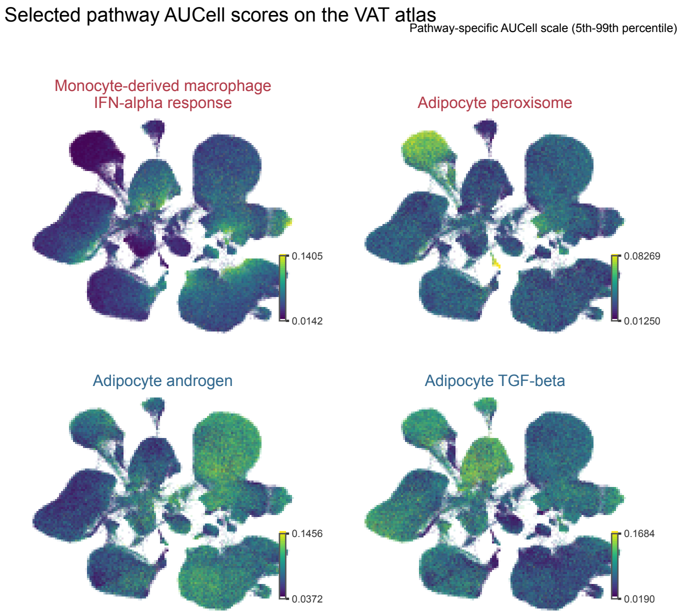

**i**

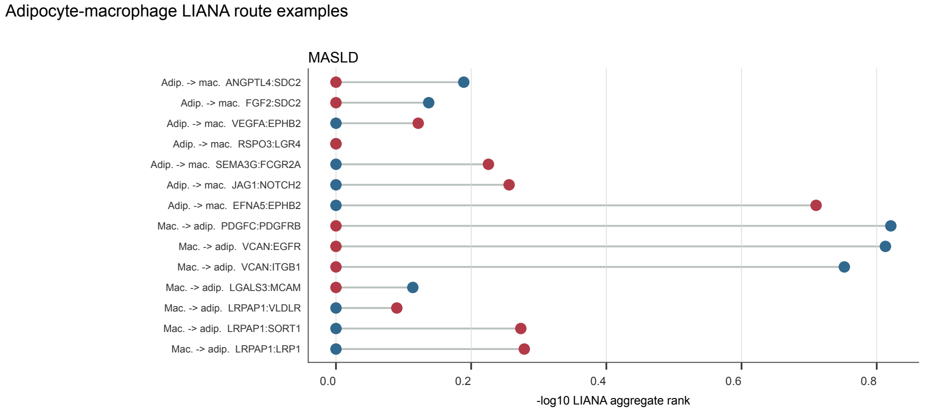


**j**

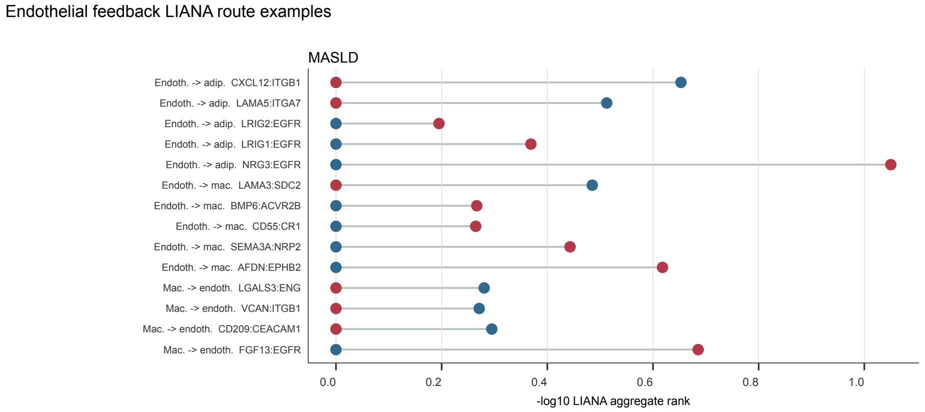


**k**

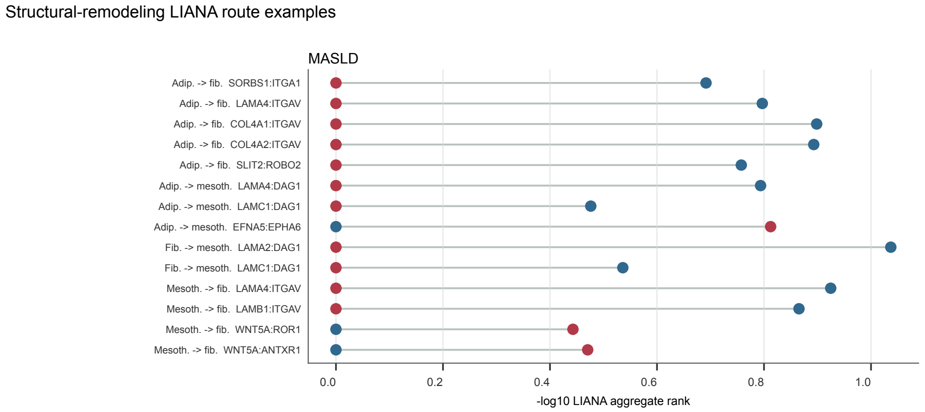


**l**

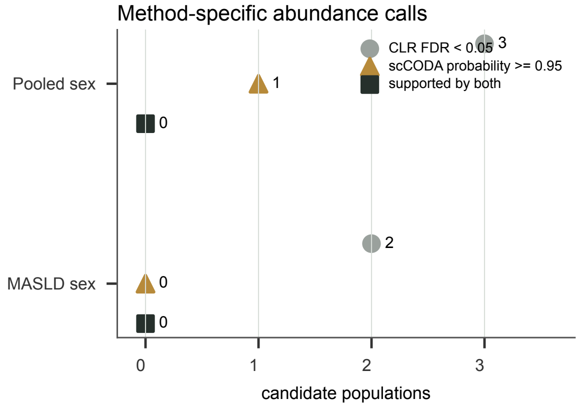


No population was supported by both methods

Pooled sex
CLR: Regulatory T cells (male higher); Lymphatic endothelial cells (female higher); Fibroblasts (male higher)
scCODA: Homeostatic macrophages (female higher)

MASLD sex
CLR: Regulatory T cells (male higher); Resident scavenging macrophages (male higher)
scCODA: none
