## Supplementary figures and images for "Sex-stratified adipose-liver circuits in human MASLD"

### Extended Data Figure 4

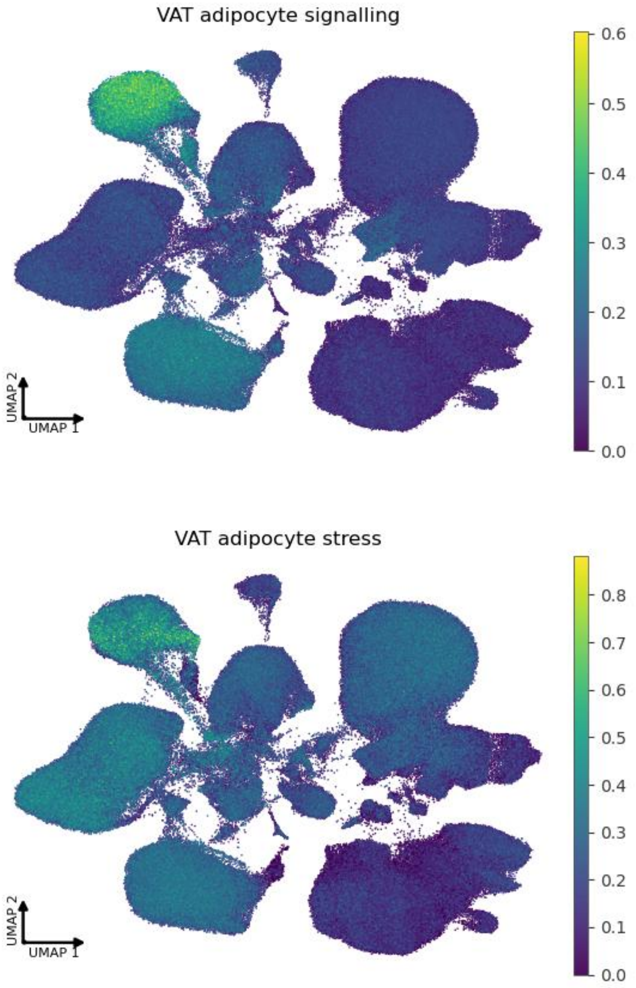

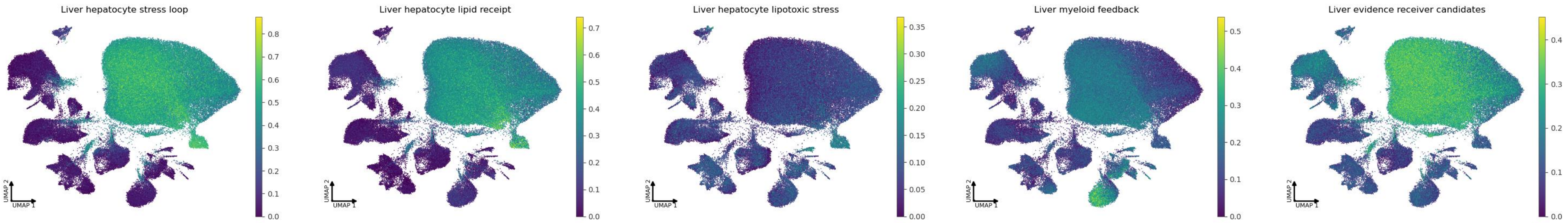

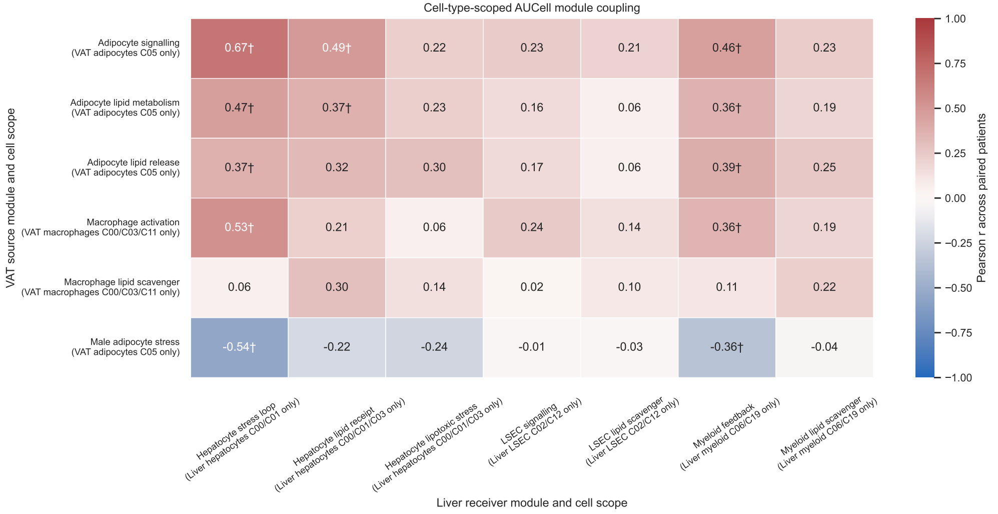

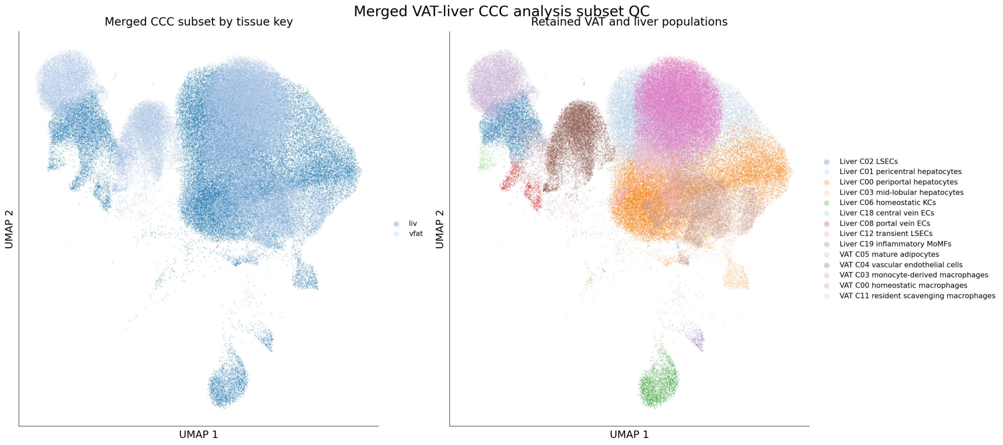

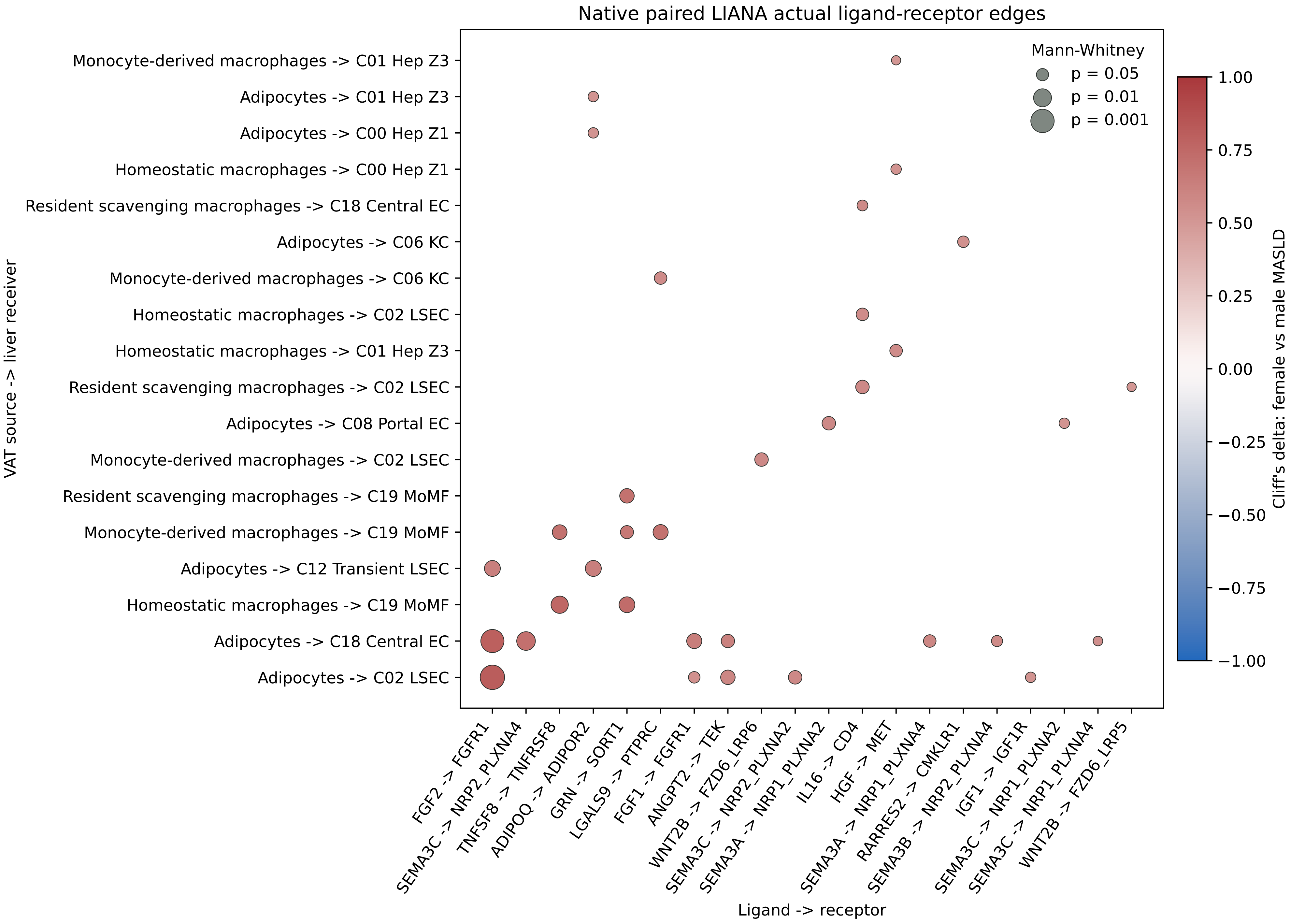

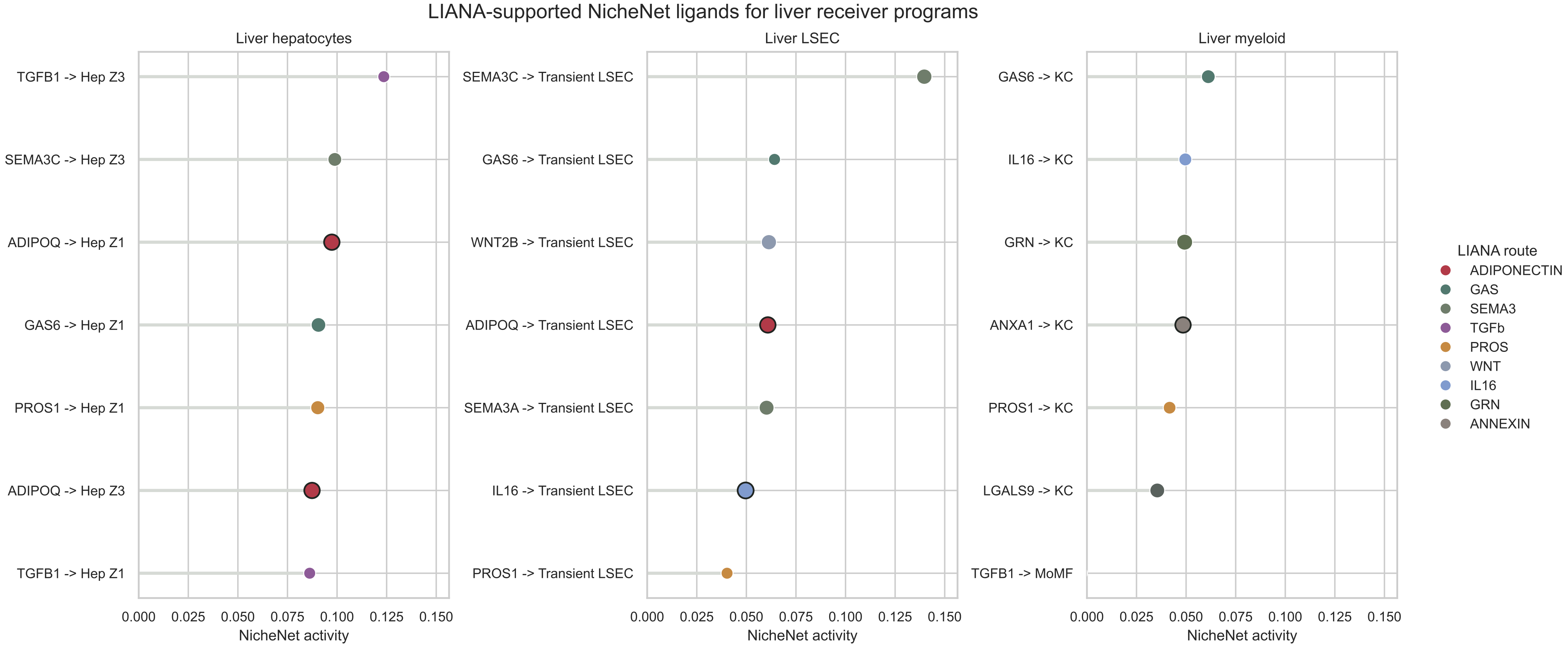

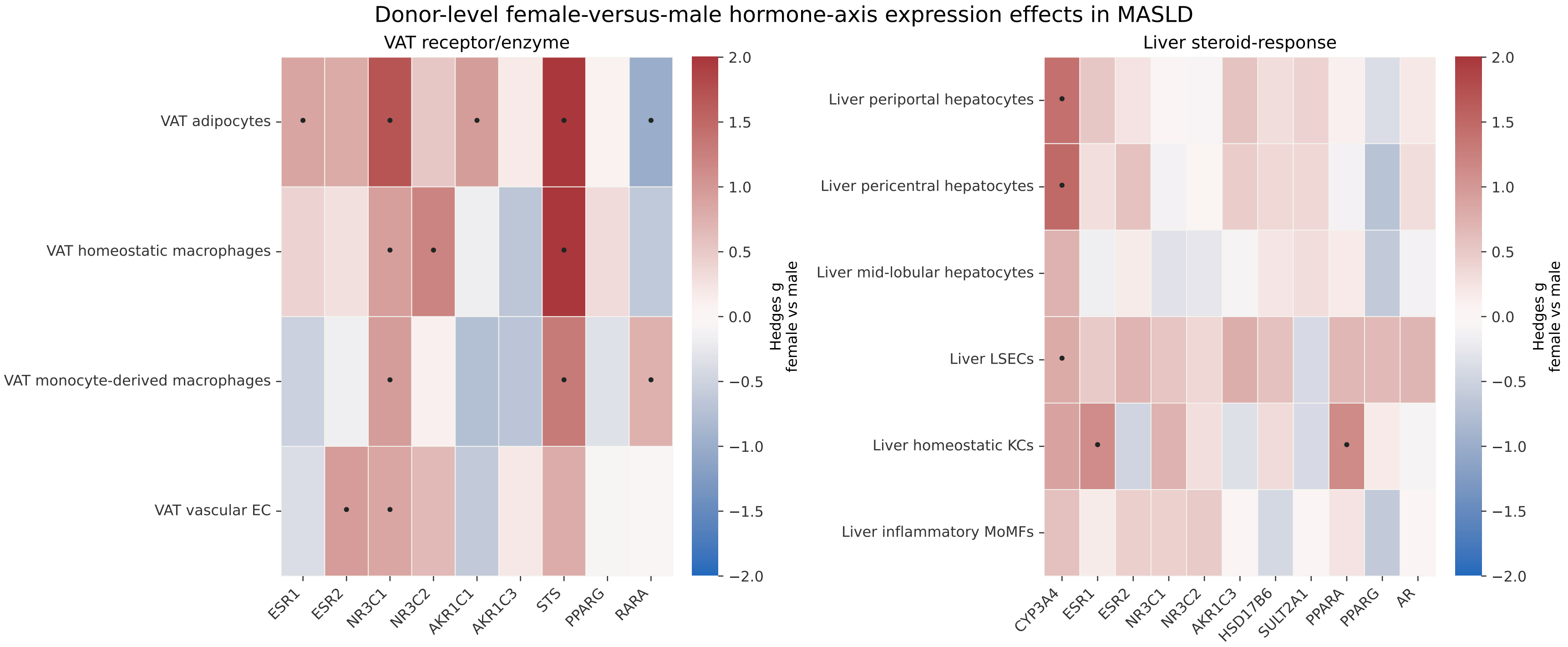

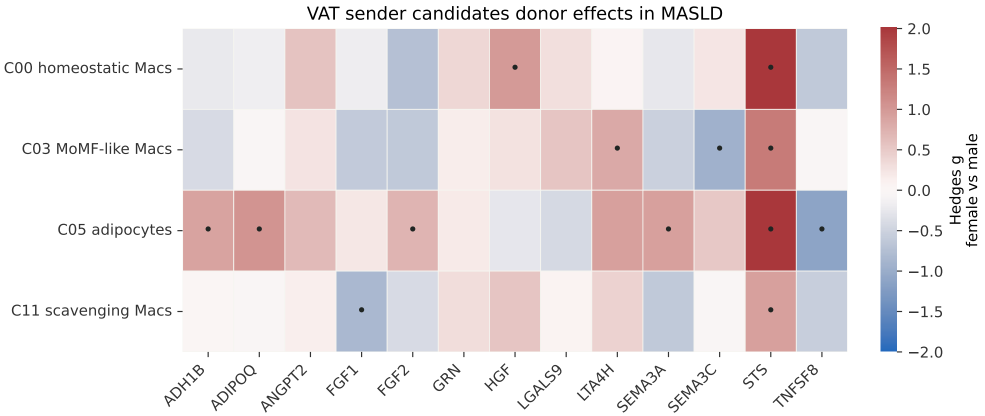

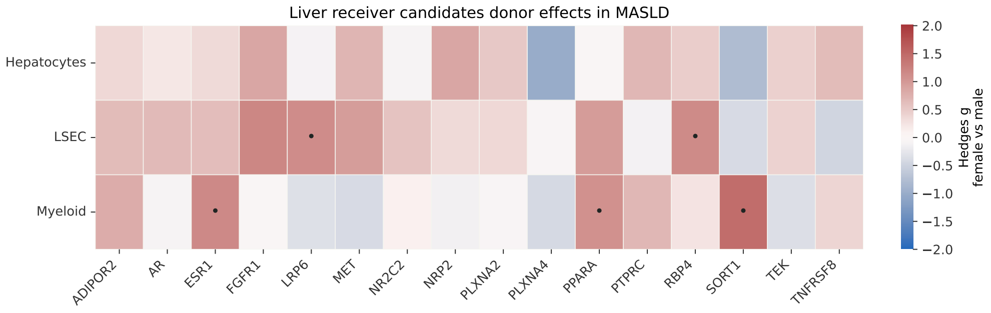

### Extended Data Figure 5

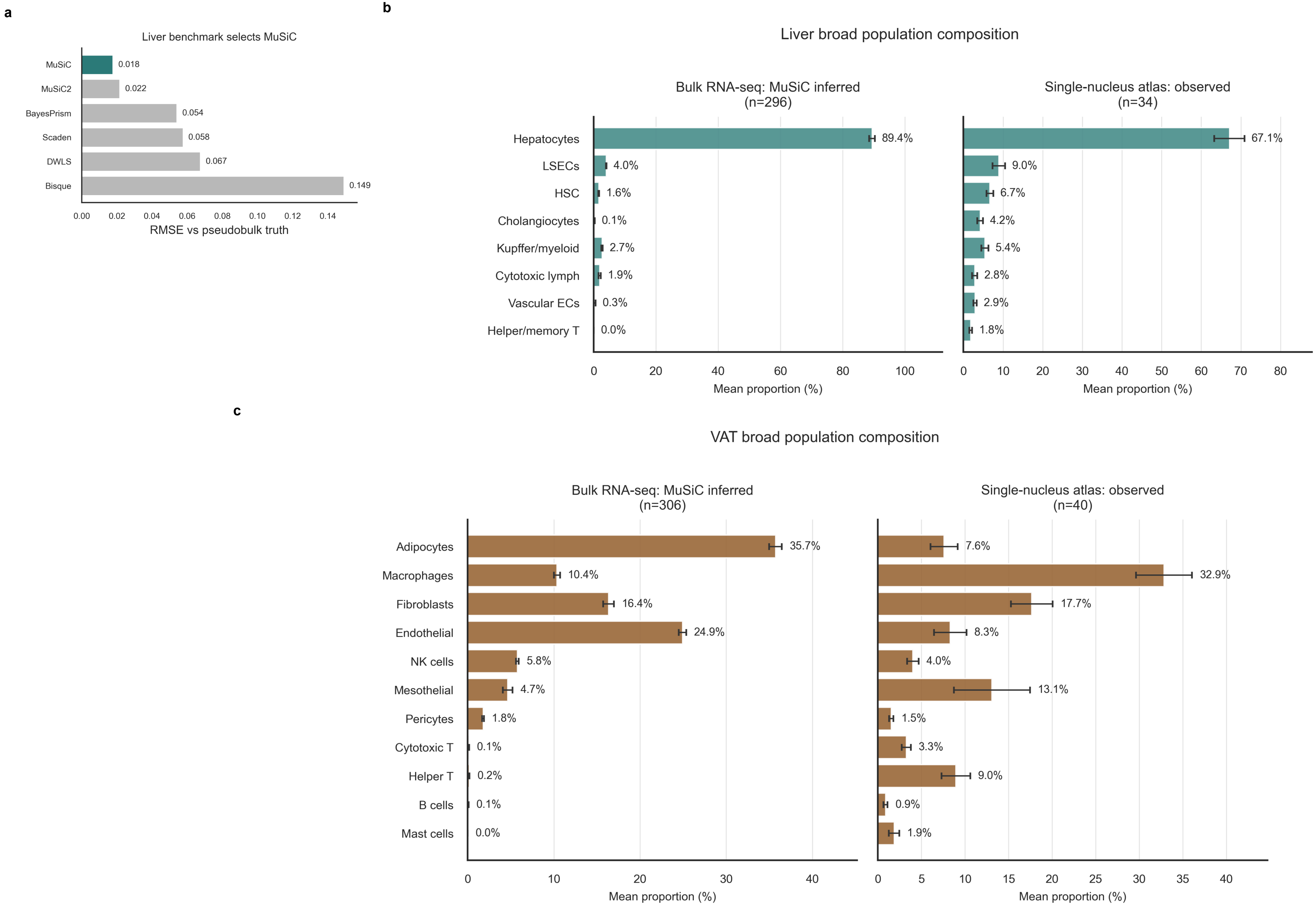

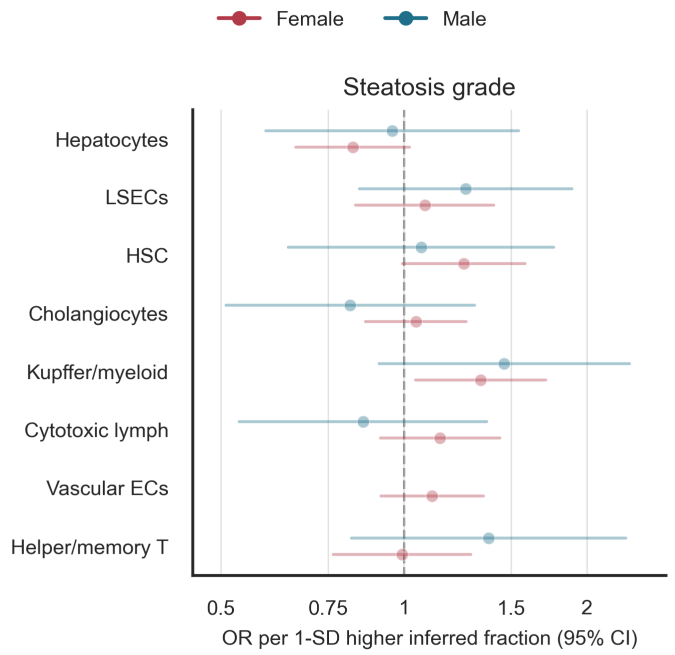

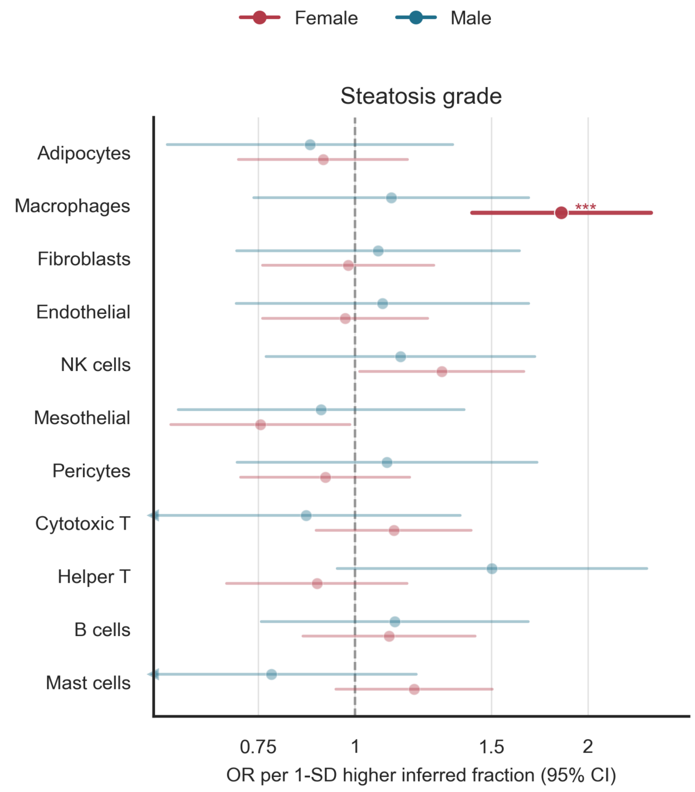

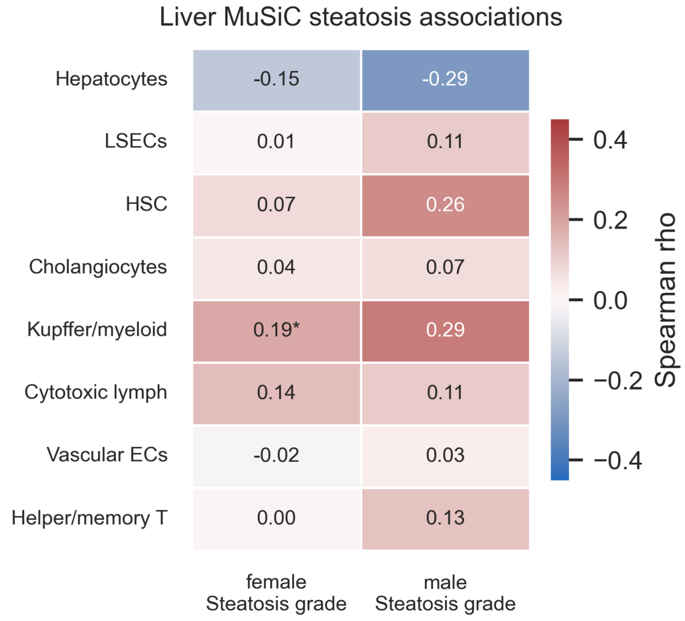

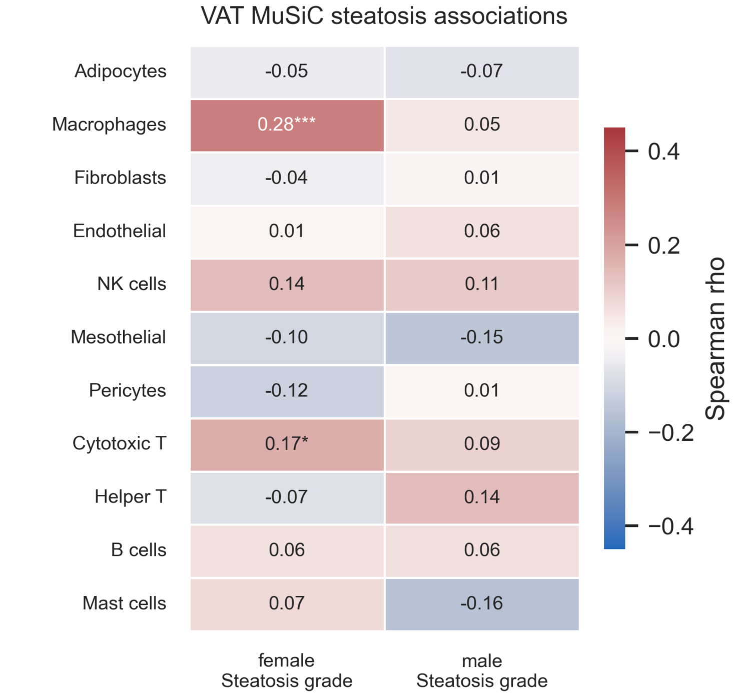

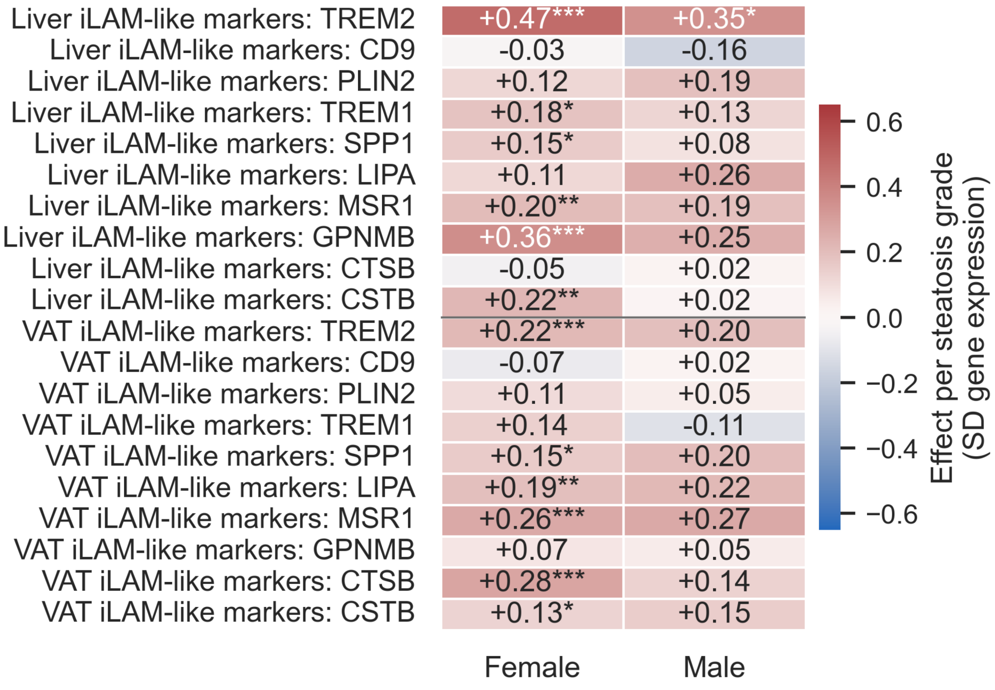

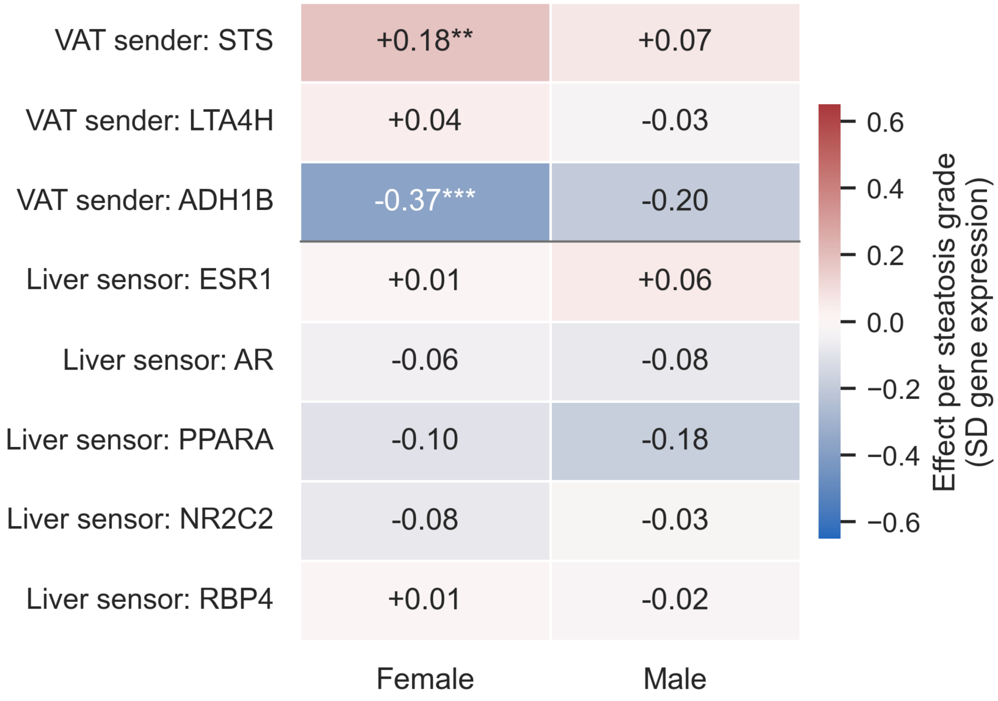

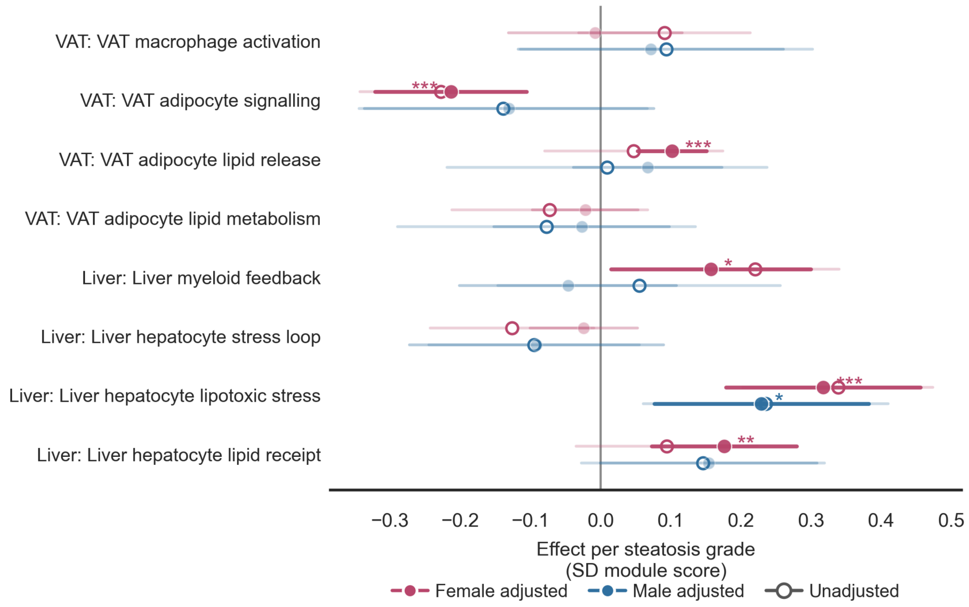

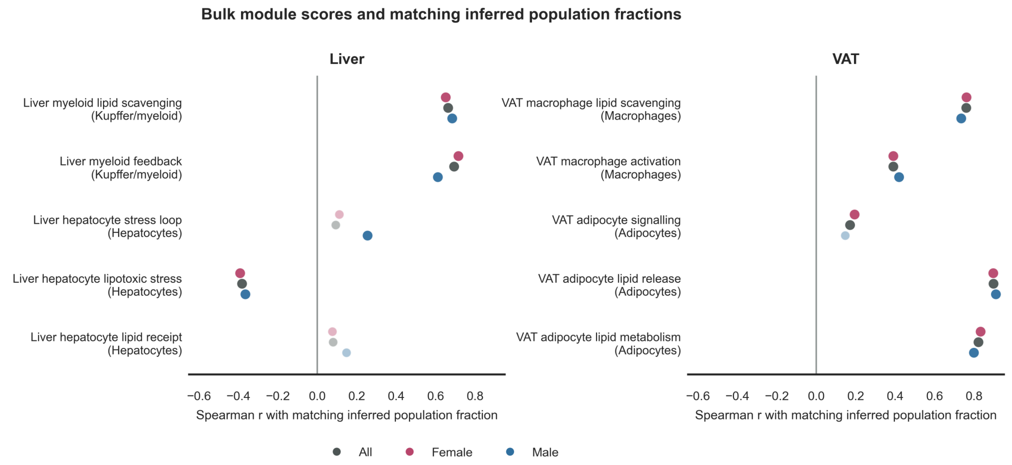

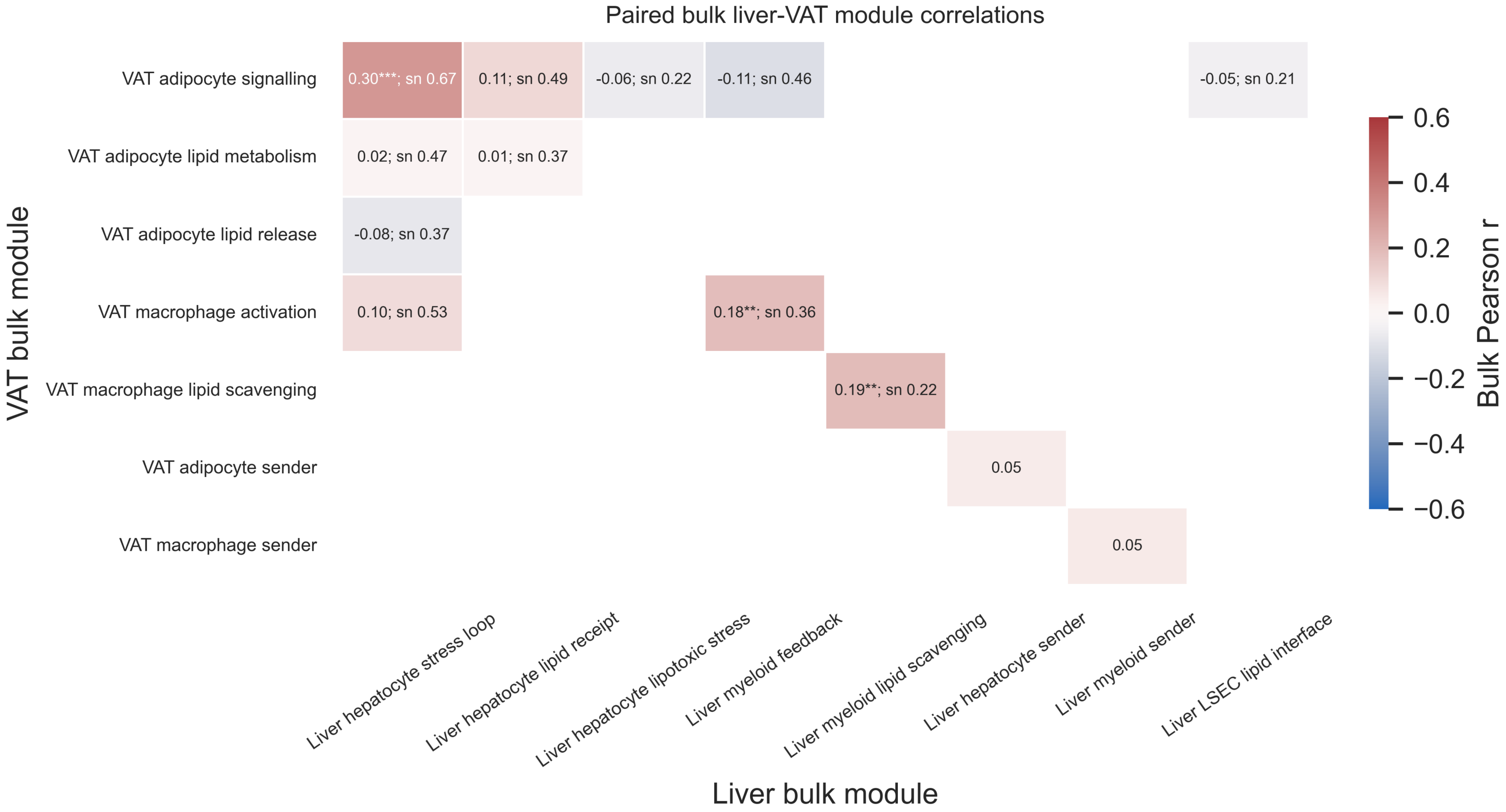

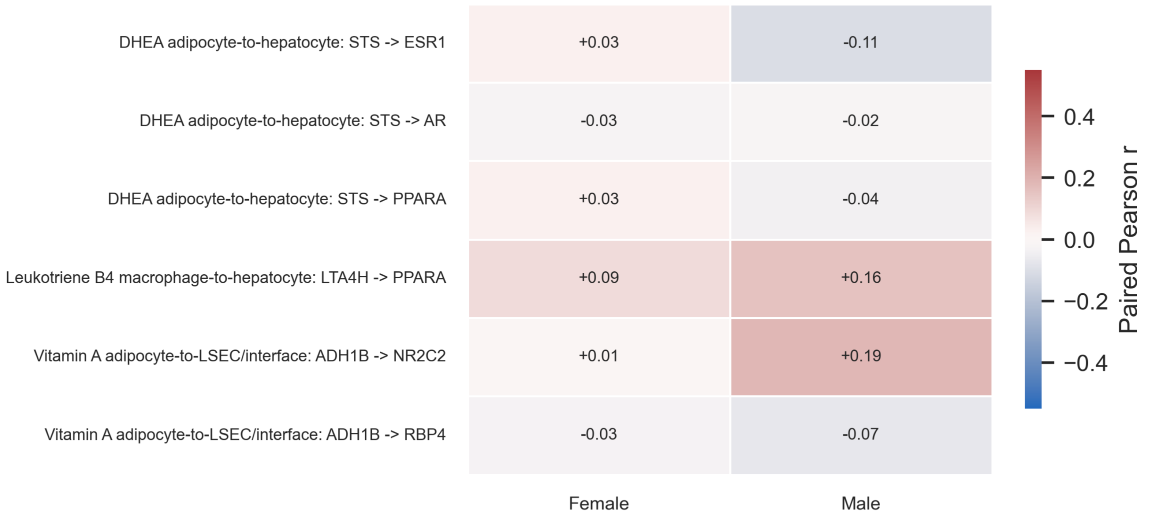

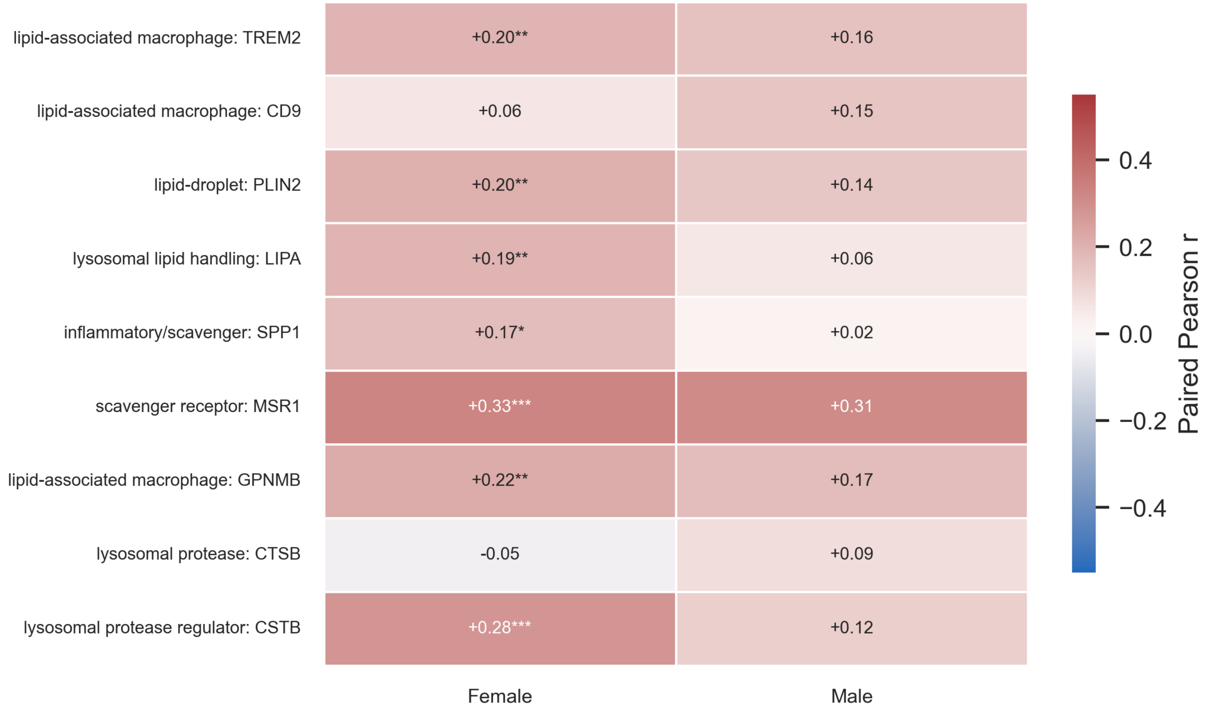
